# Multi-task spatial omics analytics for high-precision modeling of tissue architecture with STAX

**DOI:** 10.64898/2025.12.30.697018

**Authors:** Zhen-Hao Guo, De-Shuang Huang, Shihua Zhang

## Abstract

The rapid advancement of spatial omics technology has revolutionized biomedical research, unveiling unprecedented insights into the molecular and cellular architecture of biological systems. Despite this progress, a critical limitation arises from the widespread use of disparate analytical tools within a single workflow, leading to inconsistencies caused by heterogeneous preprocessing pipelines and extensive method-specific hyperparameter tuning. To this end, we aim to unify diverse analytical tasks within a multi-task learning platform, STAX, for diverse spatial omics types. STAX excels across numerous critical applications, including spatial domain identification, spatial slice integration, cohort-level spatial analysis, spatial spot completion, cell-gene co-embedding, expression profile denoising, and 3D spatial multi-slice simulation. Extensive evaluations demonstrate that STAX consistently delivers superior performance, robustness, and biologically meaningful interpretations, establishing it as an indispensable tool for spatial omics research. In short, STAX effectively addresses multiple analytical challenges in spatial omics, empowering researchers with a unified platform to accelerate biomedical discoveries and deepen understanding of complex biological systems.

## Introduction

Single-cell omics technology has revolutionized biological research by enabling the precise characterization of cellular states at finer resolution, facilitating comprehensive multi-omics analyses that span transcriptomics, epigenomics, proteomics, and beyond [1]. Unlike traditional bulk sequencing, which averages signals across cell populations, single-cell sequencing delivers high-resolution data with rich insights into cellular heterogeneity and function [2]. However, a critical limitation of this approach lies in the tissue dissociation step, which inherently disrupts the native spatial architecture of cells, resulting in the loss of crucial contextual information [3]. This spatial context is indispensable for deciphering the complex interplay within tissue microenvironments. For instance, the spatial distribution of tertiary lymphoid structures has been shown to strongly correlate with patient treatment outcomes and prognosis, underscoring the importance of preserving spatial information in biological studies [4, 5].

Spatial omics technology has emerged as a transformative tool that simultaneously captures different omics expression profiles and the precise spatial coordinates of cells within intact tissues [6, 7]. The groundbreaking potential of spatially resolved transcriptomics was formally recognized [8]. The field has witnessed rapid advancements, with the development of cutting-edge spatial transcriptomics platforms, including 10x Visium [9], Slide-seq [10], Stereo-seq [11], MERFISH [12], and STARmap [13]. Beyond transcriptomics, the spatial omics revolution is expanding to encompass diverse molecular layers, including spatial epigenomics (spatial ATAC–RNA-seq [14]), spatial translatomics [15], spatial proteomics (CODEX [16]), and spatial metabolomics (DESI-MS [17]). These technological innovations are collectively pushing the boundaries of spatially resolved biological insights, offering unprecedented opportunities to unravel the intricate molecular and cellular interactions within tissues [18].

The processing and analysis of spatial omics data present a wide variety of significant computational and methodological challenges. Key issues include the precise delineation of spatial domain boundaries within complex tissue contexts [19], technical variability arising from differences in sample preparation protocols and sequencing platforms [20], data sparsity due to spot gaps or tissue degradation, and the limited interpretability of analytical results. As spatial omics technologies continue to advance and their applications expand into three-dimensional tissue modeling, these challenges have become increasingly critical, demanding innovative computational solutions to unlock the full potential of spatial biology [21].

The challenges above arise from the trade-offs among multiple disciplines during technology development, yet they simultaneously present opportunities for advancing data modeling and computational methodologies [22]. For instance, SpaGCN pioneered the application of graph neural networks (GNNs) in the spatial domain identification task [23]. STAGATE substantially enhanced accuracy through self-attention mechanisms and extended its applicability to diverse scenarios [24]. In the batch-effect correction task, PASTE represents an early algorithmic solution for spatial transcriptomics alignment and integration [25], achieving three-dimensional reconstruction through probabilistic mapping based on gene expression and spatial position similarity. STAligner advanced this field by employing triplet loss to accurately identify similar structures across conditions, technologies, and developmental stages, effectively eliminating batch effects [26]. For the spatial resolution enhancement task, iStar utilizes a hierarchical vision transformer to extract information from histopathological images corresponding to sequencing slices, enabling spatial expression prediction [27]. STAGE eliminates the requirement for stained images [28], thereby expanding its applicability. SIMBA has demonstrated success in single-cell data analysis through the co-embedding of samples and features in a common space [29], enabling the depiction of cellular heterogeneity and the discovery of clustering-free markers. Yet, similar computational frameworks have not yet been developed for spatial omics applications.

More critically, the use of different computational tools for the same data analysis brings severe challenges. They primarily stem from inconsistencies in data preprocessing workflows and the extensive hyperparameter tuning required across different methods. Such variability impedes the comparability of results and limits the generalizability of findings. Therefore, there is an urgent need for a unified computational framework to address the above tasks comprehensively.

To this end, we develop STAX to address the challenges associated with multiple analytical tasks in a single framework. By consolidating these diverse analytics into a cohesive framework, STAX minimizes the need for disparate preprocessing pipelines and observably reduces the complexity of hyperparameter optimization. Through rigorous evaluation across a wide range of scenarios, we demonstrate that STAX consistently delivers efficient, high-performance results while ensuring stability. This positions STAX as a robust and reliable solution for navigating the complexities of spatial omics data analysis, ultimately advancing the field toward more conforming, reproducible, and interpretable biological and medical discoveries.

## Results

### Overview of STAX

STAX employs a graph attention network (GAT) encoder-decoder architecture, leveraging self-attention mechanisms to capture complex spatial context relationships (**Fig. 1a, Supplementary Fig. S1a** and **Methods**). For multi-slice integration, STAX utilizes the domain-specific batch normalization to keep the unique characteristics of each slice while ensuring data-wide consistency. To obtain gene embeddings and enhance gene-level analysis, STAX estimates the marginal probabilities of cell embeddings and decoder parameters, allowing for the computation of new gene embeddings that reflect both spatial and functional relationships. The reconstruction loss function can effectively denoise and impute the gene expression. For spot completion, STAX applies a masked loss function, which ignores simulated (fake) spots and focuses on reconstructing real data spots, enabling the expression to spread from the real spots to the fake spots through the physical edges. To ensure scalability, STAX implements subgraph sampling, enabling efficient mini-batch training applicable to spatial omics datasets of arbitrary size. By constructing inter-slice edges, STAX extends its capabilities to 3D spatial omics data analysis, facilitating the exploration of spatial patterns across multiple stacked spatial slices. Overall, STAX integrates various tasks into a unified framework through a carefully designed architecture and task-specific loss functions, while minimizing additional computational costs to ensure the model remains efficient. At the same time, its requirement for the format of the input data is relatively lenient and can easily accommodate diverse data types, further enhancing its applicability in the field of spatial omics biology (**Supplementary Fig. S1a**).

**Figure 1.**
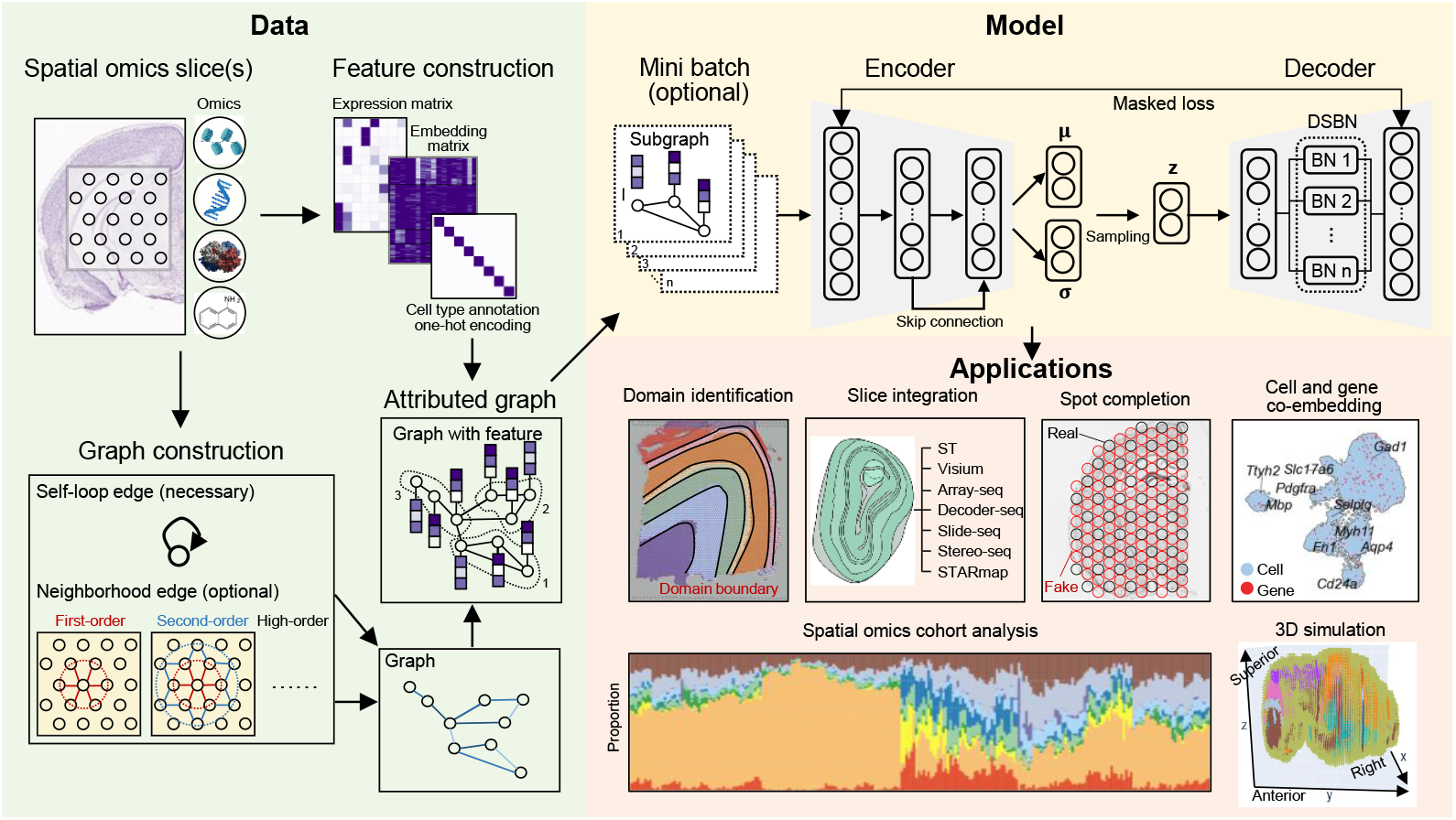
Overview of STAX. STAX begins with the construction of omics features (e.g., expression, cell type one-hot encoding, etc.) and neighbor graphs. Then, the attributed graphs are fed into a multi-task graph variational autoencoder. The cell embeddings generated by the encoder can be utilized for downstream tasks such as domain identification, slice integration, and cohort analysis. The decoder parameters are transformed to serve as gene embeddings. Additionally, STAX supports spot completion tasks through a masked reconstruction loss, which only calculates the error of the real spot. Furthermore, by constructing inter-slice edges, STAX extends all the above functionalities to 3D spatial multi-slice data analysis, enabling seamless simulation analysis across multiple stacked spatial slices.

### STAX outperforms competing methods in benchmark datasets for single-omics spatial domain identification

To quantitatively assess the spatial domain identification performance of STAX, we employed five single-omics benchmark datasets (**Supplementary Fig. S1b**): mouse brain dataset 1 (epigenomics) [14], human dorsolateral prefrontal cortex dataset (DLPFC, transcriptomics) [9], mouse brain dataset 2 (translatomics) [15], mouse spleen dataset (proteomics) [16], pig embryo dataset (metabolomics) [17]. Among these, the mouse brain dataset 1, DLPFC, and pig embryo datasets have a resolution of approximately 50µm, encompassing thousands of spots per dataset. The mouse spleen dataset (proteomics) comprises three slices, each containing approximately 80,000 cells, although it measures only 30 proteins. In contrast, the mouse brain dataset 2 includes 60,000 cells and quantifies over 5,000 translation genes. The above datasets collectively offer a diverse range of conditions. Each dataset exhibits unique characteristics, allowing us to simulate various real-world scenarios and comprehensively assess the efficacy of different algorithms under diverse conditions.

For evaluation metrics, the DLPFC and mouse spleen datasets, which include manually annotated domain labels, allow us to calculate the adjusted Rand Index (ARI) by comparing the predicted results with the manual annotations. For the remaining three datasets, we employ Moran’s I and Geary’s C to evaluate the spatial smoothness of the identified domains. Specifically, we quantify the spatial smoothness at the dimension level of the embeddings using Moran’s I and Geary’s C. We further extend this analysis to the predicted domain labels to evaluate their spatial coherence (**Methods**). Furthermore, we conduct a qualitative assessment by comparing the identified domains with corresponding stained images or annotations from the Allen Brain Atlas, providing an intuitive evaluation of the identified spatial domains.

For baseline comparisons, we evaluated several competing methods, including STAGATE [24], GraphST [30], SEDR [31], DeepST, PAST [32], MENDER [33], SpaceFlow [34], scNiche [35], INSTINCT [36], Louvain [37], and Leiden [38]. However, due to computational constraints such as out-of-memory errors during neighbor graph construction and GPU training, or sensitivity to hyperparameter tuning, we selectively applied appropriate baselines to each dataset based on their feasibility and performance.

For the mouse brain dataset 1, a spatial RNA-ATAC multi-omics dataset, we exclusively utilized the ATAC features for training both STAX and STAGATE (**Fig. 2a**). Our analysis revealed that STAX successfully identified nearly all structures with well-defined boundaries, including the Cerebral Cortex (CTX) layers 1-6, Anterior Cingulate Area (ACA), Genu of Corpus Callosum (CCG), Lateral Ventricle (VL), Caudoputamen (CP), Lateral Septal nucleus (LS), Lateral Preoptic Area (LPO), Medial Septal nucleus (MS), Nucleus Accumbens (ACB), Anterior Commissure Olfactory limb (ACO), Cortical subplate (CTXsp), and Piriform area (PIR). In contrast, STAGATE achieved comparable results but failed to detect LPO and CTXsp, and exhibited confusion between CTX1, ACA, and MS. Furthermore, we applied SpatialGlue, the state-of-the-art multi-omics integration algorithm, using both RNA and ATAC features as inputs. While SpatialGlue produced similar domain identification results, the boundaries were less smooth compared to those identified by STAX and STAGATE, as reflected in their Moran’s I and Geary’s C values (**Fig. 2a**). Notably, the absence of LPO and CTXsp in STAGATE’s results led to higher label Moran’s I and Geary’s C values compared to STAX. This observation suggests that Moran’s I and Geary’s C provide only a rough estimate of smoothness and can’t serve as definitive metrics (i.e., golden standard) for evaluating domain identification performance. Additionally, all methods identified a concordant result, i.e., they vertically divided the ACA into two distinct regions and conflated MS with PIR. These findings suggest that spatial domains, defined by structural anatomy and the actionable expression of chromatin, exhibit inconsistencies.

**Figure 2.**
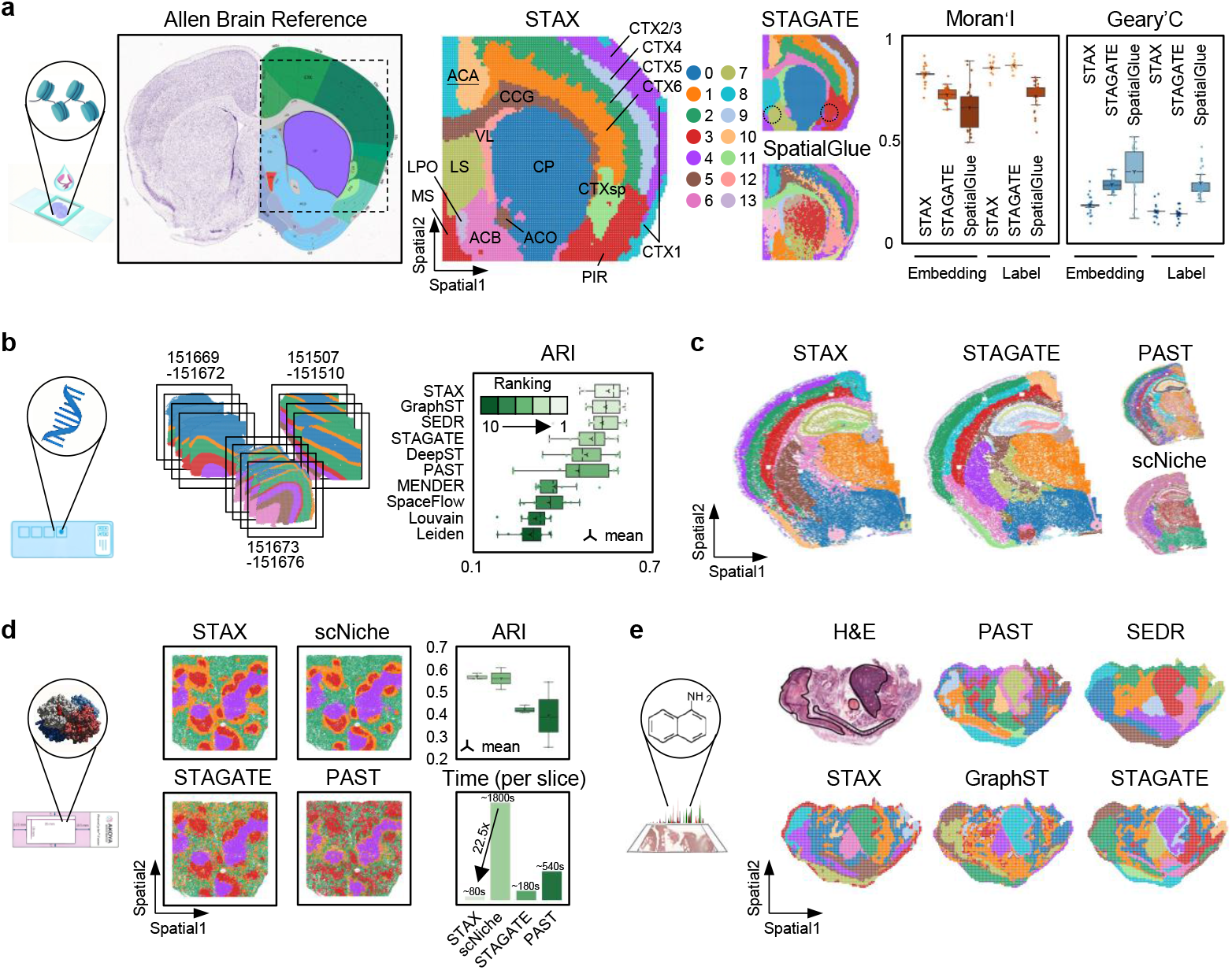
STAX outperforms the competing methods in benchmark datasets for single-omics spatial domain identification. **a**, The Allen brain reference annotation (left) and the results of STAX, STAGATE, and SpatialGlue are presented on the mouse brain dataset 1. Comparison of Moran’s I and Geary’s C metrics is reported on the right. **b**, The ARI performance of STAX and baseline methods on the 12 DLPFC slices. **c**, Spatial domains of STAX, scNiche, STAGATE, and PAST on the mouse brain dataset 2. **d**, Spatial domains and comparison of STAX, STAGATE, PAST, and scNiche on the BALBc1 slice of the mouse spleen dataset. **e**, Spatial domains of STAX, PAST, GraphST, SEDR, and STAGATE on the pig embryo dataset.

For the DLPFC dataset, a widely recognized benchmark for 10x Visium spatial transcriptomics, the dataset comprises 12 slices organized into three groups, with each group containing four slices (**Fig. 2b**). As described by Maynard et al. [9], each slice was manually annotated based on histologically stained images and marker genes, delineating cortical layers 1-6 as well as the white matter (WM). We evaluated all baseline algorithms on this dataset and found that STAX achieved the best performance, as measured by ARI (**Fig. 2b** and **Supplementary Fig. S2**).

For the mouse brain dataset 2, we conducted a comparative analysis using STAX, STAGATE, PAST, and scNiche (**Fig. 2c**). PAST, due to its early stopping strategy, failed to train adequately and was unable to accurately identify the structural regions. scNiche also demonstrated suboptimal performance, failing to achieve the desired outcomes in this context. In contrast, both STAX and STAGATE effectively identified the main brain structures.

For the mouse spleen dataset, a spatial proteomics dataset generated using CODEX technology, the data is manually annotated and primarily includes regions such as the B-zone, PALS (periarteriolar lymphoid sheaths), Marginal zone, and Red pulp (**Fig. 2d** and **Supplementary Fig. S3**). Each sample in this dataset comprises approximately 80,000 cells and 30 features. The substantial cell count poses serious computational challenges, particularly in constructing the cell network and conducting GPU-based training, both of which are highly memory-intensive. As a result, many existing methods that lack optimization for these steps either fail to execute or exhibit prohibitively slow performance. To ensure a rigorous comparison with STAX, we conducted fair evaluations against three methods: (i) scNiche, the most recent top method in this dataset; (ii) STAGATE (lightweight version), a computationally efficient alternative; and (iii) PAST, which represents, to the best of our knowledge, the first domain identification algorithm capable of graph mini-batching. As illustrated in **Fig. 2d** and **Supplementary Fig. S3**, STAX demonstrates consistent superiority across both clustering accuracy and computational efficiency, achieving a remarkable 20-fold speedup compared to scNiche. These results underscore STAX’s exceptional scalability and efficiency in handling large-scale spatial proteomics datasets.

For the pig embryo dataset, the sequencing spots exhibit a square structure, closely resembling the spatial arrangement observed in ST data. We evaluated STAX, STAGATE, GraphST, PAST, and SEDR on it (**Fig. 2e**), and nearly all methods successfully identified the primary embryonic structures, including the proencephalon, mesencephalon, heart, and liver, with clear delineation, showing the general applicability of STAX to spatial metabolomics data.

In short, STAX demonstrates remarkable improvements in performance compared to existing methods, establishing itself as a powerful tool for domain identification. Its ability to handle datasets of varying scales highlights its scalability. Furthermore, the flexibility of STAX is illustrated by its successful application across multiple omics types, including epigenomics, transcriptomics, and proteomics, as well as its compatibility with spatial technologies of different resolutions, such as 10x Visium and CODEX. This versatility positions STAX as a versatile tool for researchers seeking to uncover spatial patterns in complex biological systems.

### STAX accurately identifies spatial domains across multiple slices of different techniques, resolutions, and scales

Compared to analyzing a single spatial slice, multi-slice integration analysis offers a valuable opportunity to fully decipher different information from cross-platform spatial omics technologies such as Visium, Slide-seq, and STARmap. Each has unique strengths and limitations in resolution, throughput, and detection sensitivity. To comprehensively assess the integration capability of STAX in spatial transcriptomics, we collect: ST (100µm, 262 spots) [39], 10x Visium (55µm, 1,185 spots) [homepage], Array-seq (30µm, 7.7k spots) [40], Decoder-seq (15µm, 1,710 spots) [41], Slide-seq (10µm, 20k cells) [42], Stereo-seq (1µm, 19k cells) [43], and STARmap (single cell, 20k cells) [44] (**Supplementary Fig. S1b**). The MOB, with its well-defined and straightforward hierarchical structure, is widely utilized as a benchmark for evaluating the performance of sequencing technologies. These datasets span a broad spectrum of resolutions, scales, and gene numbers, simulating real-world experimental conditions and presenting significant challenges for integration algorithms.

We applied STAX to this diverse dataset and successfully identified eight distinct spatial domains across all samples (**Fig. 3a**), including olfactory nerve layer (ONL1, ONL2), glomerular layer (GL), mitral cell layer (MCL), granule cell layer (GCL), rostral migratory stream (RMS), granular layer of the accessory olfactory bulb (AOBgr), and accessory olfactory bulb (AOB). The UMAP visualization demonstrates that STAX effectively preserves the hierarchical organization of the MOB, from its inner to outer layers. The accuracy of these domains was further validated using established marker genes (**Fig. 3c, d**). Notably, AOBgr and AOB were exclusively identified in the Slide-seq and STARmap datasets, while the remaining structures were consistently detected across all datasets, indicating the precision of STAX. Furthermore, the absence of the RMS domain in the Decoder-seq data, as reported in the original study, confirms that STAX avoids over-integration. Of particular interest is the successful identification of substructures ONL1 and ONL2, as evidenced by the expression patterns of *Apod* and *Npy* in the heatmap. For comparative analysis, we employed STAligner and Harmony as baseline methods (**Fig. 3b** and **Supplementary Fig. S4a**). Although STAligner effectively integrated sequence-based ST data with others, it has difficulty correctly aligning the STARmap data, which is an image-based dataset. Harmony, on the other hand, failed in this scenario. STAX demonstrates competitive performance in handling integration tasks across different technologies.

**Figure 3.**
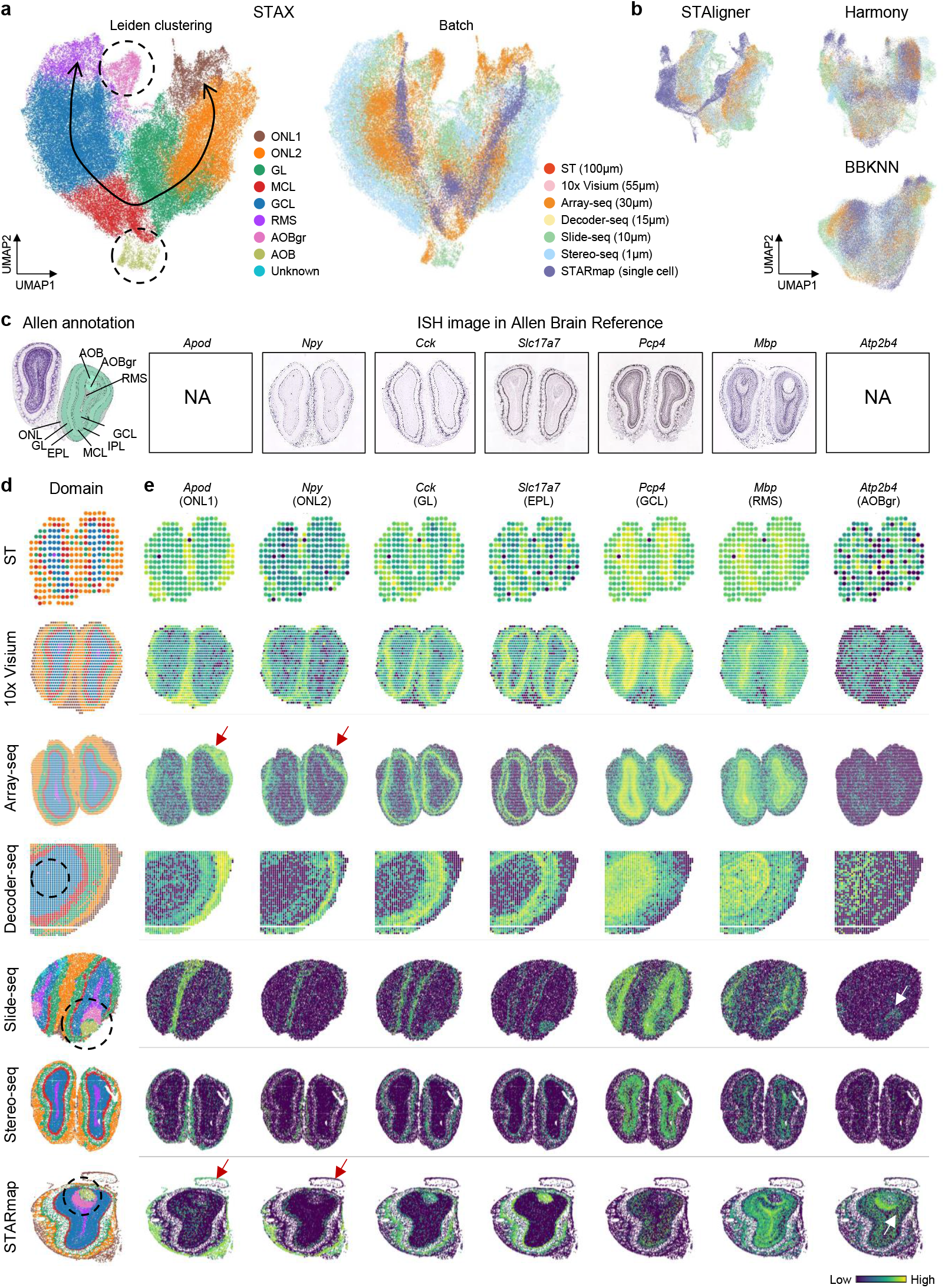
STAX accurately identifies spatial domains across seven mouse olfactory bulb (MOB) datasets generated by seven distinct technologies. **a**, UMAP visualization illustrating the integration results of STAX. **b**, UMAP visualization displaying the baseline integration results, including STAligner, Harmony, and BBKNN. **c**, The Allen reference annotation and *in situ* hybridization images of marker genes in the mouse olfactory bulb are shown. **d**, Expression patterns of domain marker genes across different datasets are presented.

Additionally, we benchmarked STAX against the competing methods on the spatial ATAC-seq dataset (mouse brain dataset 1). Specifically, we compared STAX with INSTINCT, a recently developed approach for spatial chromatin accessibility analysis. To ensure a fair comparison, we aligned the features of three ATAC-seq slices using INSTINCT’s preprocessing workflow. Our findings indicate that STAX generates smoother spatial domain boundaries while successfully identifying all cortical structures (**Supplementary Fig. S4b**). In contrast, INSTINCT produces coarser boundaries, although it identifies two additional subregions, i.e., the lateral preoptic area (LPO) and cortical subplate (CTXsp). Overall, STAX exhibits a competitive edge even when compared to specialized spatial ATAC-seq integration algorithms.

For spatial proteomics (mouse spleen dataset), we compared STAX with scNiche, the leading method for spatial proteomics data analytics, across three tissue slices. STAX consistently outperformed scNiche in terms of accuracy and computational efficiency, whether processing single or multiple slices (**Supplementary Fig. S5a, b**). We also conducted extensive hyperparameter tuning and ablation studies to provide users with practical guidance for selecting optimal configurations for specific applications (**Supplementary Fig. S5c-h**). These experiments further validate STAX’s robustness and adaptability across diverse analytical scenarios.

### STAX enables integration of spatial omics cohort across hundreds of individuals

With the continuous reduction in the cost of spatial technologies, spatial cohort analyses have become increasingly prevalent. The emergence of large-scale spatial omics cohorts comprising hundreds of samples has imposed new demands on model scalability. Here, we apply STAX to integrate over 100 spatial tissue slices for characterizing domain functions by identifying marker genes and cell-type composition, and to associate spatial domains with phenotypes. We demonstrate the applicability of STAX using two large-scale cohorts: a CosMx spatial transcriptomics cohort of ovarian cancer with 100 slices and ~0.5 million cells [45] and a CODEX spatial proteomics cohort of hepatocellular carcinoma with 238 slices and ~2.6 million cells [46].

For the first spatial transcriptomics cohort, we considered two feature types as inputs for STAX (**Methods**), i.e., gene expression profiles and one-hot encoded cell type annotations. Drawing upon a previous stability metric [47], we observed that the domains identified using one-hot encoded cell type annotations exhibited superior performance (**Fig. S6a**). Furthermore, compared to the 960-dimensional gene expression feature, the 13-dimensional one-hot encoded cell type annotations demonstrated significantly faster training convergence and reduced computational time (**Fig. S6b**). Finally, we employed the latter as features to integrate the cohort and identified 12 functionally distinct domains (**Fig. 4a, b**), depicting the disease heterogeneity in this cohort.

**Figure 4.**
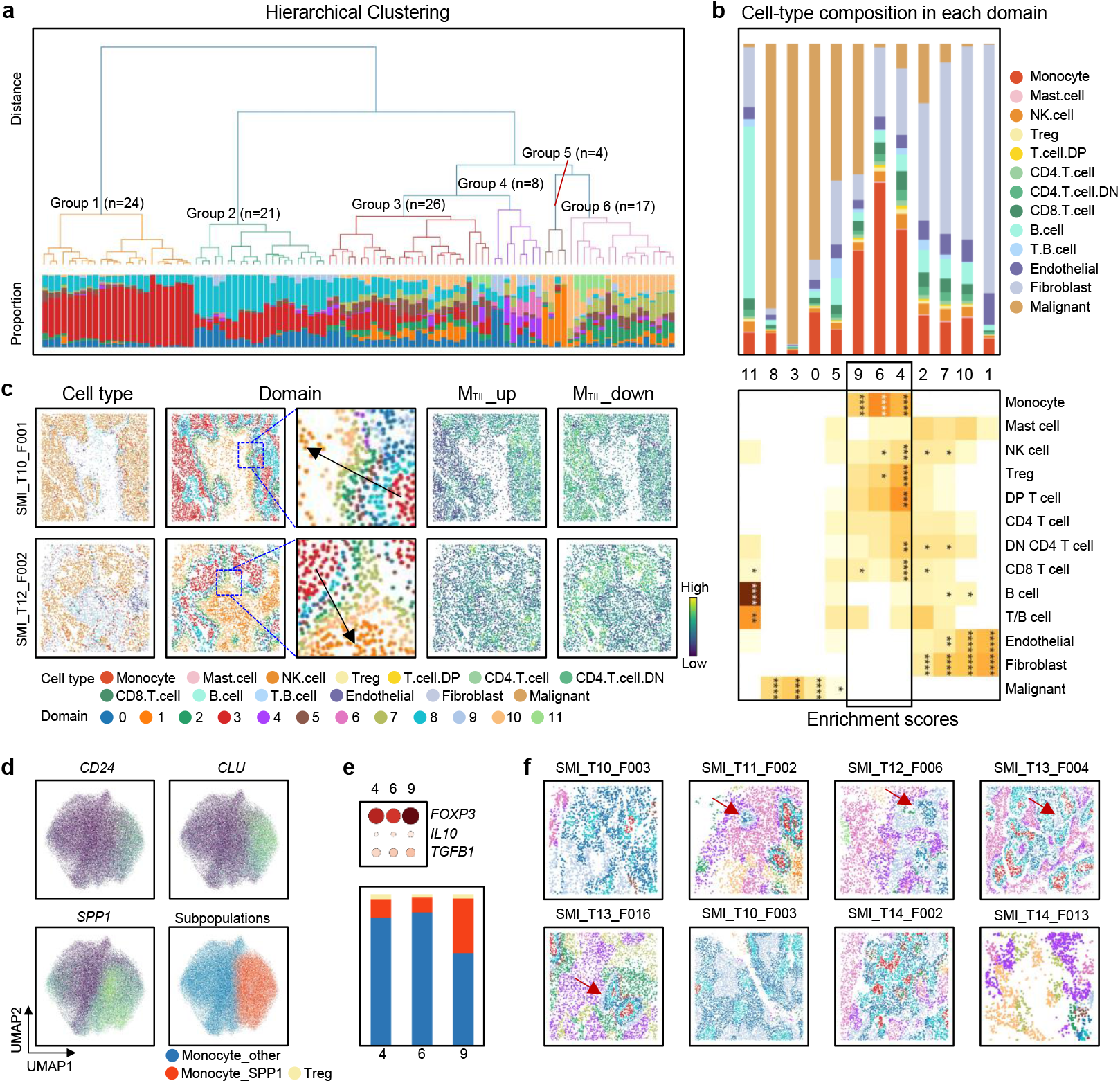
STAX enables integration of a spatial transcriptomics cohort across hundreds of individuals. **a**, Hierarchical clustering of 100 slices based on the proportion of spatial domains identified by STAX, with different colors indicating domains. **b**, Cell-type composition of each domain (up), as well as the enrichment scores of the cell types identified by scNiche in the different domains (down). **c**, Distribution of the cell type, domain, and expression of MTIL_up and MTIL_down in the SMI_T10_F001 and SMI_T12_F002 slices. **d**, Expression of *CD24, CLU* and *SPP1* in monocytes. The division of Monocyte_other and Monocyte_SPP1 in Monocyte. **e**, Dotplot of *FOXP3, IL10*, and *TGFB1* expressed by Treg in domains 4, 6, and 9 (up). The proportion of three cell types, including Monocyte_other, Monocyte_SPP1, and Treg, in domains 4, 6, and 9 (down). **f**, Spatial domains of eight slices that contain domains 4, 6, 9.

We identified distinct molecular signatures characterizing each spatial domain through differential expression analysis (**Supplementary Fig. S6c, d**). For example, domains 0, 3, and 8 exhibited dual expression of epithelial markers (*KRT* family) and antigen presentation genes (*CD74, HLA-DRA*), strongly suggesting these regions represent malignant cells with immune-evasive properties. The monocyte-enriched domains 4, 6, and 9 illustrated distinct functions, as evidenced by co-expression of macrophage markers (*CD68, LYZ*) and MHC class II molecules (*HLA-DR/DQ*/*DP*). Domains 5 and 11 were defined by plasma cell markers (*IGKC*/*IGHG1*/*IGHG2, MZB1, JCHAIN*), reflecting their specialized role in humoral immunity through immunoglobulin (IgG/IgA) production. Conversely, domains 1, 2, 7, and 10 showed a distinct abundance of Extracellular Matrix (ECM) components (like the COL family proteins), being consistent with Cancer-Associated Fibroblasts (CAFs) actively remodeling the tissue structure around the tumor.

Furthermore, we validated these findings through a histogram plot of cell type proportions and quantitative analysis via scNiche-derived enrichment scores (**Fig. 4b**). The results clearly showed a distinct cell distribution across the regions: Malignant cells were the main population in domains 0, 3, and 8. Immune cells were highly present in domains 4, 6, and 9. Stromal cells (such as endothelial cells and fibroblasts—the supportive tissue cells) dominated domains 1, 2, 7, and 10. Additionally, the spatial distribution in samples SMI_T10_F001 and SMI_T12_F002 revealed functionally significant topological patterns, with domain 3 occupying the tumor cores while domains 0 and 8 localized to the invasive fronts (**Fig. 4c, left**). Moreover, tumor-infiltrating lymphocytes (TILs) are a class of immune cells that migrate from the bloodstream to the interior of Tumor tissue, and high levels of TILs (MTIL_up) are generally associated with better survival rates. Domain 3 expressed more MTIL_down, while domains 0 and 8 expressed more MTIL_up (**Fig. 4c, right**). We also identified domain-specific enrichment patterns with prognostic implications. Specifically, domains 3 and 8 were markedly enriched in untreated patients, and domain 3 was significantly enriched in dead samples. These findings collectively demonstrate how spatial domain organization reflects both biological function and clinical behavior in the tumor ecosystem (**Supplementary Fig. S6e**).

Finally, we divide monocytes into two functional subtypes, i.e., Monocyte_SPP1 and Monocyte_other (**Fig. 4d**). The histogram plot revealed that the Monocyte_SPP1 subtype exhibited more in domain 9 (**Fig. 4e**). However, although the proportional abundance of regulatory T cells (Tregs) in this region showed no significant change, the expression levels of their key functional markers (including *FOXP3, IL10*, and *TGFB1*) were markedly upregulated. This finding was corroborated by spatial distribution patterns that domain 9 maintained close spatial proximity to the tumor region (domains 0, 3, 8), further supporting the biological basis for the formation of this immunosuppressive microenvironment (**Fig. 4f**). The results of STAX for each slice can be seen in **Supplementary Fig. S7-S8**.

Next, we applied STAX to the spatial proteomics cohort [46]. Similar to the spatial transcriptomics cohort study, we observed that the domains identified using one-hot encoded cell type features exhibited superior performance and less time than protein expression (**Supplementary Fig. S9a, b**). Finally, we identified 14 functionally distinct domains.

Given the limited number of protein features (n=36) in this dataset, we are unable to perform comprehensive molecular-level differential analysis or functional enrichment. Instead, we directly assessed the cellular composition of each structural domain by analyzing (1) the proportion of cell types within each domain and (2) the enrichment patterns of cell types across domains to infer their potential biological functions (**Supplementary Fig. S9c**). Domains 2, 5, 6, and 12 exhibited high immune activity, suggesting associations with antitumor immunity or inflammatory responses. Domains 3, 4, and 11 were predominantly composed of stromal cells (e.g., fibroblasts) and endothelial cells, likely contributing to tissue integrity and angiogenesis. Domain 11 showed enrichment in cholangiocytes and macrophages, potentially implicating roles in biliary repair or inflammation. Domains 0, 1, and 9 contained mixed populations of tumor cells, immune cells, and stromal components, possibly representing transitional zones for tumor-immune interactions. Domains 7, 8, and 10 were primarily tumor cell-enriched, reflecting core tumor microenvironment regions. Notably, domain 10 was more frequently detected in deceased patients compared to domains 7 and 8 (**Supplementary Fig. S9d**). Further functional annotation using established protein markers confirmed its association with aggressive tumor phenotypes (**Supplementary Fig. S9e**). The results of STAX for each slice can be seen in **Supplementary Fig. S10-S15**.

Overall, despite the considerable complexity of tumor microenvironment heterogeneity across multiple samples, STAX effectively integrates hundreds of spatial omics slices for precisely identifying both shared and unique structural domains. By providing distinct molecular, cellular, and clinical characterization of each domain’s composition and function, STAX enables the construction of a comprehensive cohort atlas that systematically maps relationships between cells, structural domains, and clinical phenotypes.

### STAX achieves spot completion while deciphering the complex tumor microenvironment

To overcome the inherent resolution limitations of Visium, 10x Genomics developed Xenium, an innovative platform enabling single-cell resolution analysis. Recently, 10x Genomics applied these complementary technologies to breast cancer tissue, characterized by significant heterogeneity, thereby offering unique opportunities for comparative evaluation of their analytical capabilities (**Fig. 5a**). Following alignment of Xenium and Visium datasets using Spateo [48], we performed an integrated comparative analysis of both datasets through STAX.

**Figure 5.**
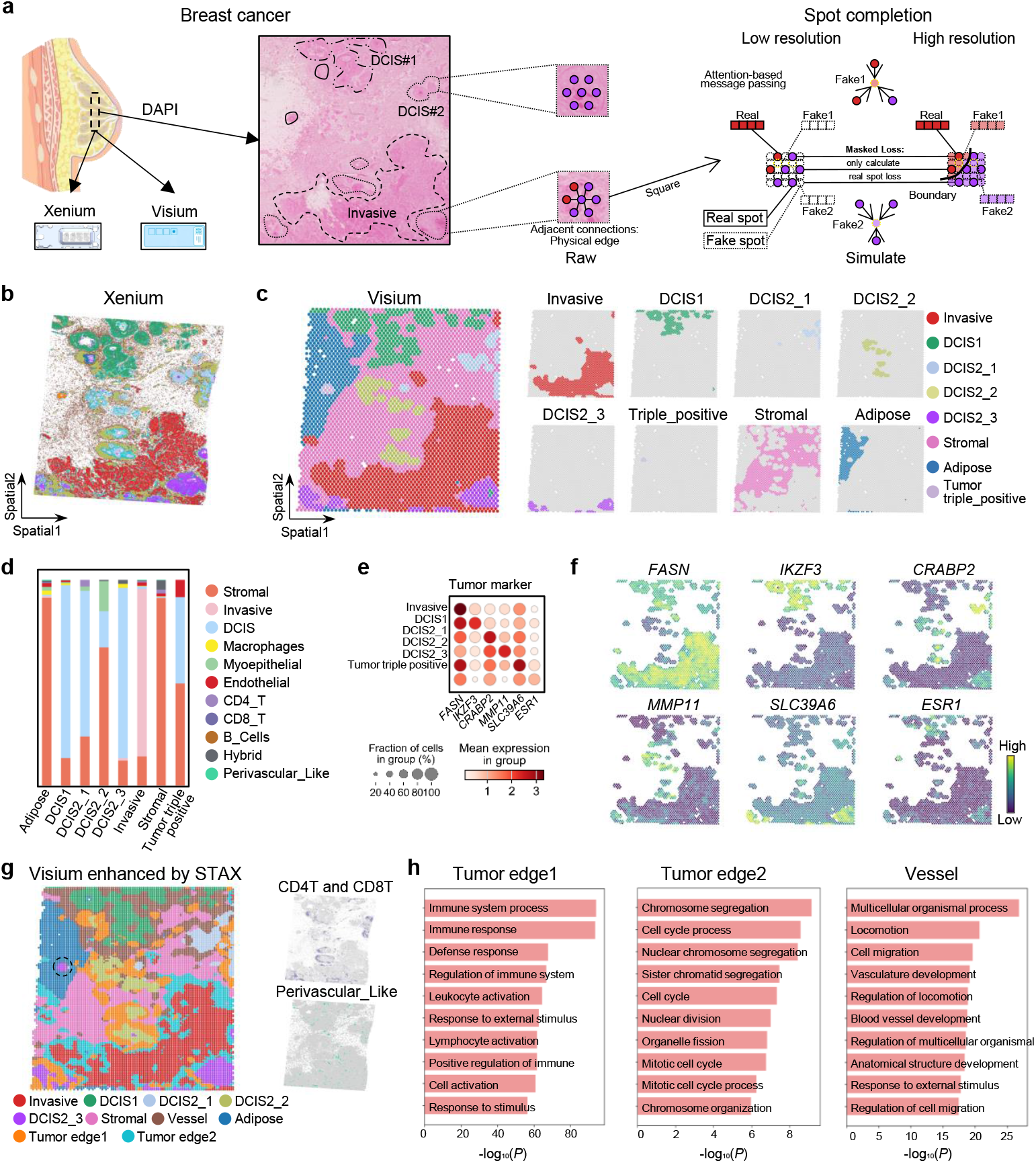
Application to integrating breast cancer data profiled by Xenium and Visium platforms illustrates that STAX can achieve spot completion for deciphering the tumor microenvironment. **a**, Schematic diagram of the breast cancer data and spot completion performed by STAX. **b** and **c**, Spatial domains identified on them by STAX, respectively. **d**, Cell proportions in different domains identified on the Visium dataset by STAX. **e**, Dot plot of six tumor markers, presenting on the Visium dataset but not the Xenium one. **f**, Heatmap of six tumor markers, presenting on the Visium dataset but not the Xenium one. **g**, Spatial domains identified by STAX after spot completion (left), distribution of CD4T/CD8T, and perivascular-like cells on the Xenioum dataset (right). h, GO enrichment analysis of the differentially expressed genes (DEGs) for the tumor_edge 1 (525 genes), tumor_edge 2 (262 genes), and vessel (321 genes) domains. DEGs for each domain were identified using a two-sided t-test by comparing the cells to all remaining ones on the imputed data. Genes with a p-value < 0.05 and log_2_ fold change ≥ 0.5 were considered as DEGs.

We applied STAX on the Xenium dataset, delineating nine spatial clusters (**Supplementary Fig. S16a**). Through differential expression analysis, we identified potential cluster-specific marker genes (**Supplementary Fig. S16b-d**), including *FASN* for the Invasive domain [49], and *CEACAM6* for the DCIS1 domain [50]. Although the original study classified DCIS into two subtypes, our results revealed further heterogeneity within DCIS2, which we subdivided into DCIS2_1/2 (characterized by differential expression of *KRT* family genes) and DCIS2_3 (marked by TENT5C expression). Furthermore, we identified a rare triple-positive breast cancer region exhibiting high co-expression of *ERBB2, ESR1*, and *PGR* (**Supplementary Fig. S17a**). We also analyzed the cell type composition of each cluster (domain) (**Supplementary Fig. S17b**). Finally, we defined them as nine distinct spatial domains, including invasive, ductal carcinoma *in situ* (DCIS), triple-positive tumor, myoepithelial, and tumor_edge regions (**Fig. 5b, Supplementary Fig. S17c**).

We applied STAX to the Visium dataset, delineating ten spatial clusters (**Supplementary Fig. S18a, b**). Despite analyzing the same tissue location, STAX enabled the identification of additional spatial domains in Visium, including invasive, DCIS, triple-positive tumor, myoepithelial, stromal, and adipose regions (**Fig. 5c, d**). We performed differential analysis to obtain marker genes for each cluster (**Supplementary Fig. S18c**). Compared to the Xenium analysis, we achieved further refinement of DCIS subtypes in the Visium data, distinguishing DCIS2_1 from DCIS2_2. DCIS2_1 exhibited high expression of *CPB1* and CRABP2, while DCIS2_2 was characterized by elevated expression of *KRT* family genes. DCIS2_3 showed predominant expression of *TFF* family genes (**Supplementary Fig. S19a, b**). Many of these differentially expressed genes, including members of the *KRT* [51] and *TFF* family [52], have established associations with cancer progression and prognosis, further underscoring the extensive heterogeneity within breast tumor cells. Furthermore, our analysis revealed potential marker genes in the Visium data that were not detected in the Xenium one (**Fig. 5e, f**), including *IZKF3* [53], *CRABP2* [54], *MMP11* [55], and *SLC39A6* [56]. Of particular interest was the identification of an adipose domain in the Visium data that was absent in the Xenium one. This discrepancy may be attributed to insufficient cell capture at the corresponding tissue location in the Xenium data, compounded by the lack of definitive adipose marker genes, which likely hindered the identification of a distinct adipose structure in it (**Supplementary Fig. S20a, b**).

In summary, the above results highlight Visium’s strengths in comprehensive gene coverage. However, its spatial resolution limitations, particularly the inability to measure gene expression in inter-spot regions, may obscure some structural details. To fully use the Visium data, we implemented STAX to enhance spatial resolution through spot complementation (**Fig. 5a**). Our approach involved three key steps: First, we generated pseudo-spots in the interstitial regions between existing spots. Second, we constructed a neighborhood graph encompassing both real and pseudo-spots, incorporating a neighbor proximity network to provide complementary topological information (**Fig. 5a**). Additionally, our training process exclusively considered the reconstruction loss for real spots, with the trained model subsequently generating expression profiles for fake spots.

This approach enabled us to expand the Visium dataset from the original 4,000 spots to approximately 10,000 (100×100) spots. The enhanced dataset revealed structural domains roughly consistent with the original analysis, including invasive, DCIS, stromal, and adipose regions, while additionally identifying three new distinct domains: tumor_edge1, tumor_edge2, and vessel (**Fig. 5g, left** and **Supplementary Fig. S21a-d**). Each domain exhibited unique biological characteristics. Specifically, the tumor_edge1 region was identified as an immune-related domain. GO enrichment analysis revealed significant associations with immune system processes and immune responses (**Fig. 5h**).

This finding may be attributed to the relatively benign surrounding ductal carcinoma. In contrast, the tumor_edge2 domain encompasses numerous cell cycle and nuclear division processes (**Fig. 5h**). This suggests that tumor cells in this region are highly active, which complements the characteristics of the adjacent infiltrative tumor domains. The vessel region was associated with tissue organization, specifically ductal formation, showing enrichment for multicellular organismal processes and blood vessel morphogenesis (**Fig. 5h**). We also observed the distribution of CD4T/CD8T, and perivascular_like cells in the Xenium dataset (**Fig. 5g, right**), which appears to align with the spatial locations of the tumor_edge1 and tumor_edge2 domains identified by STAX.

Moreover, we conducted a benchmark comparison using the 12-slice DLPFC dataset to evaluate the performance of STAX against STAGE [28] (**Supplementary Fig. S22a**). In this evaluation, we randomly removed 50% of the spots and utilized the remaining 50% for training. Based on a previous study indicating the limited informational value of DLPFC staining images [28], we did not compare STAX with some image-based prediction methods such as iStar [27] or OmiCLIP [57]. The computational efficiency comparison revealed a substantial advantage for STAX, with an average processing time of 20 seconds per slice compared to STAGE’s 1200 seconds per slice (**Supplementary Fig. S22b**). For performance assessment, we employed spot completion slices by STAX and STAGE as inputs to STAX and STAGATE, respectively, for quantitative comparison in spatial domain identification tasks. The results demonstrated superior performance of STAX-generated data across both evaluation metrics, indicating enhanced recovery capabilities (**Supplementary Fig. S22c**). To further validate our findings, we conducted qualitative analysis by visualizing layer-specific marker genes (**Supplementary Fig. S22d, e** and **Supplementary Figs. S23** and **S24**). Comparative analysis revealed consistent outperformance of STAX over STAGE across nearly all tissue slices, further substantiating the competitive ability of STAX.

### STAX pinpointed cell type, domain type, and disease-related specific genes in cell and gene co-embedding

STAX can achieve the co-embedding of cells and genes, offering an intuitive approach to decipher biologically pivotal clusters with cluster-relevant biomarkers. To demonstrate this, we applied it to a visual cortex dataset consisting of 12 tissue slices generated using MERFISH technology [12]. The dataset initially included annotations for cell type labels and their corresponding marker genes, as documented in the literature (**Supplementary Fig. S25a**). Recently, Yuan et al. [19] re-annotated the spatial domain labels and corresponding marker genes for five of these slices, providing a more refined and reliable reference. Marker genes of the medial preoptic area (MPA) and paraventricular hypothalamic nucleus (PVH) are not provided by them. Consequently, the annotations of cells, spatial domains, and marker genes in these five slices serve as a gold standard for validating the accuracy of cell and gene co-embedding.

To obtain cell-level embedding, we input cell features and a neighbor graph with the radius parameter (rad) set to 0 (i.e., containing only self-loops) into STAX to generate the co-embedding of cells and genes. Under setting 1, where the neighbor graph lacks neighbor edges with only self-loop edges, the graph autoencoder naturally simplifies to a standard autoencoder. The resulting UMAP visualization revealed a strong concordance between manually annotated cell-type labels and their corresponding marker genes (**Supplementary Fig. S25b**). Specifically, the co-embedding successfully captured the expression patterns of key marker genes across distinct cell types, including inhibitory neurons (*Gad1*), excitatory neurons (*Slc17a6*), mature oligodendrocytes (*Thyh2, Mbp*), immature oligodendrocytes (*Pdgfra*), astrocytes (*Aqp4*), microglia (*Selplg*), ependymal cells (*Cd24a*), endothelial cells (*Fn1*), pericytes, and mural cells (*Myh11*) (**Supplementary Fig. S25c**). The alignment between annotated cell types and marker genes substantiates the effectiveness of STAX in capturing biologically meaningful patterns and revealing subtle cell-gene relationships within complex tissue architectures.

To obtain domain-level embedding, we extended our analysis by inputting cell features and a neighbor graph with the radius parameter (rad) set to 50 into STAX to generate the co-embedding of cells and genes. This setting 2 allowed us to incorporate spatial context information into the model, enabling cell-gene relationships in domain-resolution representation. UMAP visualization revealed a strong concordance between manually annotated spatial domain labels and their corresponding marker genes. Specifically, the spatial domains including bed nuclei of the strata terminalis (BST, *Sytl4*), third ventricle (V3, *Cd24a*), periventricular hypothalamic nucleus (PV, *Slc18a2*), paraventricular nucleus of the thalamus (PVT, *Necab1*), medial preoptic nucleus (MPN, *Nts*), and columns of the fornix (fx, *Mbp*) were accurately identified and aligned with their respective marker genes (**Supplementary Fig. S25e-g**).

To further explore the cell-gene relationships within the co-embedding space, we constructed a network of cells and genes in the co-embedding space with settings 1 and 2, respectively. We hypothesize that nodes with higher clustering coefficients play more significant roles in the network and are more likely to serve as marker genes for specific cell types or spatial domains. Analysis of this network revealed that the clustering coefficient of genes exhibited a strong correlation with four key metrics proposed by SIMBA [29], i.e., Max, Std, Gini, and Entropy (**Supplementary Fig. S25d, h**). These four metrics, which quantify the variability and specificity of gene expression, are widely used to assess the importance of genes in single cell analysis. This observation aligns with the notion that genes with higher connectivity in the co-embedding space are functionally important. By leveraging these insights, STAX provides a novel framework for identifying candidate marker genes and elucidating their roles in tissue architecture and function. Moreover, we refined the cell-gene co-embedding network by removing the cells and retaining only the genes to explore gene-gene relationships. This pure gene-gene association network revealed potential regulatory or co-expression relationships, e.g., the edges connecting *Esr1* and *Pgr* (**Supplementary Fig. S26a, b**). This finding aligns with previous literature, which has documented important co-expression of estrogen receptor alpha (*Esr1*) and progesterone receptor (*Pgr*) with sets of steroid target genes identified in specific brain regions [58].

We also applied STAX to the Slide-seq dataset derived from the Alzheimer’s disease (AD) and normal mouse brain tissues (**Fig. 6a**). We utilized STAX to integrate data from both diseased and healthy samples and successfully identified shared structural domains across the two slices, including domain 4 (CA1), domain 5, and domain 9 (DG) (**Fig. 6a**), which align with findings from previous studies [26]. Building on the insights gained from cell and gene co-embedding, we proceeded to co-embed cells and genes within this dataset (**Supplementary Fig. S27a, b**). Notably, genes associated with DG, CA1, and CA3, such as *Prox1, Itpka*, and *Hs3st4*, were accurately and logically positioned within the co-embedded UMAP visualization (**Fig. 6b** and **Supplementary Fig. S27c**). This rational allocation underscores the ability of STAX to capture biologically relevant cell-gene relationships in complex tissue architectures.

**Figure 6.**
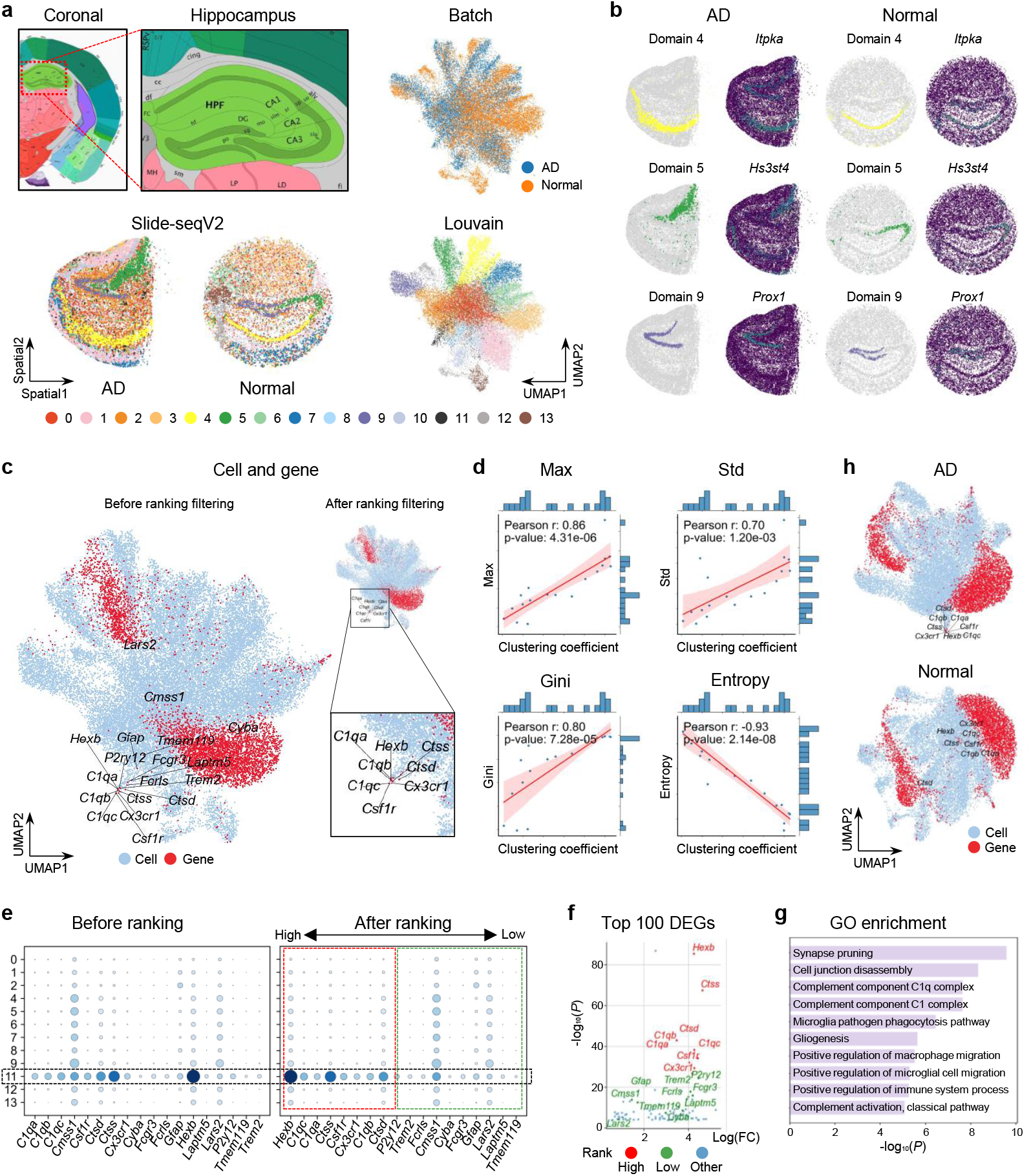
Application to integrating Alzheimer’s disease (AD) and healthy tissue slices demonstrates that STAX can pinpoint cell type, domain type, and disease-related genes in cell and gene co-embedding. **a**, Integration of the two slices for deciphering the aligned spatial domains and achieving batch effect removal by STAX. **b**, Illustration of three aligned spatial domains and their marker genes by STAX. **c**, Cell and gene co-embedding of the two slices, and genes are illustrated, with an amplified view of genes in domain 11 (AD-related domain). **d**, Correlations between the clustering coefficient in the constructed cell-gene networks and metrics such as MAX, Std, Gini, and Entropy. **e**, Dot plot of gene expression in domain 11 across all domains before and after ranking by clustering coefficient. **f**, Volcano plot highlighting differential genes in domain 11, where red, green, and blue represent high-ranking, low-ranking, and non-domain 11 marker genes identified by STAX, respectively. **g**, GO enrichment results of 18 genes for domain 11. **h**, Cell and gene co-embedding of the AD and healthy slices, as well as their genes, respectively.

Domain 11, as illustrated in the UMAP plot, was uniquely present in the AD slice and contained a distinct set of genes (**Fig. 6c**). By sorting genes based on their clustering coefficients and filtering out lower-ranked ones, we retained several genes of interest (**Fig. 6d, e**), such as *Hexb* [59], *Ctss* [60], *C1qc, Csf1r* [61], *C1qa* [62], *Cx3cr1* [63], *C1qb*, and *Ctsd*. Notably, some of these genes have been previously validated as being associated with Alzheimer’s disease. To further validate our findings, we conducted differential expression analysis with a volcano plot of marker genes specific to domain 11 and revealing that highly ranked genes exhibited statistically significance compared to lower-ranked ones, thereby indirectly corroborating the efficacy of our filtering strategy (**Fig. 6f**). Additionally, GO enrichment analysis confirmed the functional relevance of domain 11, highlighting its potential role in biological processes and pathways associated with Alzheimer’s disease (**Fig. 6g**). In addition, we performed co-embedding genes wth AD and healthy samples, respectively. In the UMAP representation of AD and gene co-embedding, AD-related genes were found to cluster closely around domain 11. In contrast, in the UMAP of healthy samples and gene co-embeddings, AD-related genes were dispersed. These results demonstrate that the cell-gene co-embedding generated by STAX is condition-specific and exhibits high accuracy under different experimental conditions (**Fig. 6h**).

In addition to the above applications in spatial transcriptomics, we also applied STAX’s spot completion and co-embedding functions to spatial proteomics and metabolomics datasets **(Supplementary Figs. S28-30)**, demonstrating its applicability to diverse spatial omics. In summary, this cell and gene co-embedding way provides an intuitive approach for validating and interpreting model predictions. By integrating gene expression data with spatial context, co-embedding helps elucidate the biological relevance of identified domains and uncovers potential new key genes that may serve as regulators of spatial organization. Thus, STAX enriches our understanding of biologically significant clusters (cell types or domains).

### STAX generates a high-resolution 3D map of the mouse embryo brain and simulates coronal, sagittal, transverse and any angle slices

Reconstructing three-dimensional (3D) organ structures pose a novel challenge for spatial omics technologies. Here, we applied STAX to a dataset of mouse embryonic brain generated using MAGIC-seq [64], demonstrating the effectiveness of all its functions in a 3D multi-slice dataset. To construct a comprehensive 3D representation, we first built a 3D neighbor graph based on the spatial positions of spots across all slices. This graph incorporates edges not only within individual slices but also between adjacent slices. Using STAX, we successfully identified ten major brain structures, including the central subpallium (CSPall), dorsal pallium (DPall), medial pallium (MPall), diencephalon, midbrain, inter and hindbrain, cerebellum, meninges, olfactory areas, and ventricular systems (**Fig. 7a, b** and **Supplementary Fig. S31a, b**). Since the cerebellum and midbrain are nearly absent in slice 296, we used slice 585 for detailed analysis. In the posterior region of slice 585, we discovered previously unannotated structures, such as the cerebellum and ventricular systems, which were validated by marker genes *Sox9* and *Pcp2* (**Fig. 7c**). Notably, *Pcp2* is a well-established marker gene for the Purkinje cell layer [65]. The spatial location of the slice further allowed us to prove the position of the cerebellum (**Fig. 7a**).

**Figure 7.**
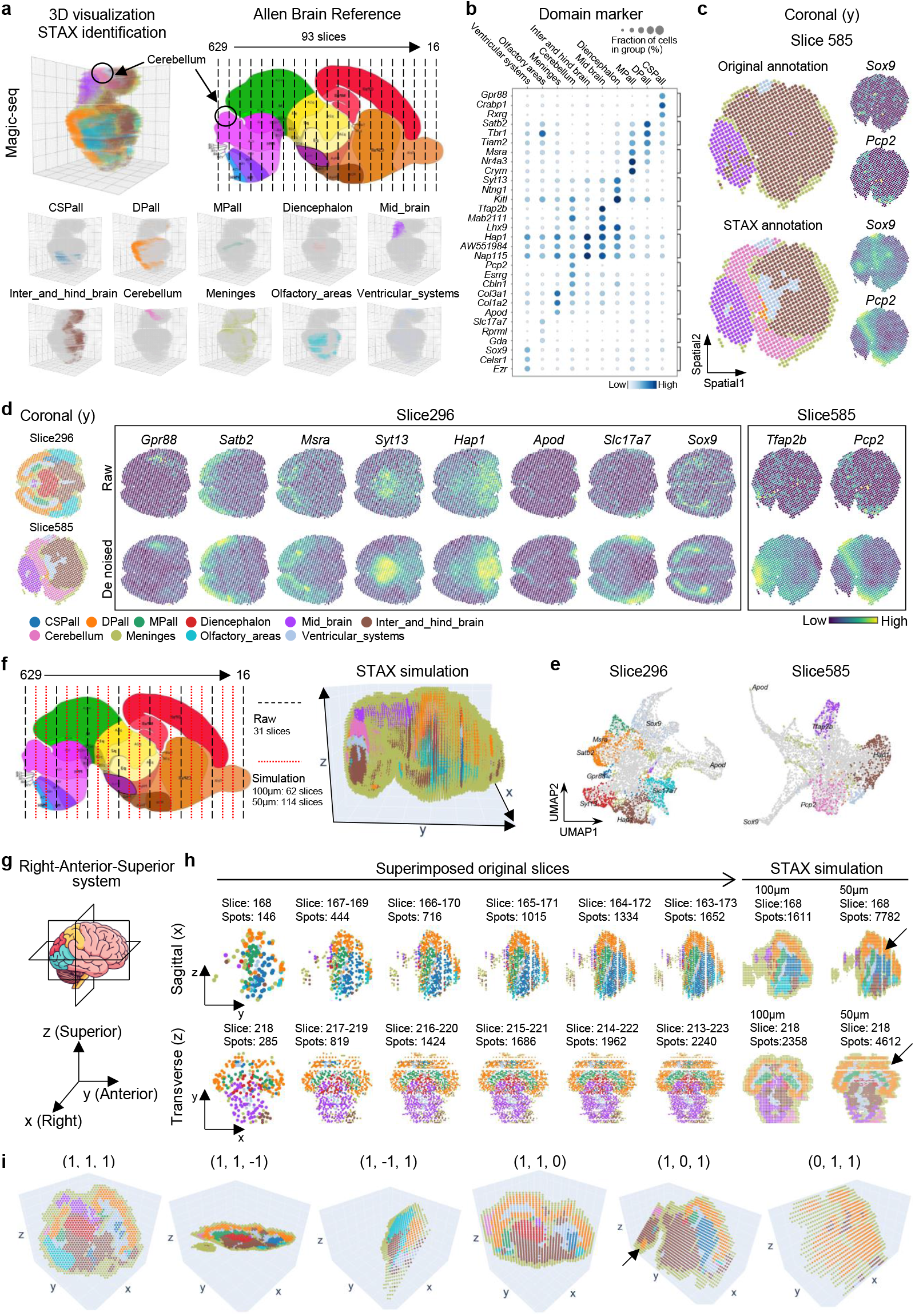
STAX generates a high-resolution 3D spatial transcriptomics atlas for the mouse embryo brain. **a**, Schematic overview of the Magic-seq workflow (right) and the corresponding 3D spatial domains reconstructed and identified by STAX (left). **b**, Dot plot illustrating the expression of the top three differentially expressed or previously confirmed genes enriched in brain regions. **c**, Spatial domains annotated by the original literature and STAX in slice 585. **d**, Comparison of the raw and STAX-denoised expression of marker genes across ten domains. **e**, UMAP plot demonstrating the co-embedding of cells and genes for slices 296 and 585. **f**, Schematic overview of how STAX generates 3D slices from spatial omics data. **g**, Illustration of the right-anterior-superior system. **h**, Visualization of the virtual slices from different anatomical perspectives, including sagittal and transverse views. **i**, Visualization of virtual slices generated using different slice orientations, defined by normal vectors (1, 1, 1) and (1, 1, –1).

STAX demonstrated the ability to denoise gene expression data. By reconstructing features, we mitigated the impact of dropouts to spatial sequencing technologies, thereby enhancing the clarity of spatial patterns and gene expression profiles (**Fig. 7d**). For validation, we utilized domain-specific genes reported in the original literature [64], After denoising, *Satb2* more accurately reflected the structure of DPall, *Sox9* exhibited clearer expression in the Ventricular systems, and *Syt13* and *Hap1* improved the differentiation between the Diencephalon and the Inter and Hindbrain regions.

Furthermore, through the UMAP visualization of the co-embedding of cells and genes, we observed that the markers and domains in slice 296 exhibited higher correspondence compared to slice 585. This discrepancy may be attributed to the presence of more refined structural categories in slice 296 (**Fig. 7e** and **Supplementary Fig. S32a**). In addition, some marker genes, such as *Msra*, did not perfectly align with their corresponding domains MPall. We hypothesize that this imperfect match may be attributable to the imprecision of domain identification and the high expression of *Msra* in both DPall and MPall (**Fig. 7b, d**).

To explore the potential of STAX in balancing experimental costs and precision, we simulated a slice generation scenario by reconstructing a refined 3D structure using low-resolution and a limited number of sparse slices. The right-anterior-superior system is a coordinate system that orients objects based on the observer’s perspective, with the positive x-axis pointing right, the positive y-axis pointing forward, and the positive z-axis pointing up (**Fig. 7g**). The Magic-seq technique performed vertical slicing along the y-axis, resulting in 93 slices with gap about 50μm (**Fig. 7a, right**; **f, left**). Firstly, along the y-axis, we replaced two out of every three original slices with virtual (fake) slices and performed super-resolution interpolation to enhance spatial resolution along the x and z axes at two resolutions: 100 μm and 50 μm resolution. Starting from approximately 30k spots, we generated a refined 3D map containing around 90k spots with 62 coronal slices (100μm) and 570k spots with 114 coronal slices (50μm), successfully identifying the same structures as those observed in the original high-resolution slices (**Fig. 7f, right** and **Supplementary Fig. S33**-**S38**). This simulation process is efficient, consuming 6 minutes for 90k spots and 9 minutes for 570k spots on a laptop with an RTX3070 GPU laptop (**Methods**).

Furthermore, we applied virtual slicing to the high-resolution 3D map from coronal, sagittal, and transverse perspectives. By generating sagittal and transverse slices from coronal slices, we achieved a cost reduction of approximately two-thirds. This approach maintains structural fidelity across different viewing angles. To validate the accuracy of the virtual slices, we examined the spatial patterns of marker genes. The changes in gene expression across cross-sections in all three perspectives further validated the effectiveness of STAX (**Supplementary Fig. S39a-c** and **S40a-c**). For example, in the coronal cross-section, the expression patterns of most genes exhibited a symmetrical distribution along the central axis, except for *Tfap2b* and *Pcp2*, which showed lower expression levels. In the sagittal and transverse cross-sections, cortical genes such as *Gpr88, Satb2*, and *Msra* were expressed more in the forebrain and expressed less in the midbrain and posterior brain. Conversely, genes such as *Tfap2b, Hap1*, and *Pcp2* displayed the opposite trend, with higher expression in the midbrain and posterior brain and lower expression in the forebrain. More importantly, a complete structural domain could not be adequately captured from sagittal and transverse angles using original slices alone, necessitating the superposition of multiple slices (**Fig. 7h, left**). A single virtual slice generated by STAX—with its uniformly sampled point cloud—achieves a level of structural detail comparable to that of up to 11 overlaid original slices (**Fig. 7h, right** and **Supplementary Fig. S36–S38**). Additionally, by using a point’s coordinates and a normal vector, we can generate virtual slices at any angle (**Fig. 7i**). Overall, the above observations align with known biological patterns, showing that STAX can generate high-resolution virtual slices.

## Discussion

The emergence of spatial omics technologies has fundamentally reshaped the paradigms of biomedical research. Nevertheless, the intrinsic complexities stemming from data variability and technical limitations have substantially impeded spatial data processing. This has given rise to a multitude of analytical challenges in spatial omics, encompassing spatial domain identification, spatial slice integration, spatial cohort analysis, cell-gene co-embedding, spatial data denoising, spatial spot completion, and 3D reconstruction of multi-slice spatial data. Although some algorithms have been developed to tackle these issues, their task-specific nature often compromises the efficiency and consistency of data analysis. To address these constraints, we introduce STAX, a novel multi-task deep learning framework designed to resolve these challenges through a unified approach.

By leveraging a meticulously designed architecture, we have seamlessly integrated multiple tasks into a unified framework, STAX, which enables comprehensive functionality while consistently delivering exceptional performance across diverse benchmark tests. When considering only self-loop connections, STAX achieves precise clustering analysis at the cellular level. By incorporating physical spatial relationships, STAX can further enhance its clustering capability to the domain level. STAX is a lightweight framework that supports mini-batch training, ensuring efficient scalability to datasets of varying scales and resolutions. Remarkably, all aforementioned tasks, including cohort analyses and 3D simulation tasks, were successfully executed on a personal laptop equipped with an i7-11800H CPU, 64 GB of RAM, and an NVIDIA GeForce RTX 3070 Laptop GPU with 8 GB of dedicated memory (**Methods**). We are confident that by utilizing larger datasets and more powerful computational resources, the adaptability of STAX will further solidify its position as a spatial foundation model, capable of addressing a broader range of downstream tasks and providing robust initialization representations for subsequent analyses.

While STAX demonstrates competitive capabilities, several limitations present opportunities for future enhancement. First, its performance exhibits sensitivity to key hyperparameters, particularly those related to neighborhood size in graph construction. The selection of clustering methods based on cell and gene embeddings also significantly impacts the analytical outcomes, requiring careful optimization. Secondly, the virtual slices from STAX do have limitations. For example, large gaps between some slices make it impossible to reconstruct the entire structure (**Fig. 7h**, arrow). This is likely due to the inherent smoothing characteristic of GNNs and the omission of distance weighting during graph construction. Thirdly, the current implementation is constrained to unimodal data analysis, with node representations limited to single-omics expressions. To address these limitations, we propose several strategic directions for future development: (1) establishing unified graph construction methodologies coupled with standardized post-processing step including clustering algorithms to streamline spatial omics data analysis pipelines; (2) incorporating distance weighting into graph construction or adopting distribution-based sampling approaches, such as diffusion or flow models, as alternatives to GNN-based interpolation; (3) developing a multi-modality architecture capable of seamless cross-omics integration; and (4) leveraging the inherent extensibility of STAX to construct spatial-based foundation models that provide robust initial representations for more downstream analytical tasks. Collectively, these enhancements are expected to improve the model’s applicability, computational efficiency, and biological relevance, enabling more comprehensive and insightful interpretation of increasingly complex spatial omics data.

## Methods

### STAX model

#### Feature construction

STAX supports multiple forms of one given type of omics feature input, e.g., expression count matrix, feature embeddings (e.g., outputs from PCA (Principal Component Analysis), LSI (Latent Semantic Indexing), or other methods), and one-hot encodings of cell types (or subtypes) (**Supplementary Fig. S1a**). While STAX does not enforce a unified data preprocessing or feature construction pipeline, it provides a comprehensive suite of built-in functions to facilitate user customization. For example, for transcriptomic expression matrices, the workflow typically begins with optional feature selection, which can be performed using the ‘scanpy.pp.highly_variable_genes’ function, followed by optional normalization and scaling steps, e.g., log transformation, z-score standardization, or min-max scaling.

Generally, feature construction is a step that often requires substantial expertise and abundant experience. For instance, filtering low-quality cells or genes with low expression levels is not always mandatory, as spatial proteomics data (e.g., CODEX) often measure only a limited range of protein expression. In the case of spatial epigenomics, such as spatial ATAC-seq data, each sample may contain hundreds of thousands of features, necessitating dimensionality reduction techniques like LSI transformation as an initial step. Ultimately, feature construction is user-defined, allowing researchers to iteratively refine their preprocessing pipeline based on the specific characteristics of their dataset and experimental scenarios.

#### Graph construction

STAX supports the construction of a neighbor graph, i.e., physical adjacent graphs (**Supplementary Fig. S1a**). Based on the hypothesis that spatially proximate nodes (spots or cells) exhibit similar expression patterns or interactions, we construct a physical adjacent graph. Specifically, for a given central spot or cell, we connect it to all neighboring spots or cells within a radius r. Alternatively, we also support a k-nearest neighbors (KNN) approach, where the central node is connected to its *k* closest neighbors.

Note that the physical adjacent graph is optional. If it is not selected, the graph will consist solely of self-loop edges, causing the graph autoencoder to degenerate into a standard autoencoder, and the resulting embeddings will correspond to cell-level clustering. When the Physical Adjacent Graph is selected (with edges combined via union), the embeddings will reflect domain-level clustering.

#### Attribute graph

The features and graphs are assembled into an attributed graph. Given the rapid advancement of spatial omics technologies, the number of cells or spots per slice, as well as the number of slices has significantly increased. To address this, we introduce a graph mini-batch strategy.

Specifically, during each epoch, the entire graph is randomly partitioned into subgraphs using ‘dgl.dataloading.ShaDowKHopSampler’ in DGL v2.2.1 [66]. Training is then performed on these subgraphs, with each mini-batch processed independently. By the end of the epoch, all subgraphs are trained, ensuring comprehensive coverage of the data. However, for smaller slices, we still recommend using a full-batch training strategy to preserve the complete structural information of the entire graph. For full-batch training, the default number of epochs is set to 500. When conducting mini-batch training, it is recommended to adjust the parameters such that the total number of cells, n, is divided by 100 (i.e., n/100), and the number of epochs is set to 500/100 = 5. If your computer continues to encounter memory limitations, you can further increase the value as needed. For instance, setting it to n/200 or lower can help alleviate memory constraints.

### Model architecture and loss function

#### Domain identification

To address the task of domain identification, we propose a graph Variational AutoEncoder (VAE) with a multi-layer GAT as both the encoder and decoder. Our model is designed to learn a robust latent representation of the input attribute graph, where we encode node features *x* to an embedding *z* and subsequently reconstruct the features *x*^′^ from *z*.

The goal of VAE is to maximize the log-likelihood of the data *log p*(*x*). However, since directly computing *log p*(*x*) is intractable, VAE introduces an approximate posterior distribution *q*(*z*|*x*) through variational inference and derives the evidence lower bound:

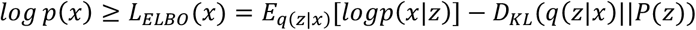

where *logp*(*x*|*z*) is the decoder part, representing the reconstruction probability of data *x* given the latent variable *z*. Maximizing this term improves the model’s ability to reconstruct data. During spot completion, we calculate the loss exclusively for real spots while disregarding the loss associated with fake spots. *D*_*KL*_(*q*(*z*|*x*)||*P*(*z*)) is the Kullback-Leibler (KL) divergence, measuring the difference between the approximate posterior distribution *q*(*z*|*x*) and the prior distribution *P*(*z*).

Both the encoder and decoder are implemented as multi-layer GATs. The computation steps for the GAT are as follows. First, the attention coefficients are calculated. For a given vertex or node, the similarity coefficients between itself and each of its neighboring vertices are computed sequentially. The attention mechanism computes similarity coefficients for a given vertex *i* and its neighbors *j* ∈ *N*_*i*_ as follows:

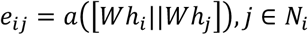

where *W* is a shared parameter matrix to map the feature into a shared space, || is the concatenate operation and *a* is to map the concatenated vector into a real number. These attention coefficients are then normalized using the softmax function to ensure they sum to one across all neighbors. Then, normalization is implemented through softmax. These similarity coefficients are then normalized using the softmax function to obtain attention weights:

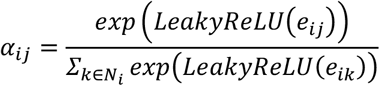

Subsequently, based on the calculated attention coefficients, a weighted sum of the features is computed through aggregation as follows:

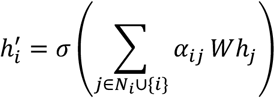

To enhance the model’s robustness, we employ a multi-head attention mechanism. Specifically, for *K* attention heads, the process is repeated independently, and the results are combined to produce the final output.

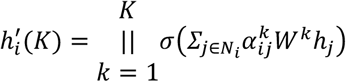

where || is the is the concatenate operation.

#### Batch correction

When dealing with multiple slices, batch effects are inevitably encountered. Drawing inspiration from prior work on domain-specific batch normalization (DSBN) for domain adaptation [67] and batch effect removal [68], we apply it in the decoder and assign a unique batch normalization layer to each batch.

#### Cell and gene co-embedding

The embedding of genes does not require additional training. We use the parameter matrix of the decoder to initialize the gene embeddings. Specifically, when the dimension of the hidden layer is *z* and the dimension of the input *x* or output *x*^′^ is *m*, and *m* represents the number of genes. The parameter of the decoder is *W* ∈ *R*^*z*×*m*^, and we treat *W*^*T*^ ∈ *R*^*m*×*z*^ as the initial gene embedding. To make the embeddings of cells and genes comparable, we employ a dynamic embedding strategy inspired by the Softmax transformation proposed in the reference [29]. This method generates new feature embeddings that are a linear combination of the cell embeddings, weighted by their estimated edge probabilities. Specifically, given the initial embeddings of cells (reference) 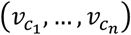 from bottle neck layer and genes (query) 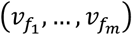 from *W*^*T*^, the model-estimated probability *P* of an edge (*c*_i_, *c*_j_) obeys:

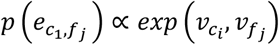

We then normalize these probabilities using a softmax transformation, which yields the probability 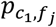 of an edge from gene *f*_*j*_ to cell *c*_1_ over all cells:

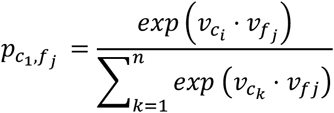

Subsequently, we compute the new embedding 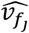 for each gene *f*_*j*_ as a weighted sum of all cell embeddings. The weights are the estimated edge probabilities 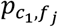 raised to a power controlled by a temperature hyperparameter *T*:

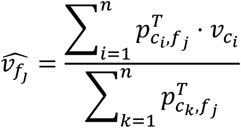

where *T* is a hyperparameter and we set it to 0.1 by default.

#### Spot completion

The spot completion technique incurs no extra computational overhead. It begins by generating fake spots near existing real spots. The subsequent training process focuses exclusively on the losses from the real spots, with the losses of the fake nodes being disregarded. For a node *i*, the loss function *L*_*i*_ is the sum of its reconstruction loss *L*_*recon,i*_ and its KL divergence loss *L*_*KL,i*_. We introduce an indicator function *I*(·).

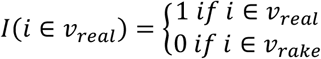

This function takes a value of 1 for nodes in the real node set and 0 for nodes in the fake node set, which is equivalent to a Boolean mask in programming. So, the final loss term is defined as:

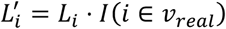

#### Clustering

We used different clustering methods including mclust [69], Louvain [37], Leiden [38] and kmeans [70, 71] to identify spatial domains based on embeddings of STAX.

#### Differentially genes identification

We used the t-test implemented in Scanpy package to identify differentially expressed genes for each spatial domain [38].

#### GO enrichment analysis

We perform the GO enrichment analysis by gprofiler [72].

#### Clustering coefficient

We calculate the clustering coefficient by networkx [73].

### Evaluation metrics

#### Adjusted Rand Index (ARI)

ARI measures the degree of similarity between two clustering results by counting the number of sample points in clusters of the same class and clusters of different classes. Here, we calculated the ARI score using the scikit-learn package [70]. The value range of ARI is [-1,1]. The closer ARI is to 1, the more similar the two clustering results are. ARI can be defined as follows:

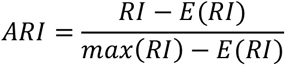

where *E*(*RI*) represents the expectation of the unadjusted Rand index *RI*, where can be defined as 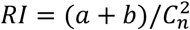. Here, *a* denotes the number of pairs correctly identified as belonging to the same cluster, *b* denotes the number of pairs correctly identified as belonging to different clusters, and 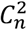 denotes the total number of possible pairs in the dataset.

#### Moran’s I and Geary’s C

Moran’s I and Geary’s C score assesses whether the pattern is clustered, dispersed, or random in physical location. Here, we calculated Moran’s I and Geary’s C score using the Squidpy package [74]. Briefly, given a feature (gene, embedding, or label) and the spatial location of observations. Specifically, Moran’s I can be defined as:

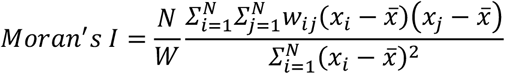

where *N* represents the number of spots or cells, *w*_*ij*_ represents the spatial weight between observations, *W* represents the sum of the spatial observations, and equals 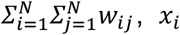 and *x*_*j*_ represents the feature, 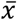 represents the mean of all *x*_*i*_ and *x*_*j*_. Geary’s C can be defined as:

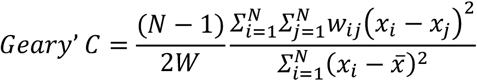

For embedding, we calculated Moran’s I and Geary’s C for each dimension of the embedding and took the average. The embedding dimensions of STAX, STAGATE, and SpatialGlue are 16, 30, and 20, respectively. For the label, we calculated Moran’s I and Geary’s C for each spatial cluster label and took the average.

#### (Local) Clustering Coefficient

The (local) clustering coefficient is used to measure the degree of connectivity between neighbors of a node. For node *v*, the clustering coefficient can be defined as follows:

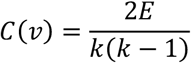

where *E* represents the actual number of edges that exist between neighbors of node *v, k* is the number of neighbors of node *v*. Here, we use the networkx.clustering [73] function to calculate it.

#### Time consuming

For a fair comparison, we use the total time consumed by the model, including data loading, data preprocessing, and model training, as the model’s overall time consumption.

## Data Availability

To facilitate accessibility and usability for other researchers, we have consolidated these datasets into the h5ad format and uploaded them to a cloud storage platform: Google Driver (https://drive.google.com/drive/folders/18tcl-PRdK9j-W59GUPdsKgy_04IJvz05). This resource is intended to support the broader scientific community in their modeling efforts. The details of the data can be seen in **Supplementary Fig. S1b**.

Furthermore, all data analyzed in this study are available from their respective original publications as follows:

## Single omics datasets in the domain identification task

Mouse brain dataset 1 (epigenomics): https://web.atlasxomics.com/visualization/Fan/

Human dorsolateral prefrontal cortex dataset (DLPFC, transcriptomics): https://research.libd.org/spatialLIBD/.

Mouse brain dataset 2 (translatomics): https://singlecell.broadinstitute.org/single_cell/study/SCP1835. Mouse spleen dataset (proteomics): https://data.mendeley.com/datasets/zjnpwh8m5b/1.

Pig embryo dataset (metabolomics):

R cardinal package (https://www.bioconductor.org/packages/CardinalWorkflows/).

## Mouse olfactory bulb datasets across seven technologies

ST (100µm):

https://www.spatialresearch.org/resources-published-datasets/doi-10-1126science-aaf2403/.

10x Visium (55µm): https://www.10xgenomics.com/cn/datasets/adult-mouse-olfactory-bulb-1-standard.

Array-seq (30µm): https://www.ncbi.nlm.nih.gov/geo/query/acc.cgi?acc=GSE266244.

Decoder-seq (15µm): https://www.ncbi.nlm.nih.gov/geo/query/acc.cgi?acc=GSE235896.

Slide-seq (10µm): https://portals.broadinstitute.org/single_cell/study/slide-seq-study.

Stereo-seq (1µm): https://db.cngb.org/stomics/datasets/STDS0000058.

STARmap (single cell) https://zenodo.org/records/8327576.

## Spatial transcriptomics ovarian cancer cohort dataset

The dataset can be accessed at https://singlecell.broadinstitute.org/single_cell/study/SCP2640/hgsc-spatial-cohort-discovery-dataset#study-summary.

## Liver cancer cohort codex dataset

The dataset can be accessed at https://data.mendeley.com/datasets/km2df7y256/2.

## Breast cancer Xenium and Visium dataset

The dataset can be downloaded at https://www.10xgenomics.com/products/xenium-in-situ/preview-dataset-human-breast.

## Mouse hypothalamic preoptic region MERFISH dataset

The original cell type annotated dataset can be accessed at https://datadryad.org/stash/dataset/doi:10.5061/dryad.8t8s248.

The domain dataset annotated by Yuan et al. can be accessed at http://sdmbench.drai.cn/ or https://figshare.com/projects/SDMBench/163942.

## The normal and Alzheimer’s disease mouse hippocampus Slide-seqV2 dataset

The dataset can be accessed at the Broad Institute portal at https://singlecell.broadinstitute.org/single_cell/study/SCP815 and https://singlecell.broadinstitute.org/single_cell/study/SCP1663, respectively.

## Mouse embryo brain Magic-seq dataset

The dataset can be accessed at https://zenodo.org/records/13934668.

## Mouse thymus protein dataset

The dataset can be accessed at https://zenodo.org/records/10362607.

## Code Availability

The code can be obtained from https://github.com/zhanglabtools/STAX and https://github.com/CocoGzh/STAX

## Computational resource

CPU: 11th Gen Intel(R)Core(TM)i7-11800H 2.30GHz.

Memory: 64 GB in total.

GPU: NVIDIA GeForce RTX 3070 Laptop GPU, 8 GB memory.

## Ethics approval and consent to participate

Not applicable.

## Consent for publication

Not applicable.

## Author Contributions

S.Z. and Z-H.G conceived and supervised the project. Z-H.G. collected the datasets and developed the STAX algorithm. Z-H.G., D.-S.H., and S.Z. validated the methods, performed the analysis, and wrote the manuscript. All authors read and approved the final paper.

## Acknowledgements

The authors would like to thank all editors and reviewers for their helpful suggestions.

