## Supplemental Figures for "Multi-task spatial omics analytics for high-precision modeling of tissue architecture with STAX"

**a**

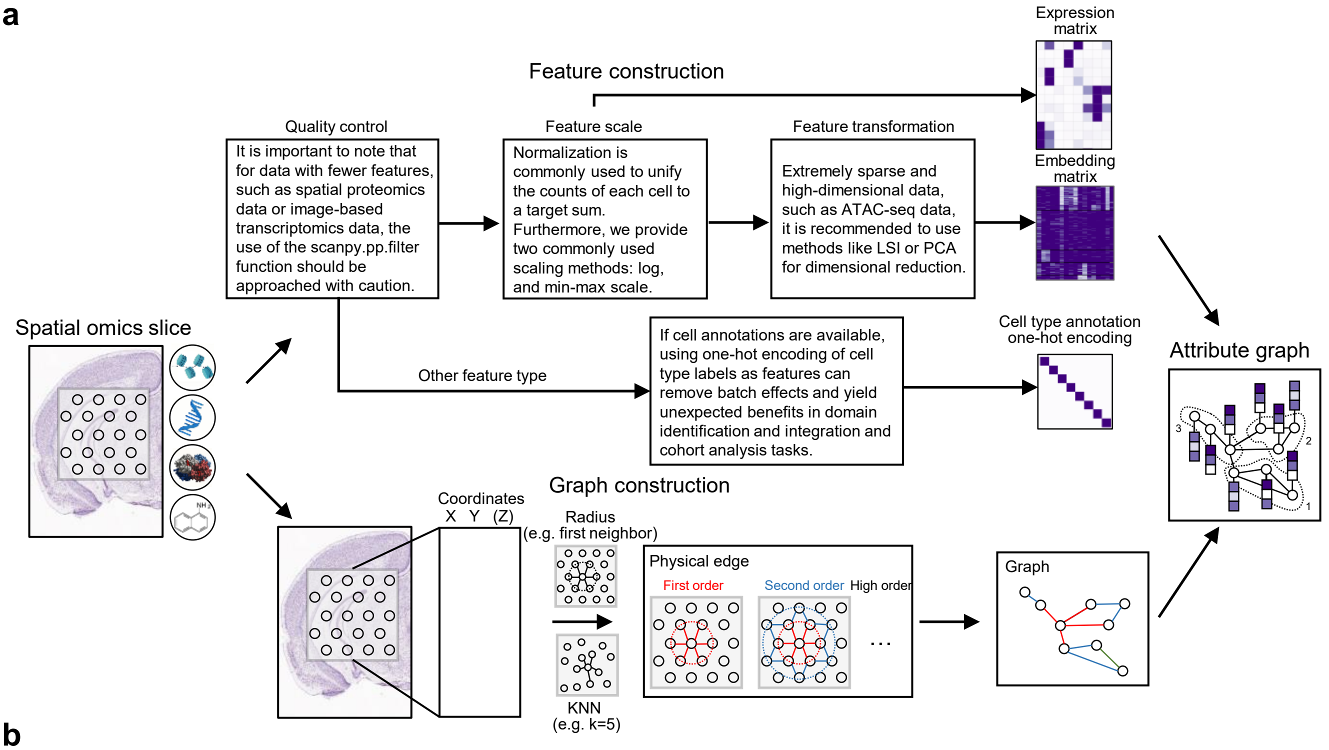


**b**

| Sec | Dataset | Specie | Organ or tissue | Technology | Omics | Resolution | Cell or spot | Feature | Preprocess |
|---|---|---|---|---|---|---|---|---|---|
| 1 | Mouse brain dataset 1 | Mouse | Brain | 10X Visium | Epigenomics | 55μm | 9,215 | 121,068 | LSI |
| | DLPFC | Human | Brain | 10X Visium | Transcriptomics | 55μm | ~4,000 | 33,538 | Norm & log |
| | Mouse brain dataset 2 | Mouse | Brain | RIBOmap | Transcriptomics | Single cell | 60,481 | 5,413 | Norm & log |
| | Mouse spleen dataset | Mouse | Spleen | CODEX | Proteomics | Single cell | ~80,000 | 30 | MinMax |
| | Pig embryo dataset | Pig | Embryo | MALDI-MSI | Metabolomics | Variable (e.g., 50μm) | 4,959 | 10,200 | Norm & log |
| 2 | MOB1 | Mouse | Olfactory bulb | ST | Transcriptomics | 100μm | 262 | 16,218 | Norm & log |
| | MOB2 | Mouse | Olfactory bulb | 10x Visium | Transcriptomics | 55μm | 1,185 | 32,285 | Norm & log |
| | MOB3 | Mouse | Olfactory bulb | Array-seq | Transcriptomics | 30μm | 7,716 | 17,243 | Norm & log |
| | MOB4 | Mouse | Olfactory bulb | Decoder-seq | Transcriptomics | 15μm | 1,710 | 26,736 | Norm & log |
| | MOB5 | Mouse | Olfactory bulb | Slide-seq V2 | Transcriptomics | 10μm | 20,139 | 21,220 | Norm & log |
| | MOB6 | Mouse | Olfactory bulb | Stereo-seq | Transcriptomics | 1μm | 19,109 | 27,106 | Norm & log |
| | MOB7 | Mouse | Olfactory bulb | STARmap | Transcriptomics | Single cell | 19,870 | 1,022 | Norm & log |
| 3 | Transcriptomics cohort | Human | Ovary | SMI | Transcriptomics | Single cell | 491,792 | 13 | Cell type one-hot |
| | Proteomics cohort | Human | Liver | CODEX | Proteomics | Single cell | 2,631,808 | 9 | Cell type one-hot |
| 4 | Breast cancer Visium | Human | Breast | 10X Visium | Transcriptomics | 55μm | 3,955 | 18,085 | Norm & log |
| | Breast cancer Xenium | Human | Breast | 10X Xenium | Transcriptomics | Single cell | 159,226 | 313 | Norm & log |
| 5 | Visual cortex dataset | Mouse | visual cortex | MERFISH | Transcriptomics | Single cell | 28,317 | 155 | Norm & log |
| | Hippocampus AD | Mouse | Hippocampus | Slide-seq | Transcriptomics | 10μm | 15,092 | 14,622 | Norm & log |
| | Hippocampus Normal | Mouse | Hippocampus | Slide-seq | Transcriptomics | 10μm | 12,822 | 14,622 | Norm & log |
| 6 | Mouse embryo brain | Mouse | Brain | MAGIC-seq | Transcriptomics | 50μm | 97,818 | 21,311 | Norm & log |

**Fig. S1.** Summary of data preprocessing and datasets. **a**, Details of feature preprocessing and network construction in STAX. **b**, Basic information of the datasets used in this study.

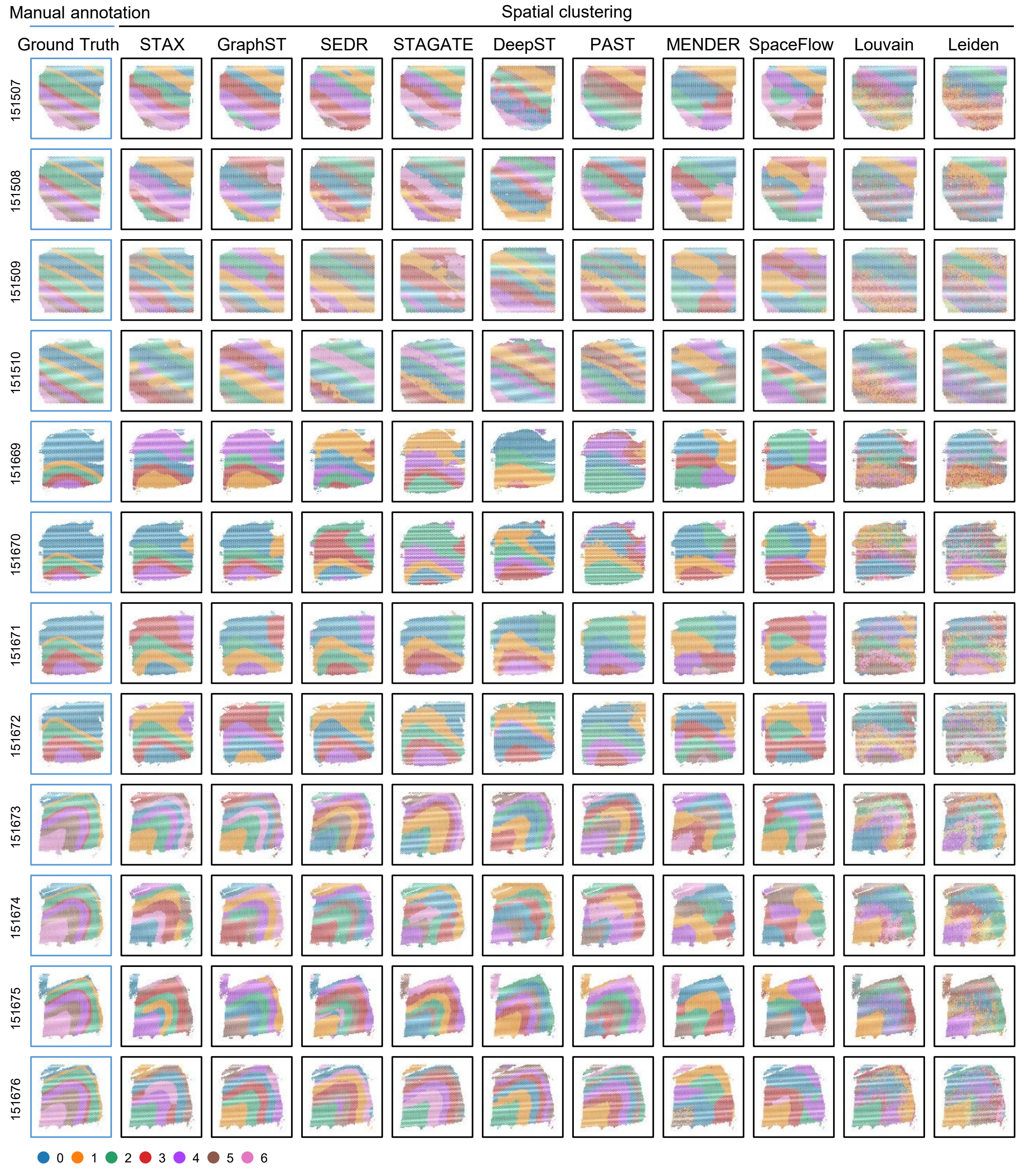


**Fig. S2.** Manual annotation and spatial domains identified by STAX and baseline methods including GraphST, SEDR, STAGATE, DeepST, PAST, MENDER, SpaceFlow, Louvain and Leiden.

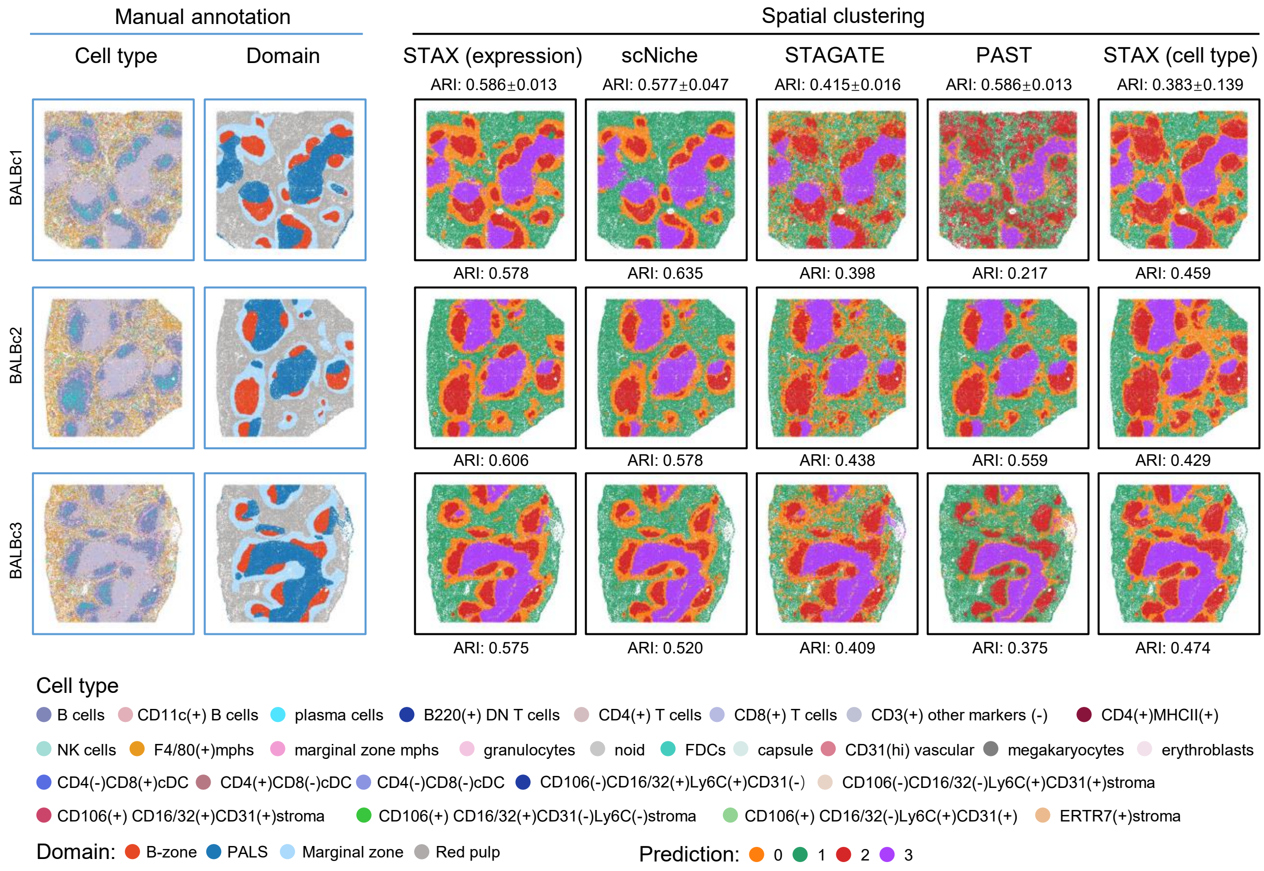


**Fig. S3.** Manual annotation for cell type and domains and spatial domains identified by STAX (protein expression) and baselin methods including scNiche, STAGATE, PAST and STAX (cell type one-hot encoding) on the mouse spleen dataset.

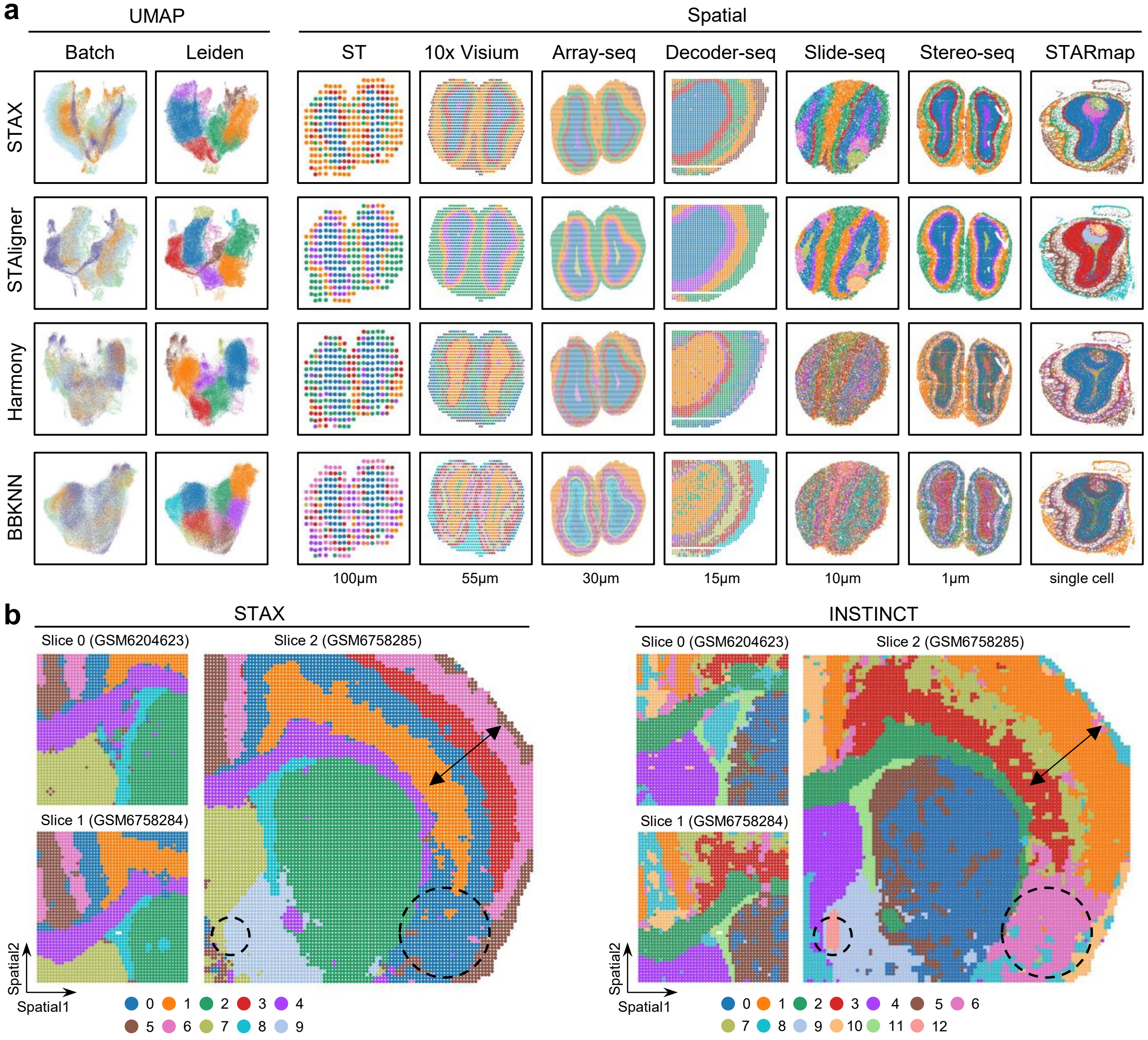


**Fig. S4.** Integration results of STAX and baseline methods on the mouse brain datasets. **a**, The integration results of STAX, STAligner, Harmony and BBKNN on the datasets across seven technologies. **b**, The integration results of STAX and INSTINCT in the mouse brain ATAC-seq dataset across three slices.

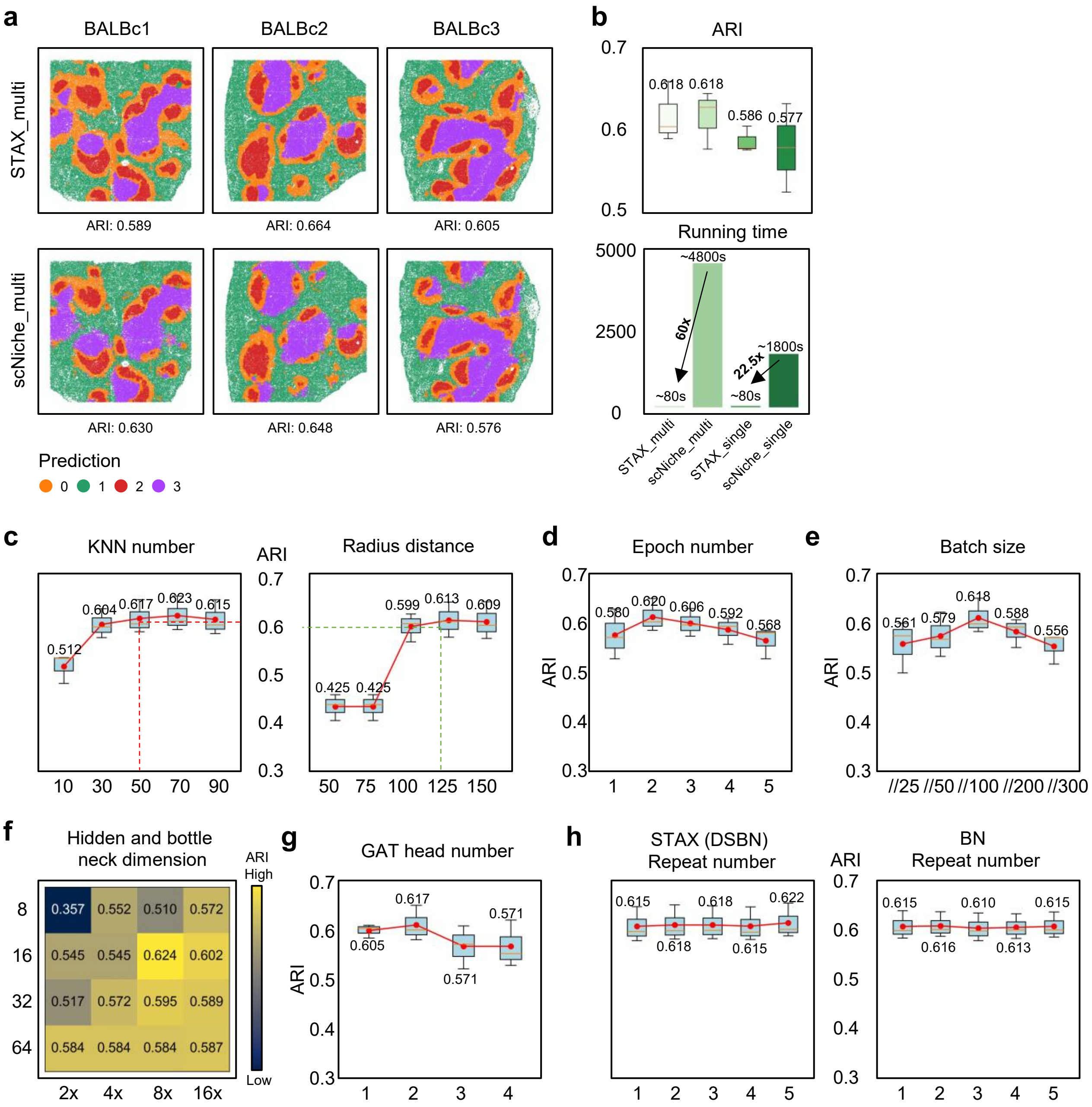


**Fig. S5.** Performance evaluation of STAX and baseline methods on the mouse spleen dataset. **a**, The integration results of STAX, scNiche in the CODEX dataset across three slices. **b**, Comparison of ARI and running time between STAX and scNiche. **c-g**, Hyperparameter comparisons in STAX including the KNN number, Radius distance, epoch number, batch size, hidden and bottleneck dimensions, and the GAT head number. **h**, Ablation experiments of DSBN and BN.

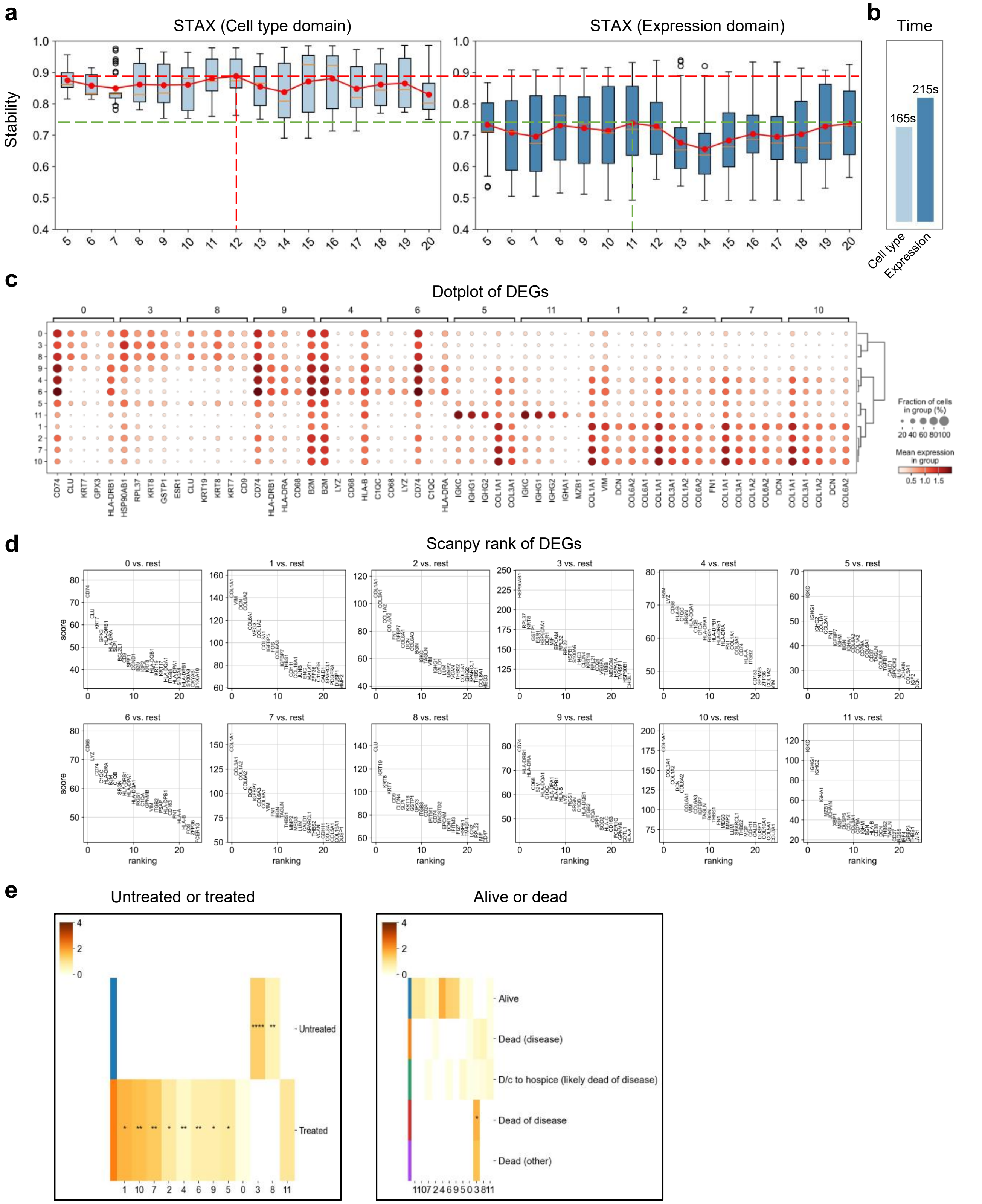


**Fig. S6.** Results of STAX on the spatial transcriptomics cohort study. **a**, Stability score of STAX when using the cell type one-hot encoding and expression as the feature. **b**, Time consumption of STAX when using the cell type one-hot encoding and expression as the feature. **c**, Dot plot of top 3 differentially expressed genes for each cluster. **d**, Differentially expressed genes identified by Scanpy. **e**, Enrichment scores of different domains calculated by scNiche in different clinical groups.

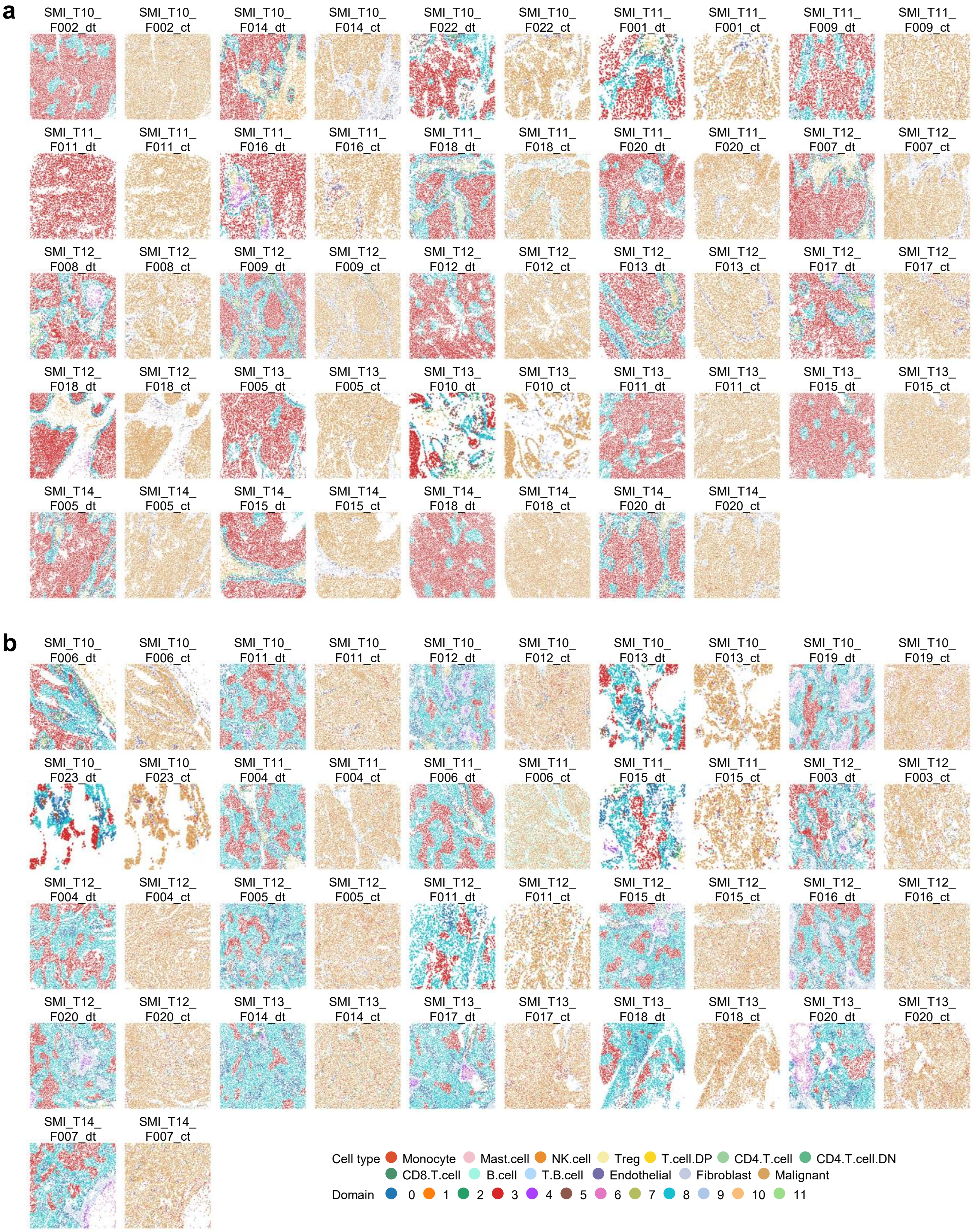


**Fig. S7.** Results of STAX on the spatial transcriptomics cohort study. **a** and **b**, Cell types annotation (right) and domains (left) identified by STAX in groups 1 (**a**) and 2 (**b**) of the hierarchical clustering, where "_dt" indicates the domain type, and "_ct" indicates the cell type.

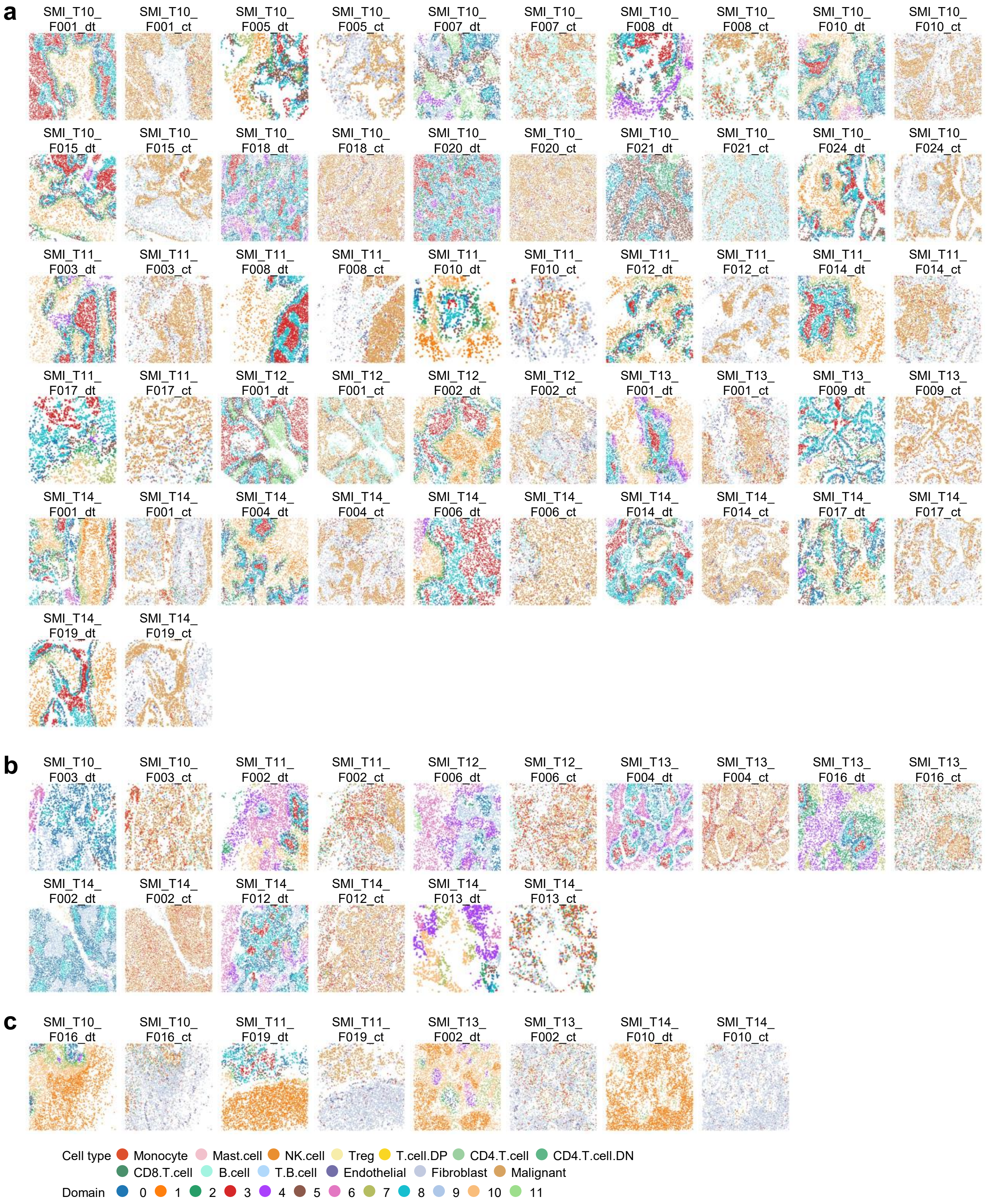


**Fig. S8.** Results of STAX on the spatial transcriptomics cohort study. **a-c**, Cell-type annotation (right) and domains (left) identified by STAX in groups 3 (**a**), 4 (**b**), 5 (**c**) of the hierarchical clustering, where “_dt” indicates the domain type, and “_ct” indicates the cell type.

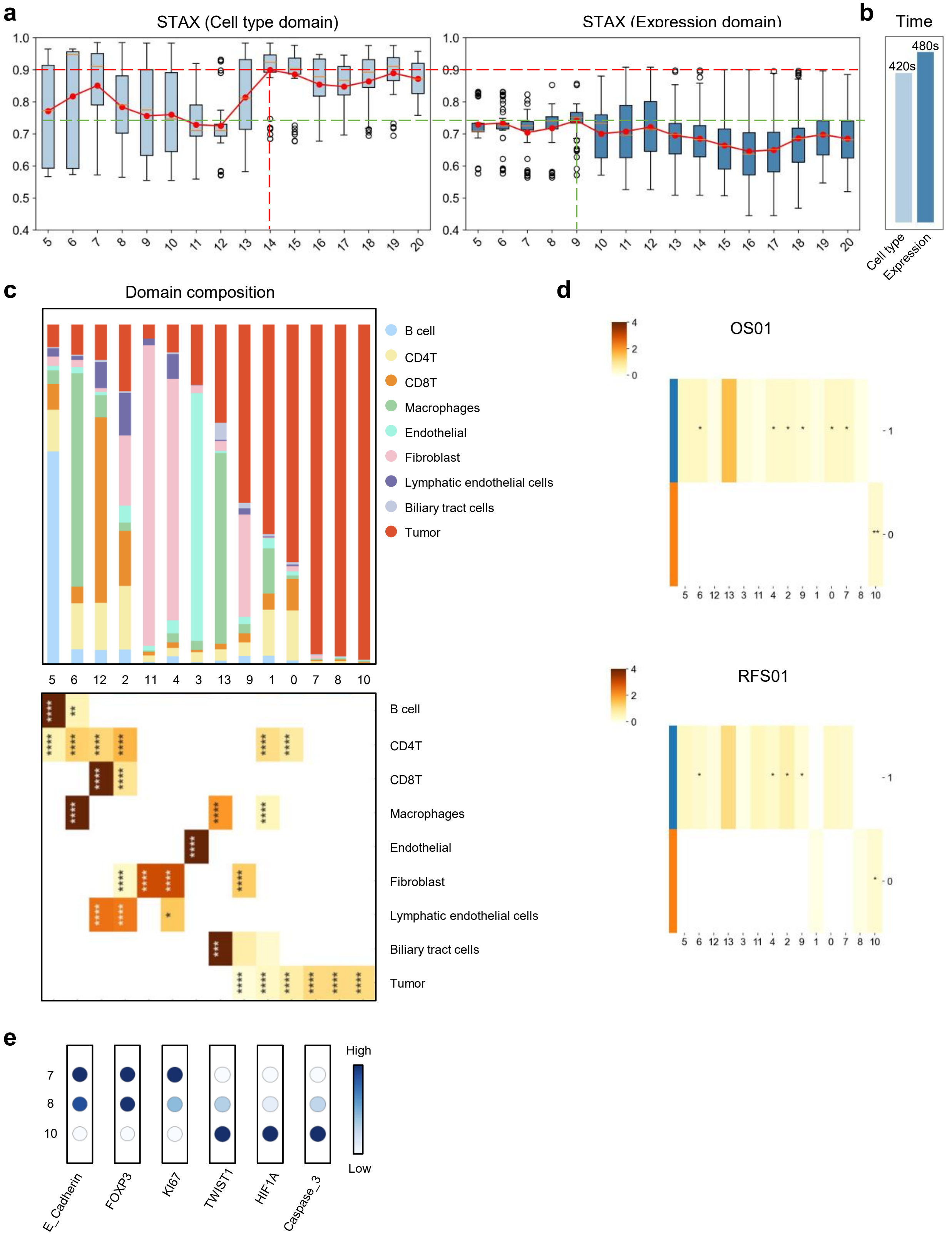


**Fig. S9.** Results of STAX on the spatial proteomics cohort study. **a**, Stability score of STAX when using cell type one-hot encoding and expression as the feature. **b**, Time consumption of STAX when using cell type one-hot encoding and expression as the feature. **c**, The cell type composition of each domain (up), as well as the enrichment scores of the cell types identified by scNiche in the different domains (down). **d**, The enrichment scores of different domains calculated by scNiche in different clinical groups. **e**, Expression of E_Cadherin, FOXP3, KI67, Twist1, HIF1a, Caspase_3 in domains 7, 8, and 10.

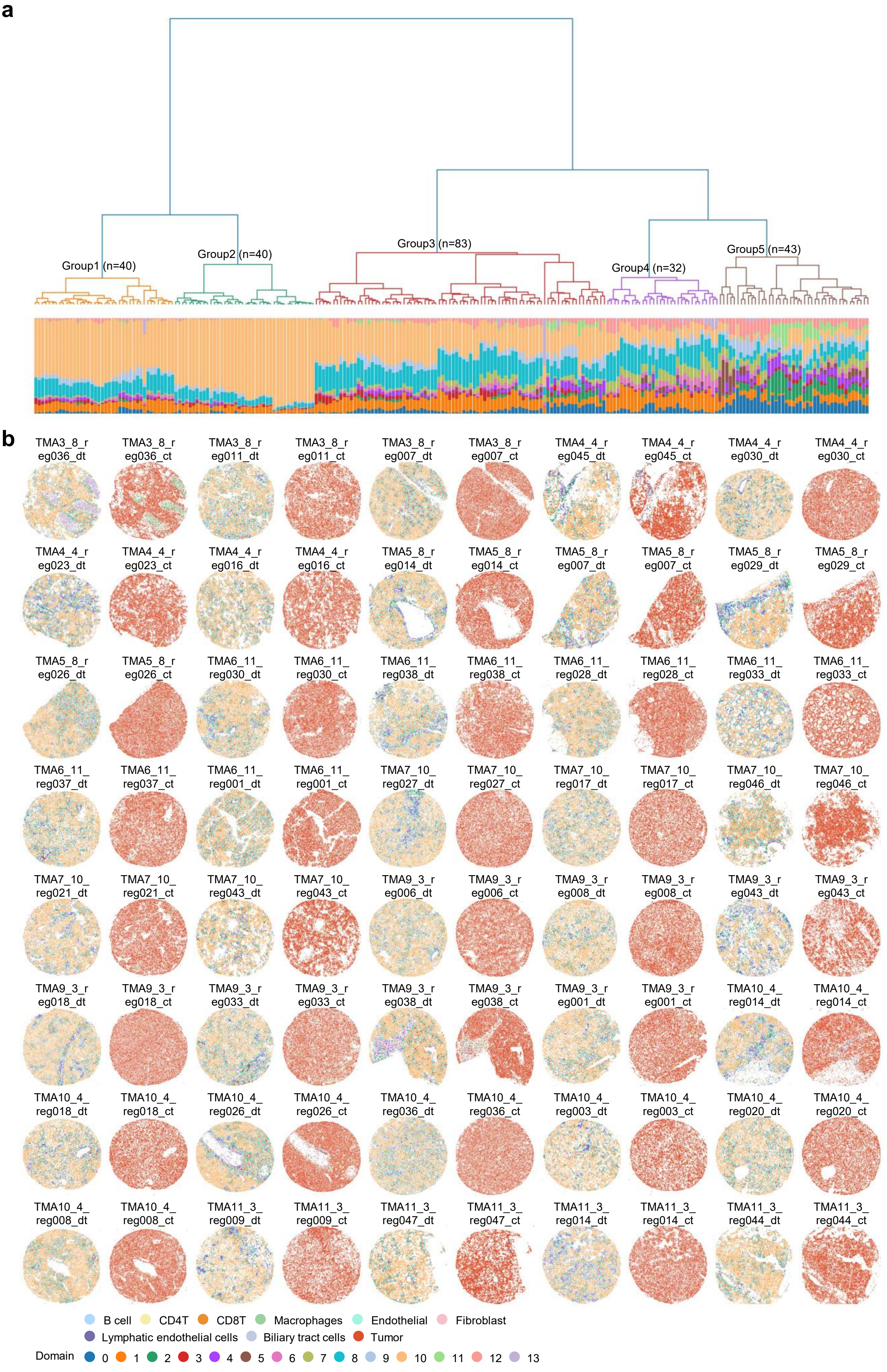
a
Group1 (n=40)
Group2 (n=40)
Group3 (n=83)
Group4 (n=32)
Group5 (n=43)
b
TMA3_8_r eg036_dt
TMA3_8_r eg036_ct
TMA3_8_r eg011_dt
TMA3_8_r eg011_ct
TMA3_8_r eg007_dt
TMA3_8_r eg007_ct
TMA4_4_r eg045_dt
TMA4_4_r eg045_ct
TMA4_4_r eg030_dt
TMA4_4_r eg030_ct
TMA4_4_r eg023_dt
TMA4_4_r eg023_ct
TMA4_4_r eg016_dt
TMA4_4_r eg016_ct
TMA5_8_r eg014_dt
TMA5_8_r eg014_ct
TMA5_8_r eg007_dt
TMA5_8_r eg007_ct
TMA5_8_r eg029_dt
TMA5_8_r eg029_ct
TMA5_8_r eg026_dt
TMA5_8_r eg026_ct
TMA6_11_ reg030_dt
TMA6_11_ reg030_ct
TMA6_11_ reg038_dt
TMA6_11_ reg038_ct
TMA6_11_ reg028_dt
TMA6_11_ reg028_ct
TMA6_11_ reg033_dt
TMA6_11_ reg033_ct
TMA6_11_ reg037_dt
TMA6_11_ reg037_ct
TMA6_11_ reg001_dt
TMA6_11_ reg001_ct
TMA7_10_ reg027_dt
TMA7_10_ reg027_ct
TMA7_10_ reg017_dt
TMA7_10_ reg017_ct
TMA7_10_ reg046_dt
TMA7_10_ reg046_ct
TMA7_10_ reg021_dt
TMA7_10_ reg021_ct
TMA7_10_ reg043_dt
TMA7_10_ reg043_ct
TMA9_3_r eg006_dt
TMA9_3_r eg006_ct
TMA9_3_r eg008_dt
TMA9_3_r eg008_ct
TMA9_3_r eg043_dt
TMA9_3_r eg043_ct
TMA9_3_r eg018_dt
TMA9_3_r eg018_ct
TMA9_3_r eg033_dt
TMA9_3_r eg033_ct
TMA9_3_r eg038_dt
TMA9_3_r eg038_ct
TMA9_3_r eg001_dt
TMA9_3_r eg001_ct
TMA10_4_ reg014_dt
TMA10_4_ reg014_ct
TMA10_4_ reg018_dt
TMA10_4_ reg018_ct
TMA10_4_ reg026_dt
TMA10_4_ reg026_ct
TMA10_4_ reg036_dt
TMA10_4_ reg036_ct
TMA10_4_ reg003_dt
TMA10_4_ reg003_ct
TMA10_4_ reg020_dt
TMA10_4_ reg020_ct
TMA10_4_ reg008_dt
TMA10_4_ reg008_ct
TMA11_3_ reg009_dt
TMA11_3_ reg009_ct
TMA11_3_ reg047_dt
TMA11_3_ reg047_ct
TMA11_3_ reg014_dt
TMA11_3_ reg014_ct
TMA11_3_ reg044_dt
TMA11_3_ reg044_ct
B cell
CD4T
CD8T
Macrophages
Endothelial
Fibroblast
Lymphatic endothelial cells
Biliary tract cells
Tumor
Domain
0
1
2
3
4
5
6
7
8
9
10
11
12
13

**Fig. S10.** Results of STAX on the spatial proteomics cohort study. **a**, Hierarchical clustering of proportions of spatial domains identified by STAX across 238 slices, with different colors representing different types of domains. **b**, Cell type annotation (right) and domains (left) identified by STAX in the group 1 of the hierarchical clustering, where "_dt" indicates the domain type, and "_ct" indicates the cell type.

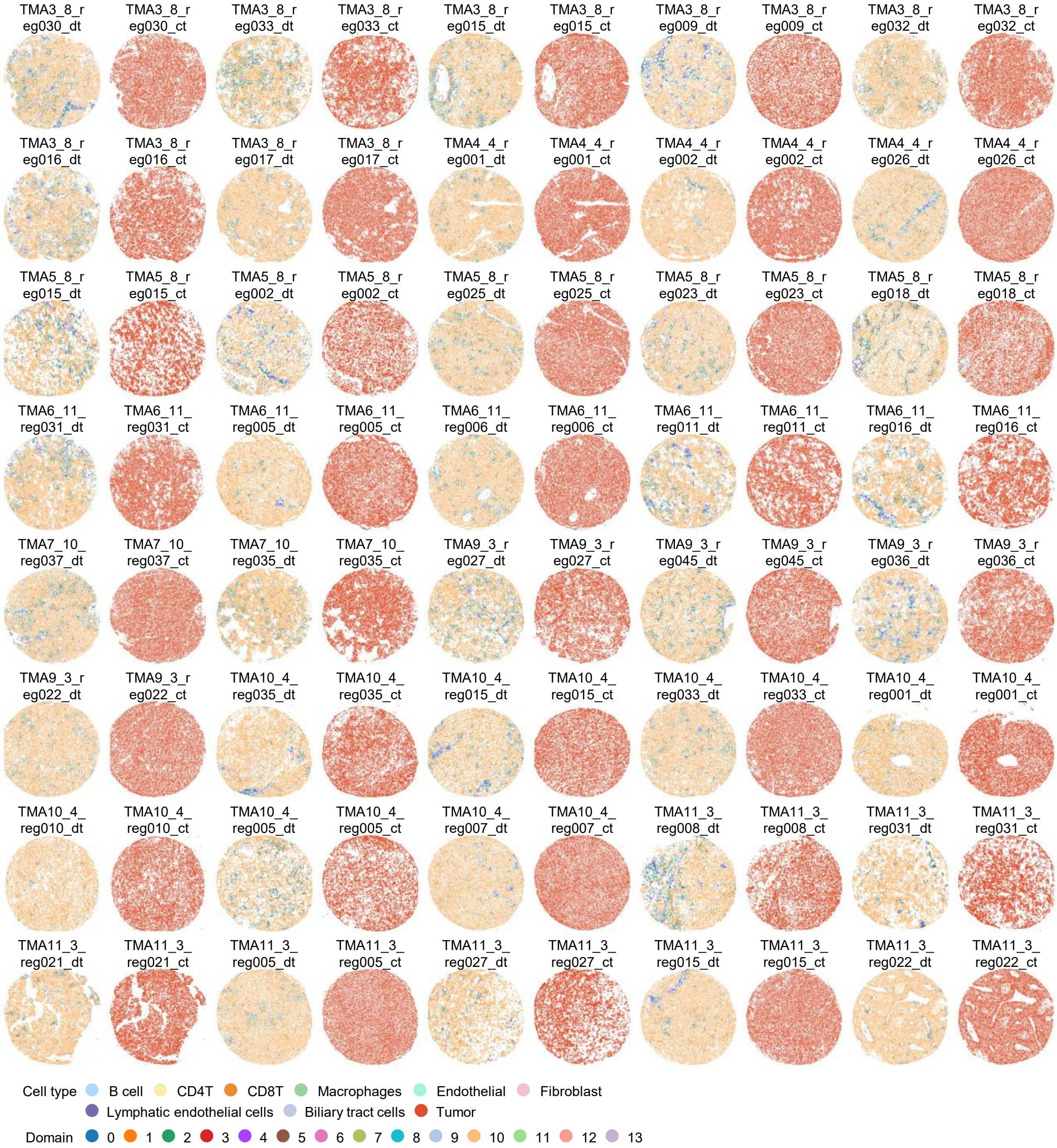


**Fig. S11.** Results of STAX on the spatial proteomics cohort study. Cell types annotation (right) and domains (left) identified by STAX in group 2 of the hierarchical clustering, where "_dt" indicates the domain type, and "_ct" indicates the cell type.

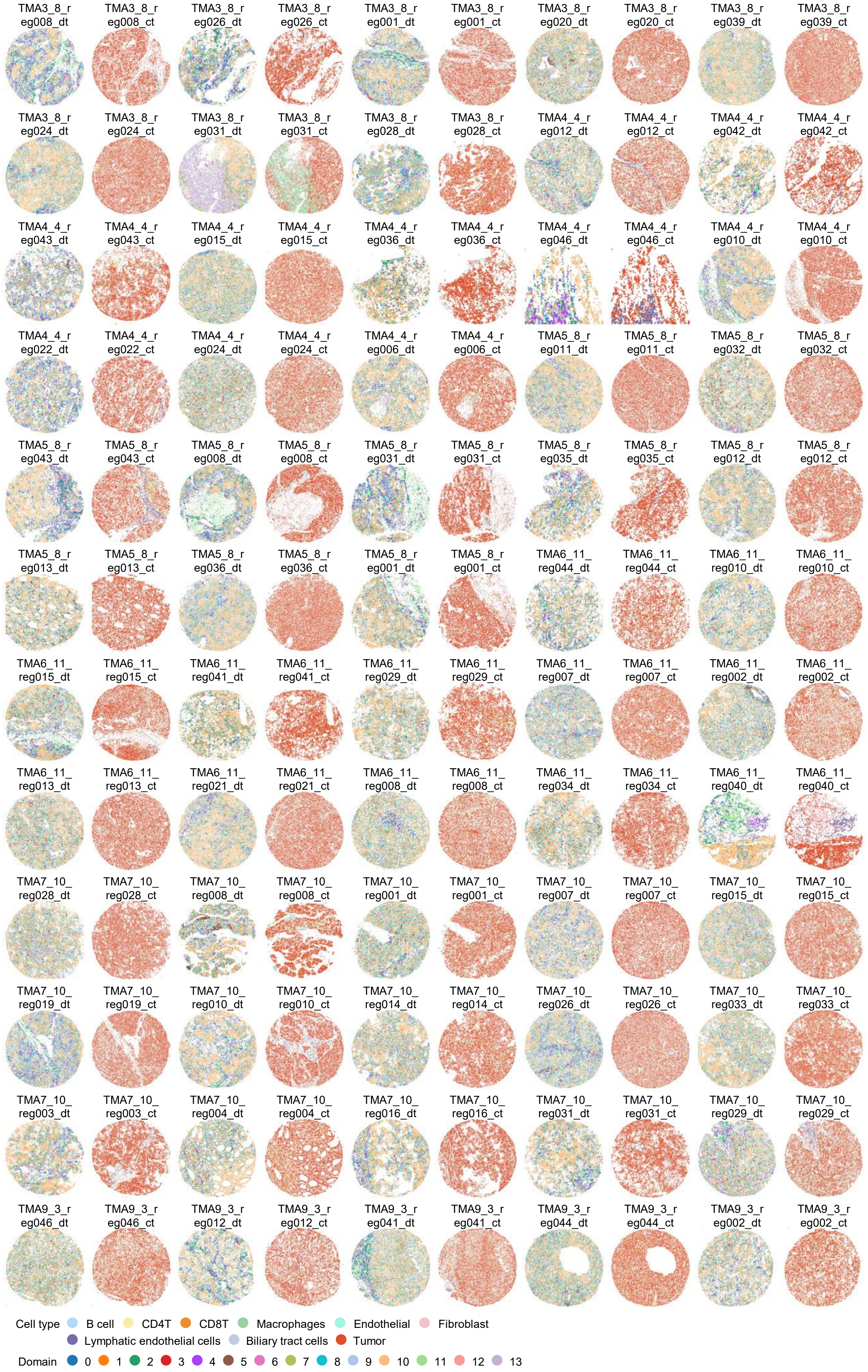

TMA3_8_reg008_dt
TMA3_8_reg008_ct
TMA3_8_reg026_dt
TMA3_8_reg026_ct
TMA3_8_reg001_dt
TMA3_8_reg001_ct
TMA3_8_reg020_dt
TMA3_8_reg020_ct
TMA3_8_reg039_dt
TMA3_8_reg039_ct
TMA3_8_reg024_dt
TMA3_8_reg024_ct
TMA3_8_reg031_dt
TMA3_8_reg031_ct
TMA3_8_reg028_dt
TMA3_8_reg028_ct
TMA4_4_reg012_dt
TMA4_4_reg012_ct
TMA4_4_reg042_dt
TMA4_4_reg042_ct
TMA4_4_reg043_dt
TMA4_4_reg043_ct
TMA4_4_reg015_dt
TMA4_4_reg015_ct
TMA4_4_reg036_dt
TMA4_4_reg036_ct
TMA4_4_reg046_dt
TMA4_4_reg046_ct
TMA4_4_reg010_dt
TMA4_4_reg010_ct
TMA4_4_reg022_dt
TMA4_4_reg022_ct
TMA4_4_reg024_dt
TMA4_4_reg024_ct
TMA4_4_reg006_dt
TMA4_4_reg006_ct
TMA5_8_reg011_dt
TMA5_8_reg011_ct
TMA5_8_reg032_dt
TMA5_8_reg032_ct
TMA5_8_reg043_dt
TMA5_8_reg043_ct
TMA5_8_reg008_dt
TMA5_8_reg008_ct
TMA5_8_reg031_dt
TMA5_8_reg031_ct
TMA5_8_reg035_dt
TMA5_8_reg035_ct
TMA5_8_reg012_dt
TMA5_8_reg012_ct
TMA5_8_reg013_dt
TMA5_8_reg013_ct
TMA5_8_reg036_dt
TMA5_8_reg036_ct
TMA5_8_reg001_dt
TMA5_8_reg001_ct
TMA6_11_reg044_dt
TMA6_11_reg044_ct
TMA6_11_reg010_dt
TMA6_11_reg010_ct
TMA6_11_reg015_dt
TMA6_11_reg015_ct
TMA6_11_reg041_dt
TMA6_11_reg041_ct
TMA6_11_reg029_dt
TMA6_11_reg029_ct
TMA6_11_reg007_dt
TMA6_11_reg007_ct
TMA6_11_reg002_dt
TMA6_11_reg002_ct
TMA6_11_reg013_dt
TMA6_11_reg013_ct
TMA6_11_reg021_dt
TMA6_11_reg021_ct
TMA6_11_reg008_dt
TMA6_11_reg008_ct
TMA6_11_reg034_dt
TMA6_11_reg034_ct
TMA6_11_reg040_dt
TMA6_11_reg040_ct
TMA7_10_reg028_dt
TMA7_10_reg028_ct
TMA7_10_reg008_dt
TMA7_10_reg008_ct
TMA7_10_reg001_dt
TMA7_10_reg001_ct
TMA7_10_reg007_dt
TMA7_10_reg007_ct
TMA7_10_reg015_dt
TMA7_10_reg015_ct
TMA7_10_reg019_dt
TMA7_10_reg019_ct
TMA7_10_reg010_dt
TMA7_10_reg010_ct
TMA7_10_reg014_dt
TMA7_10_reg014_ct
TMA7_10_reg026_dt
TMA7_10_reg026_ct
TMA7_10_reg033_dt
TMA7_10_reg033_ct
TMA7_10_reg003_dt
TMA7_10_reg003_ct
TMA7_10_reg004_dt
TMA7_10_reg004_ct
TMA7_10_reg016_dt
TMA7_10_reg016_ct
TMA7_10_reg031_dt
TMA7_10_reg031_ct
TMA7_10_reg029_dt
TMA7_10_reg029_ct
TMA9_3_reg046_dt
TMA9_3_reg046_ct
TMA9_3_reg012_dt
TMA9_3_reg012_ct
TMA9_3_reg041_dt
TMA9_3_reg041_ct
TMA9_3_reg044_dt
TMA9_3_reg044_ct
TMA9_3_reg002_dt
TMA9_3_reg002_ct
Cell type
B cell
CD4T
CD8T
Macrophages
Endothelial
Fibroblast
Lymphatic endothelial cells
Biliary tract cells
Tumor
Domain
0 1 2 3 4 5 6 7 8 9 10 11 12 13

**Fig. S12.** Results of STAX on the spatial proteomics cohort study. Cell types annotation (right) and domains (left) identified by STAX in group 3 of the hierarchical clustering, where "_dt" indicates the domain type, and "_ct" indicates the cell type.

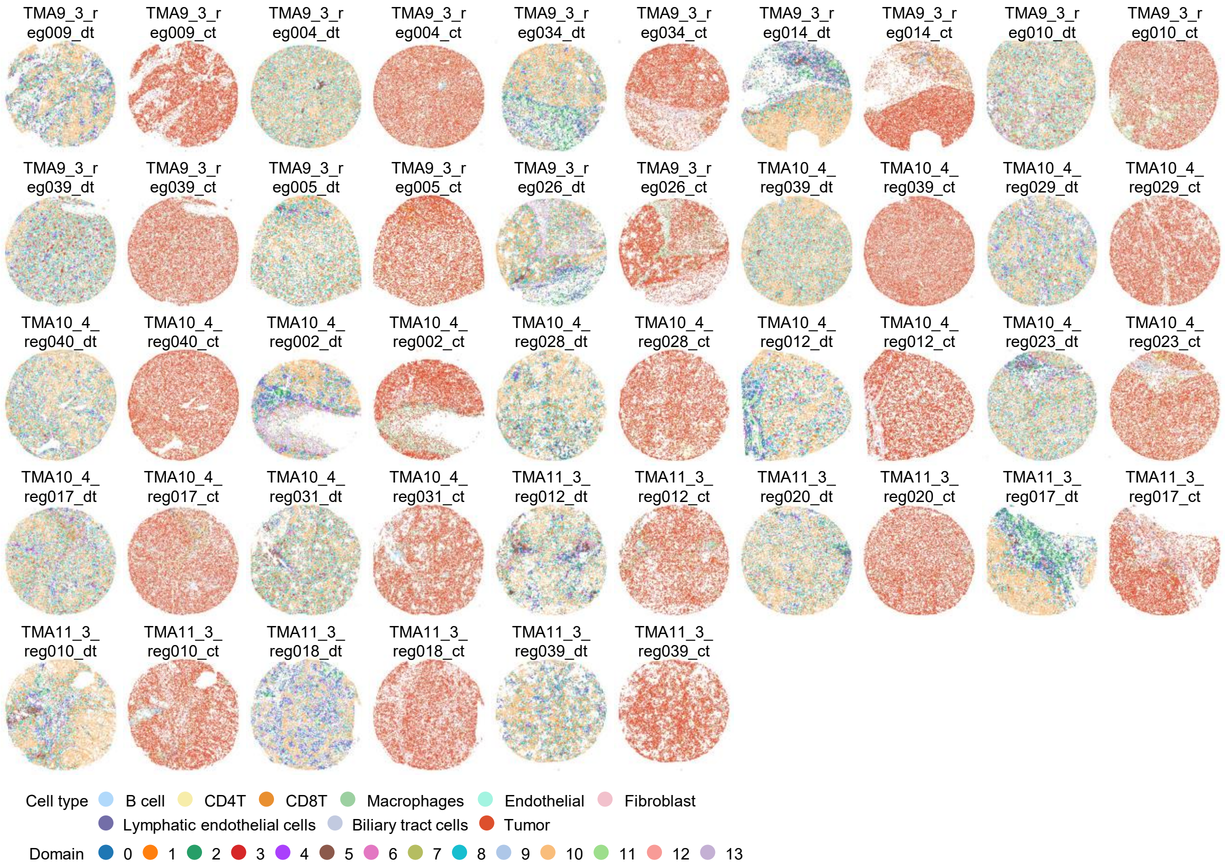


**Fig. S13 (Fig. S12 Contiuing).** Results of STAX on the spatial proteomics cohort study. Cell types annotation (right) and domains (left) identified by STAX in group 3 of the hierarchical clustering, where "_dt" indicates the domain type, and "_ct" indicates the cell type.

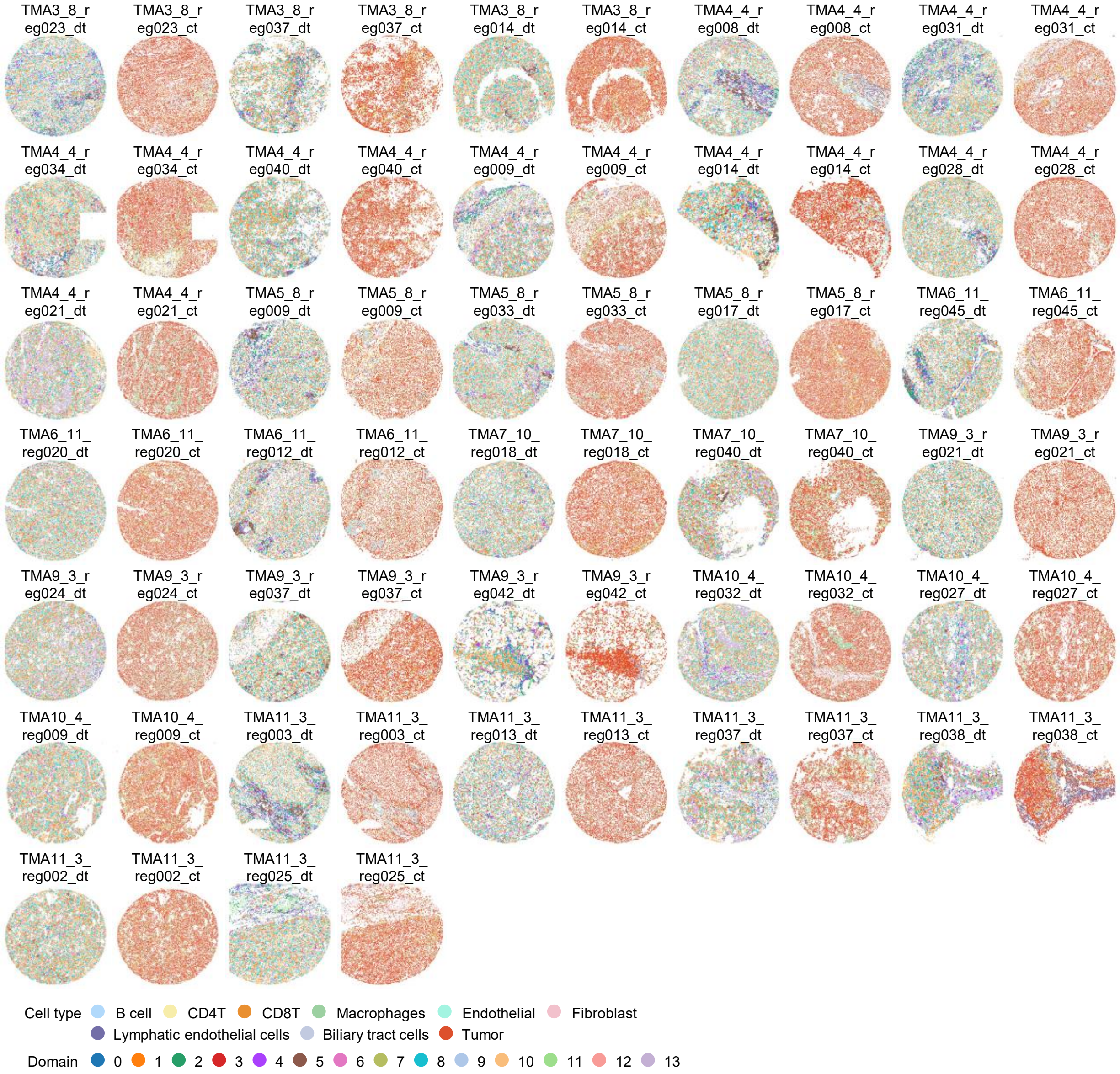


**Fig. S14.** Results of STAX on the spatial proteomics cohort study. Cell types annotation (right) and domains (left) identified by STAX in group 4 of the hierarchical clustering, where "_dt" indicates the domain type, and "_ct" indicates the cell type.

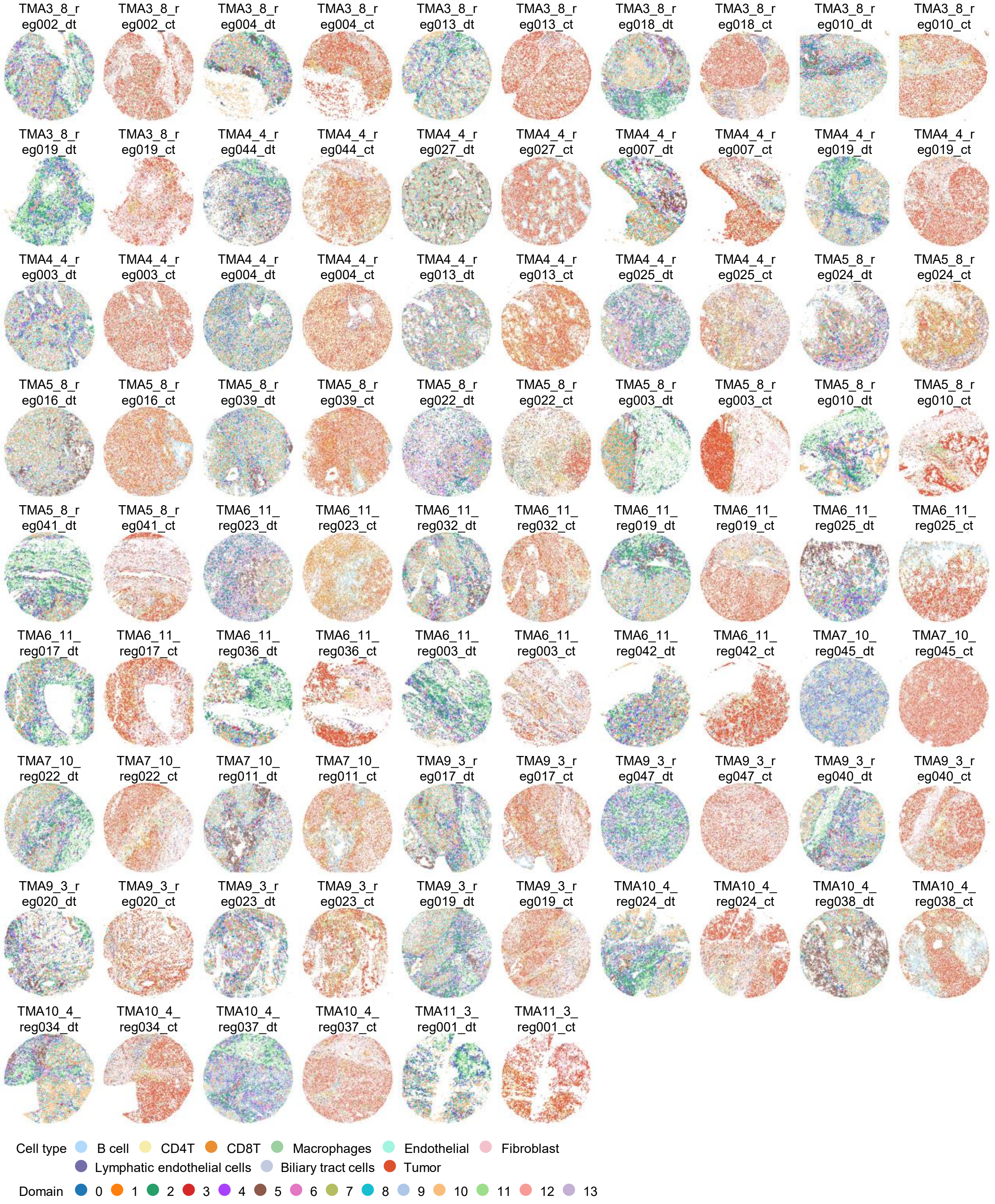


**Fig. S15.** Results of STAX on the spatial proteomics cohort study. Cell types annotation (right) and domains (left) identified by STAX in group 5 of the hierarchical clustering, where "_dt" indicates the domain type, and "_ct" indicates the cell type.

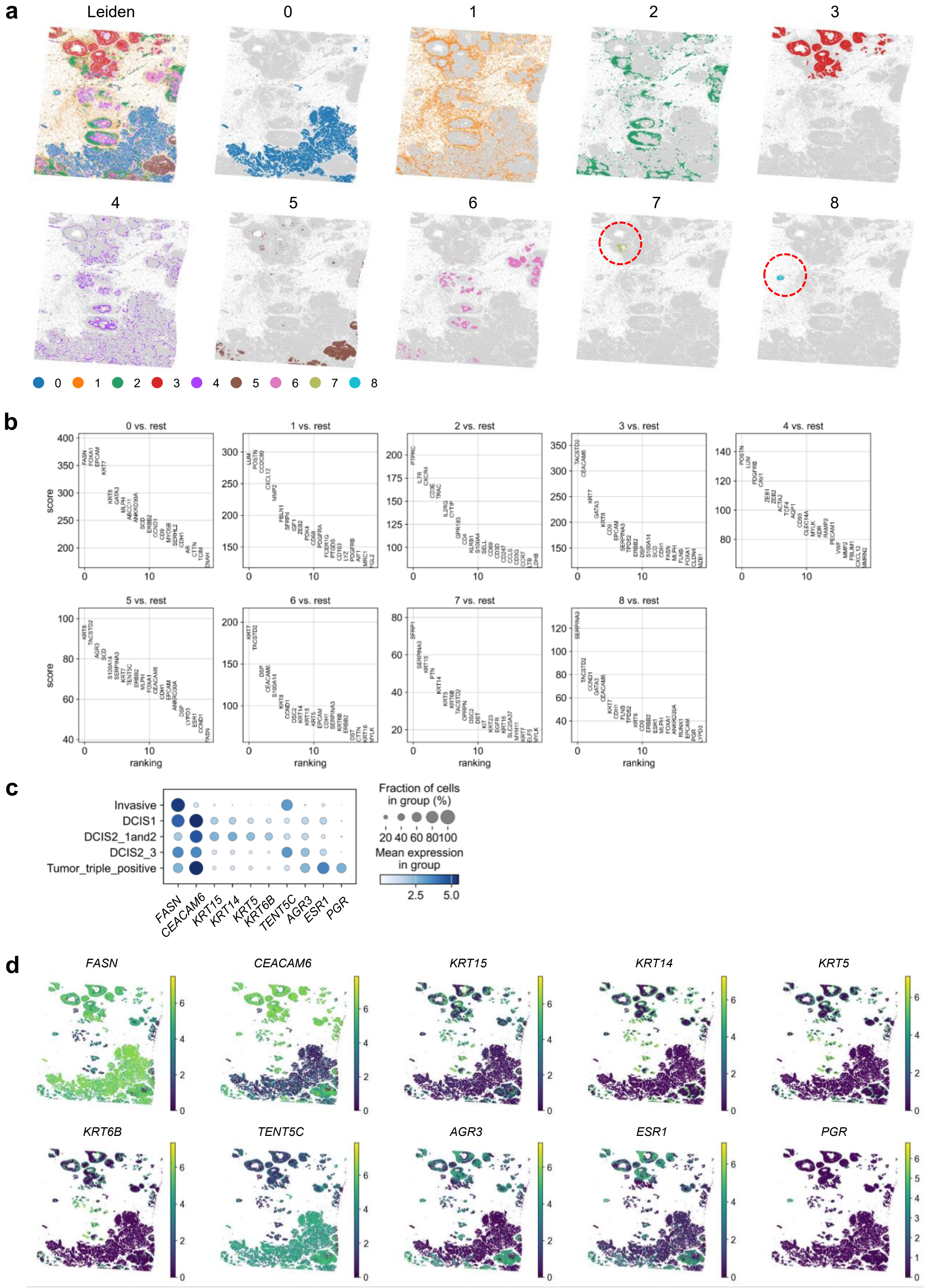


**Fig. S16.** Results of STAX on the breast cancer Xenium dataset. **a**, Spatial domains identified by Leiden based on the STAX cell embeddings. **b**, Differentially expressed genes identified by Scanpy for each domain. **c**, Dot plot of top10 differentially expressed genes for the tumor domain. **d**, Spatial map of the ten tumor markers.

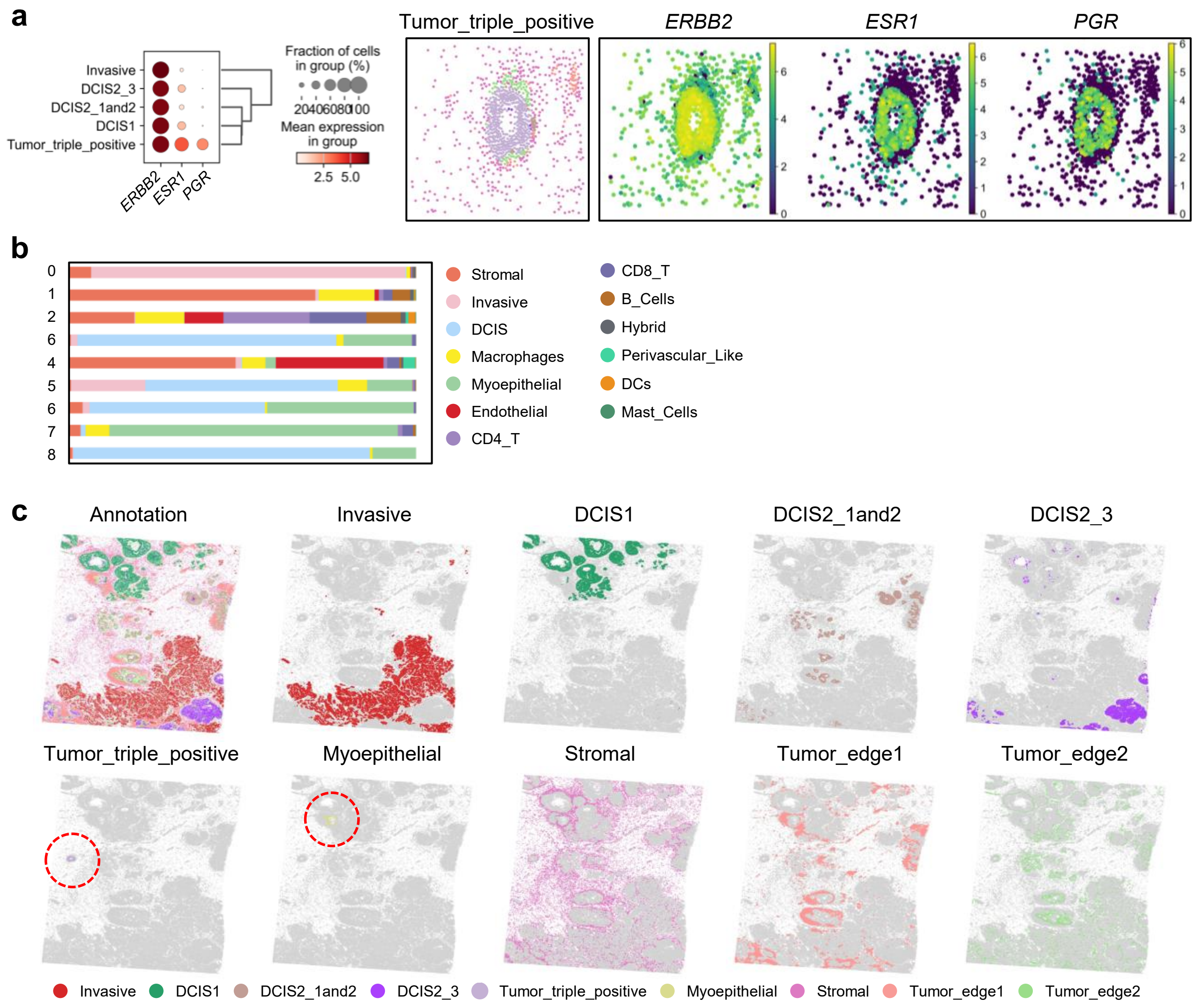


**Fig. S17.** Results of STAX on the breast cancer Xenium dataset. **a**, Expression profiles of *ERBB2*, *ESR1*, and *PGR* across various spatial domains identified by STAX, including Invasive, DCIS1, DCIS2_1 and 2, DCIS2_3, and Tumor_triple_positive. **b**, Cell proportions within the spatial domains. **c**, Spatial visualization of the domains identified by STAX.

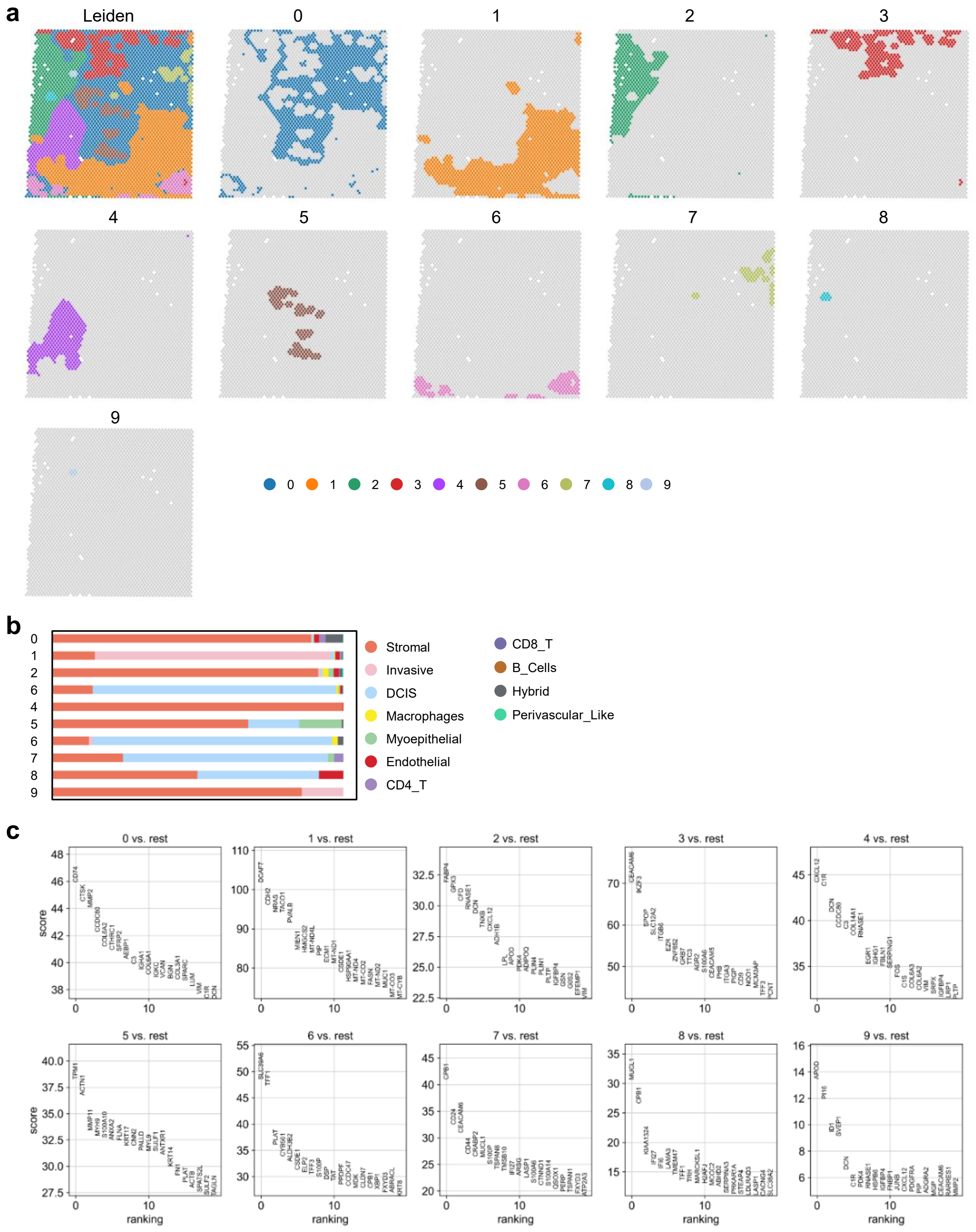

**Fig. S18.** Results of STAX on the breast cancer Visium dataset. **a**, Spatial domains identified by Leiden based on the STAX spot embeddings. **b**, Cell proportion in all domains identified by STAX. **c**, Differentially expressed genes identified by Scanpy for each domain.

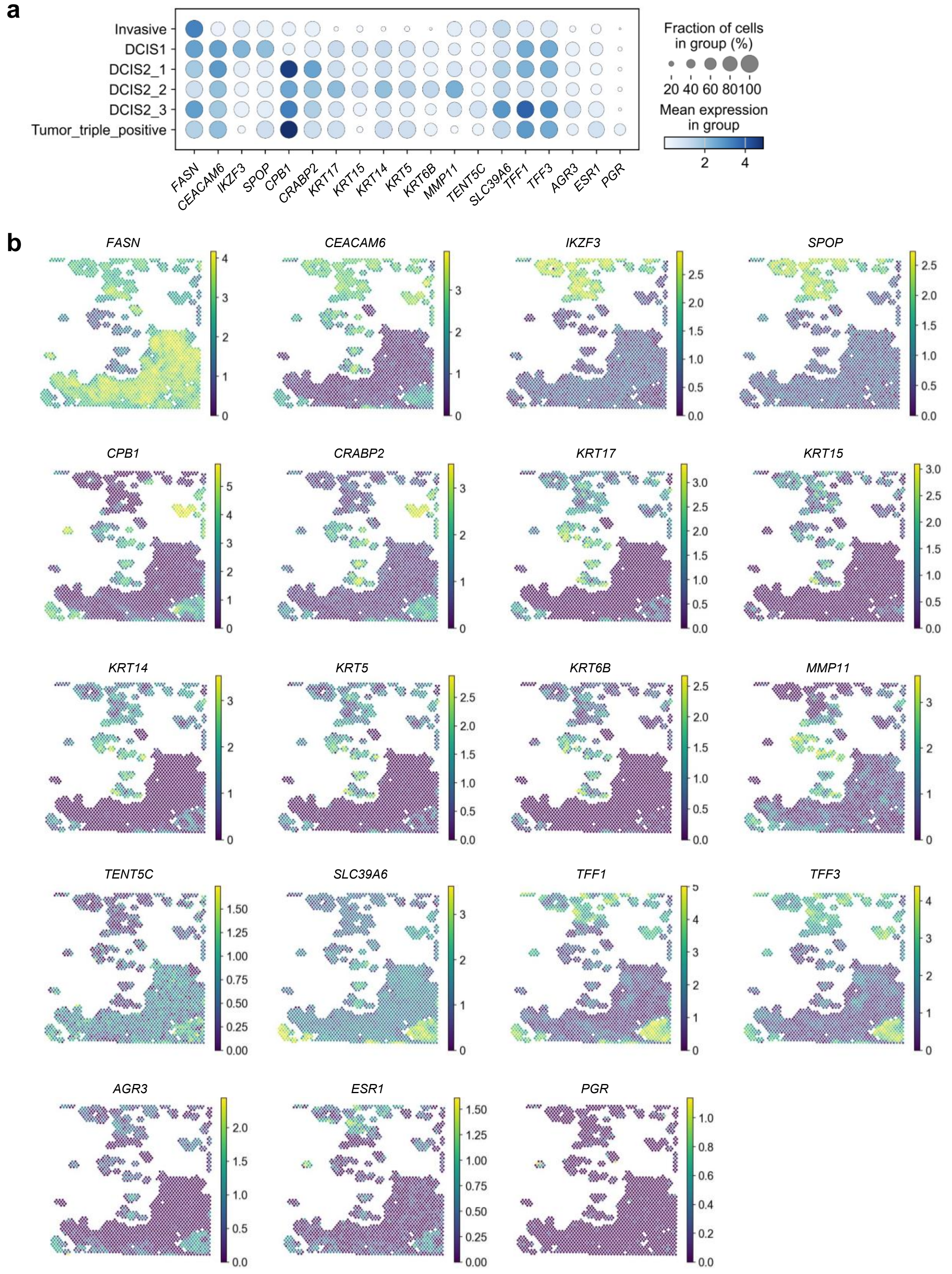


**Fig. S19.** Results of STAX on the breast cancer Visium dataset. **a**, Dot plot of several differentially expressed genes for the tumor domain. **b**, Spatial map plot of several tumor markers.

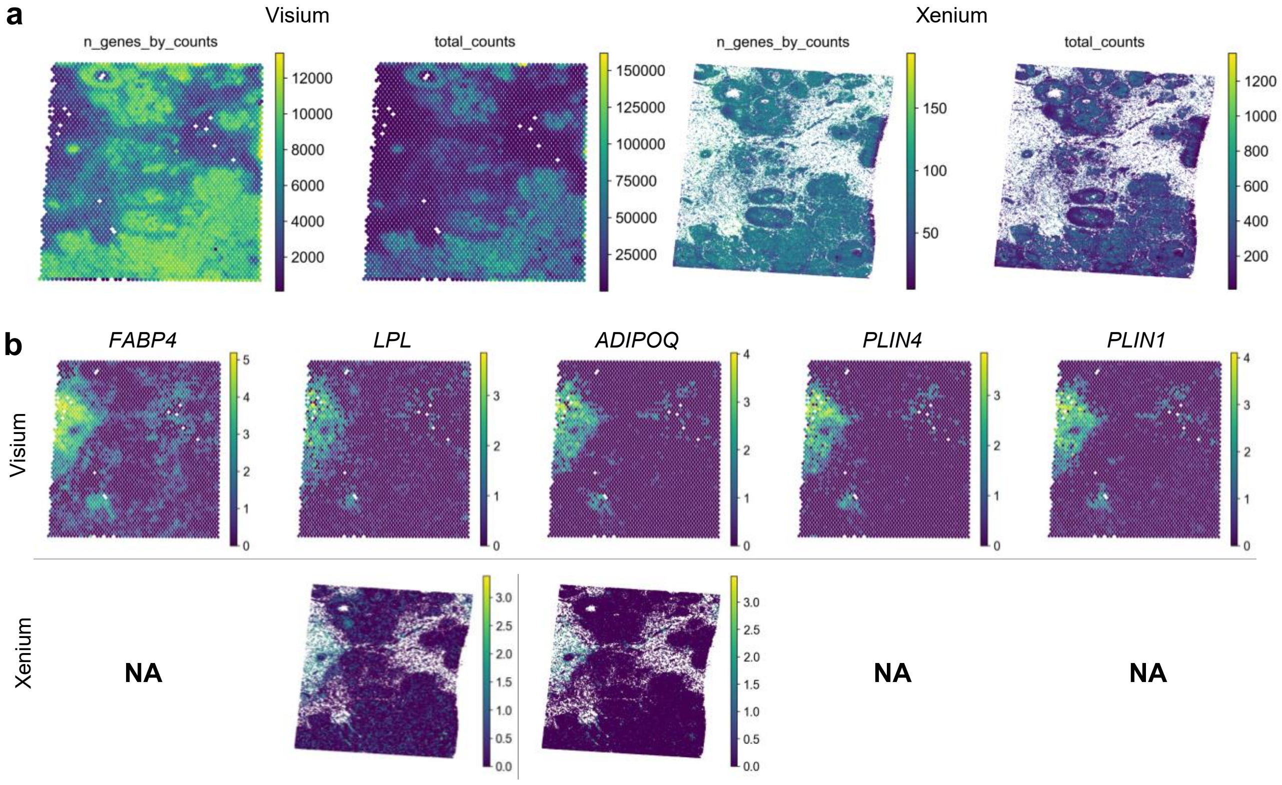


**Fig. S20.** Results of STAX on the breast cancer Visium dataset. **a**, Heatmaps displaying the number of genes with at least one detected transcript (n_genes_by_counts) and the aggregate sum of all UMIs across all genes per cell or capture spot (total_counts). Metrics were calculated using the Scanpy framework. **b**, Spatial map of adipose marker genes the in Visium and Xenium datasets, respectively.

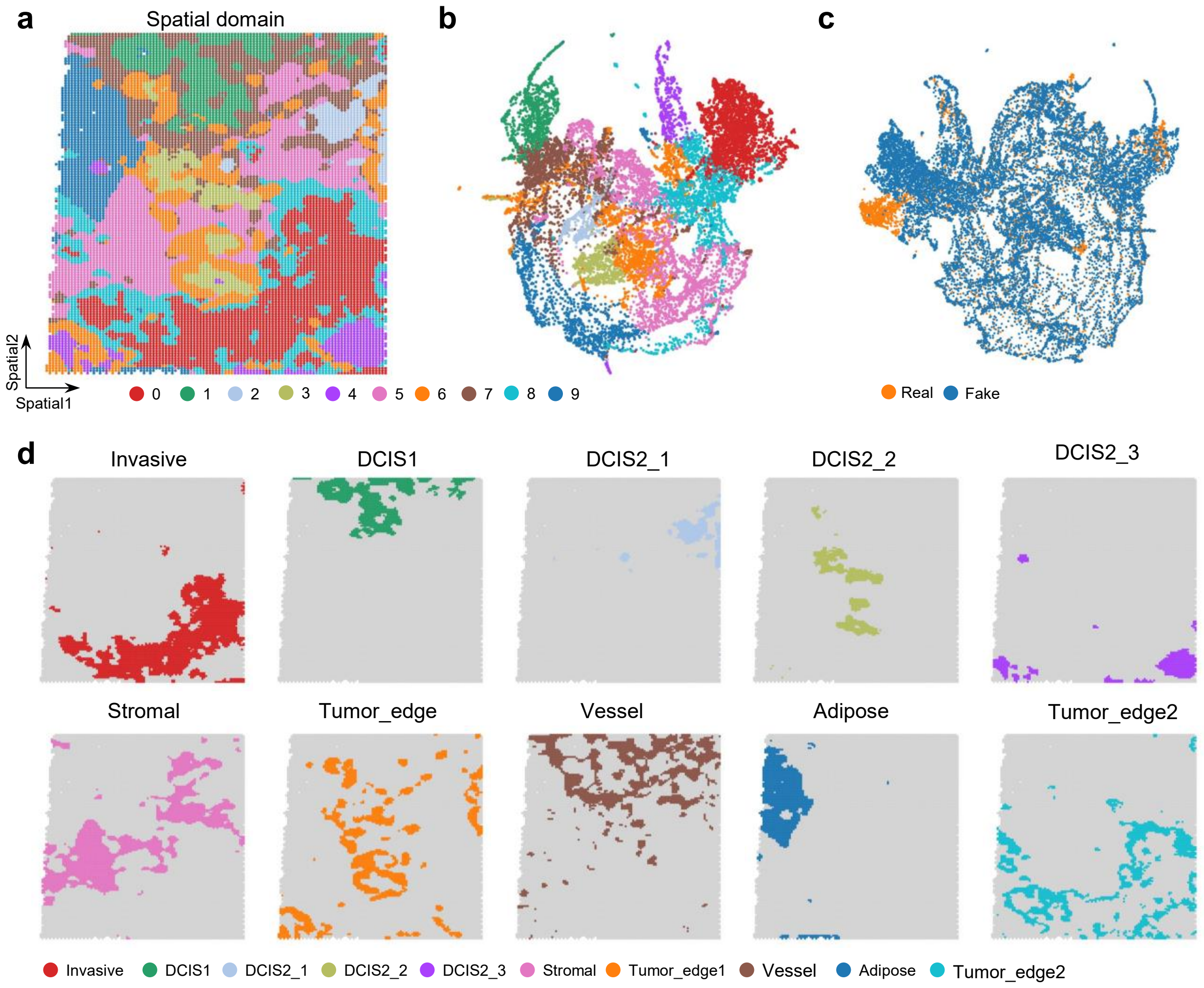


**Fig. S21.** Spot completion by STAX on the breast cancer Visium dataset. **a**, Spatial domains identified by STAX based on the completed Visium dataset. **b**, UMAP plot illustrating the domains. **c**, UMAP plot illustrating the real and fake spots. **d**, Spatial map of each spatial domain identified by STAX based on the completed Visium dataset.

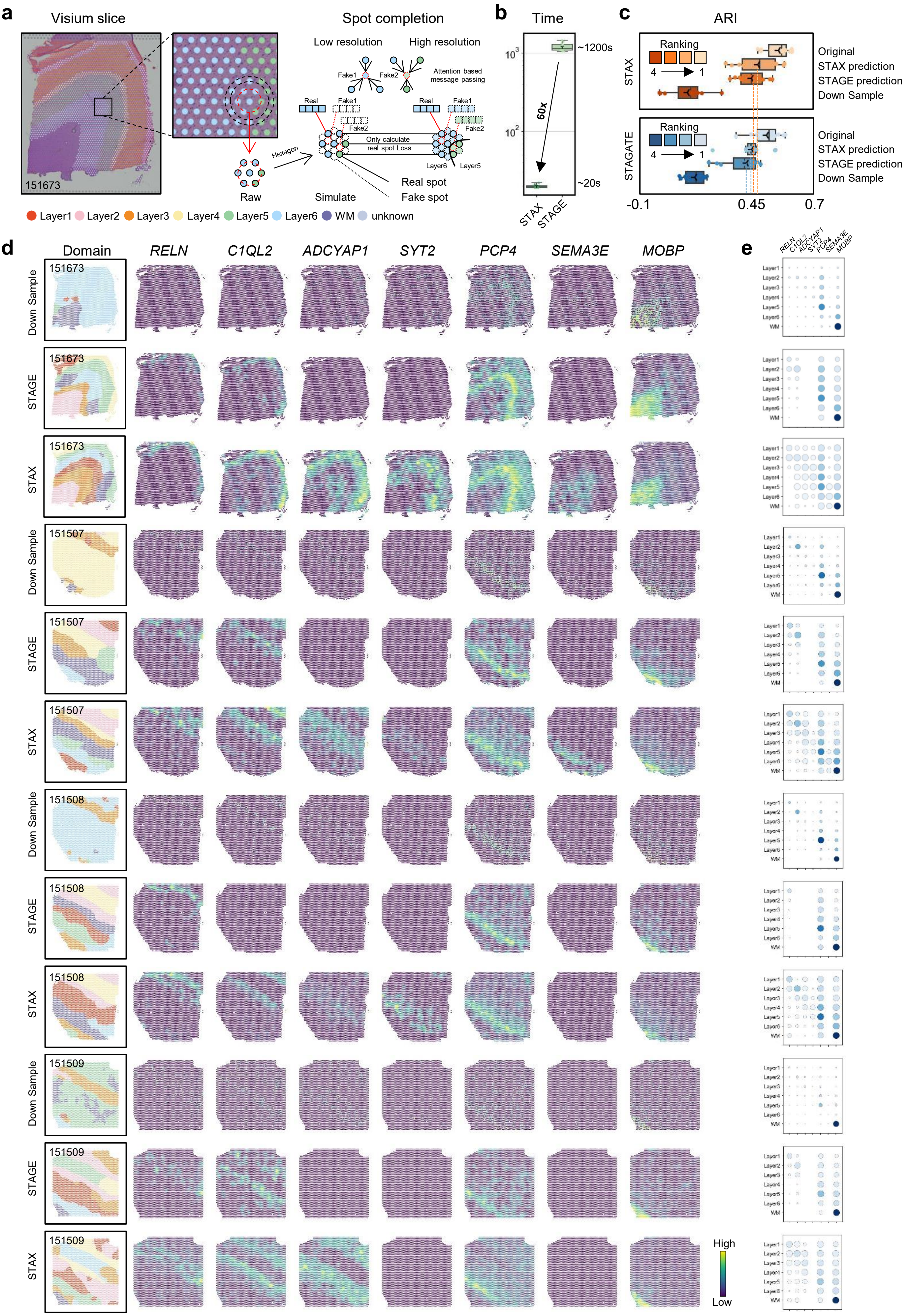

a
Visium slice
151673
Spot completion
Low resolution
High resolution
Fake1
Fake2
Attention based message passing
Real
Fake1
Fake2
Only calculate real spot Loss
Layer6
Layer5
Hexagon
Raw
Simulate
Real spot
Fake spot
Layer1 Layer2 Layer3 Layer4 Layer5 Layer6 WM unknown
b
Time
~1200s
60x
~20s
STAX
STAGE
c
ARI
Ranking
4 1
STAX
Original
STAX prediction
STAGE prediction
Down Sample
STAGATE
-0.1
0.45
0.7
d
Domain
RELN
C1QL2
ADCYAP1
SYT2
PCP4
SEMA3E
MOBP
Down Sample
STAGE
STAX
151673
151507
151508
151509
High
Low
e
Layer1
Layer2
Layer3
Layer4
Layer5
Layer6
WM

**Fig. S22.** Spot completion by STAX on the DLPFC Visium dataset. **a**, Schematic representation of the DLPFC dataset and illustration of spot completion by STAX. **b**, Comparison of running time between STAX and STAGE. **c**, Spatial domain identification by using STAX and STAGATE on the original, down-sampled, STAX spot-completion and STAGE spot-completion slices, respectively. **d**, Spatial expression of marker genes for down-sampled, STAGE-completed, and STAX-completed data. **e**, Dot plot of marker gene expression in down-sampled, STAGE-completed data, and STAX-completed data.

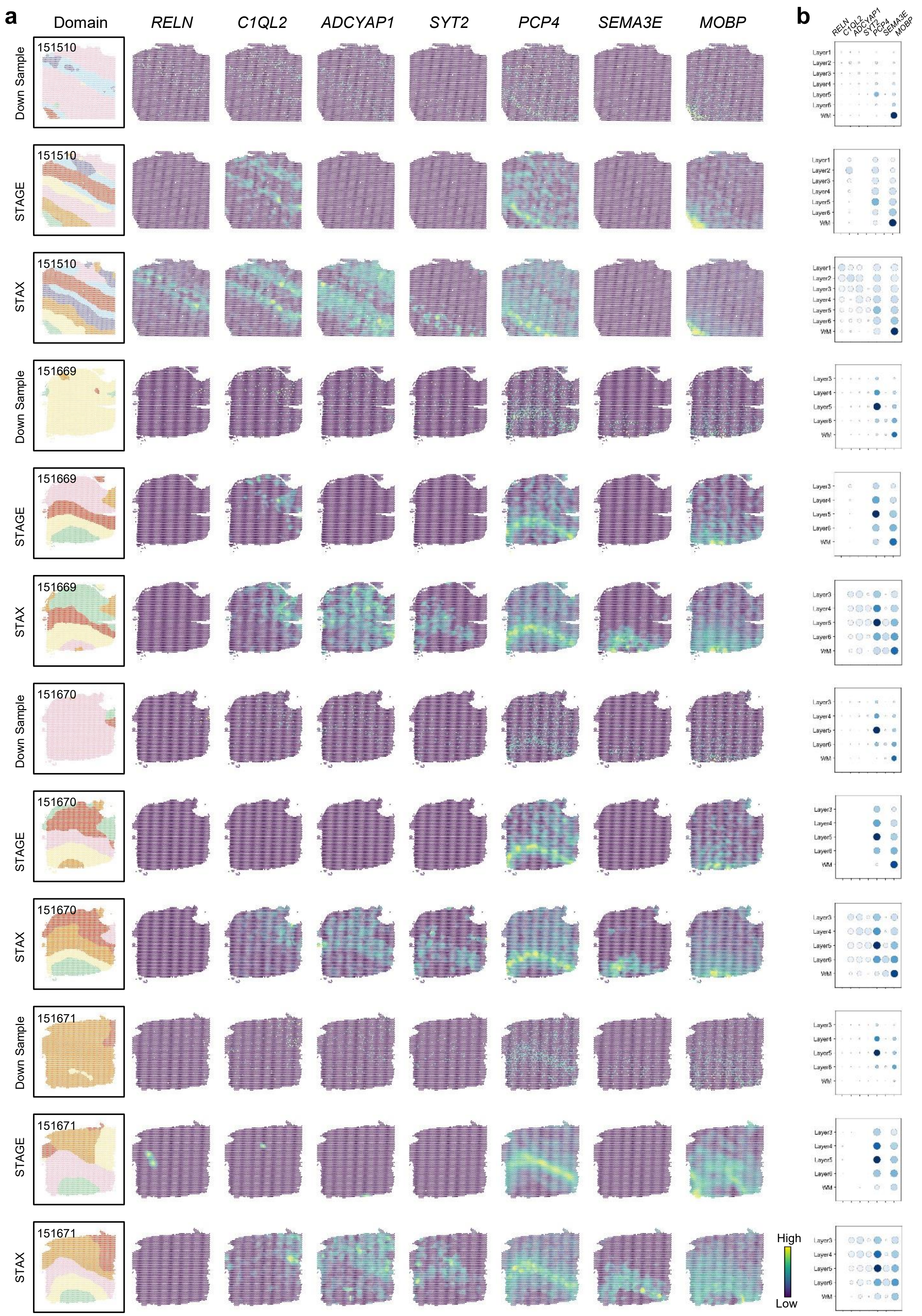


**Fig. S23.** Spot completion by STAX on the DLPFC Visium dataset. **a**, Spatial expression of marker genes on the raw, STAGE-imputed, and STAX-imputed data. **b**, Dot plot of marker gene expression in down-sampled, STAGE-imputed, and STAX-imputed data, respectively.

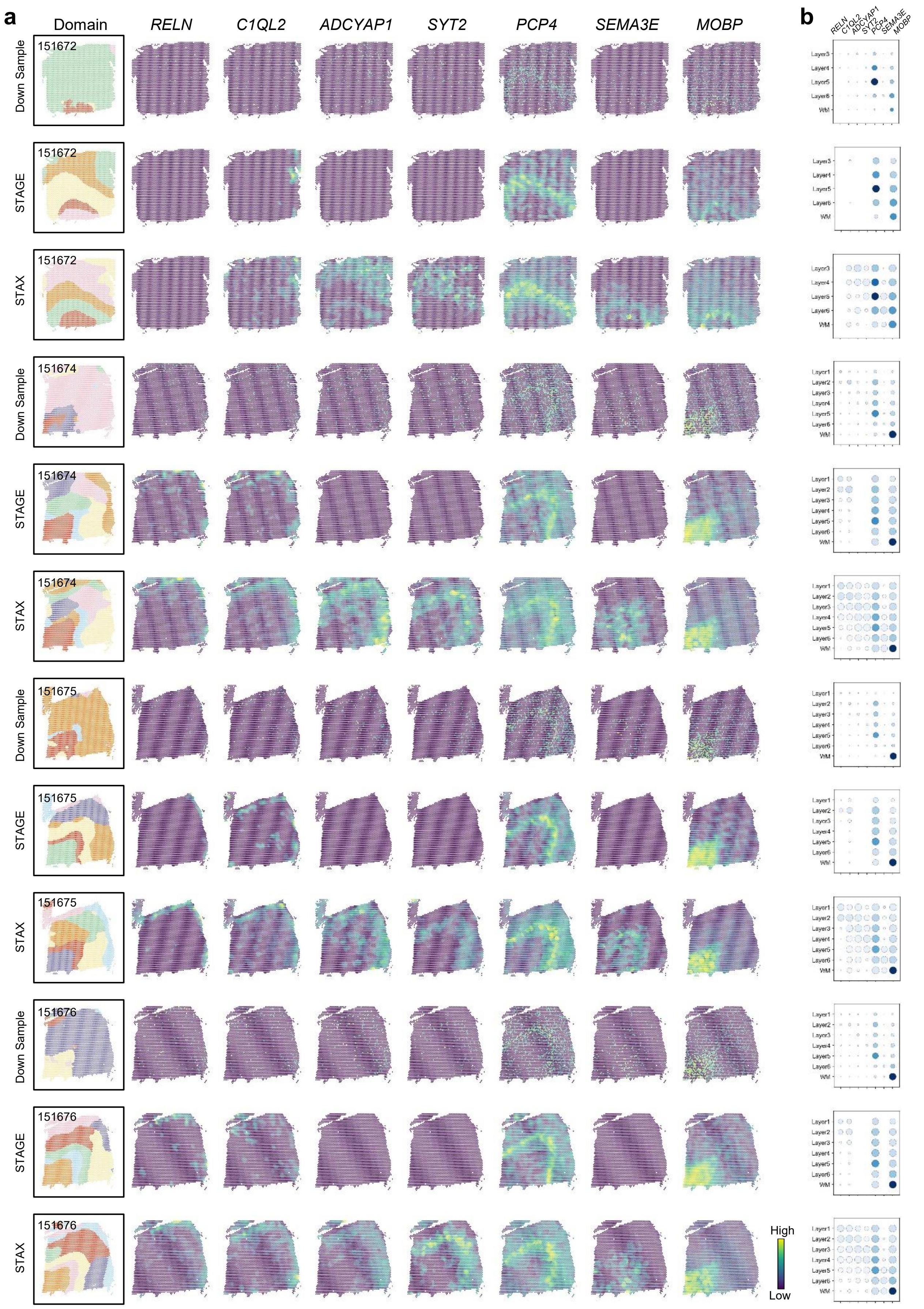


**Fig. S24.** Spot completion by STAX on the DLPFC Visium dataset. **a**, Spatial expression of marker genes on the raw, STAGE-imputed, and STAX-imputed data. **b**, Dot plot of marker gene expression on down-sampled, STAGE-imputed, and STAX-imputed data, respectively.

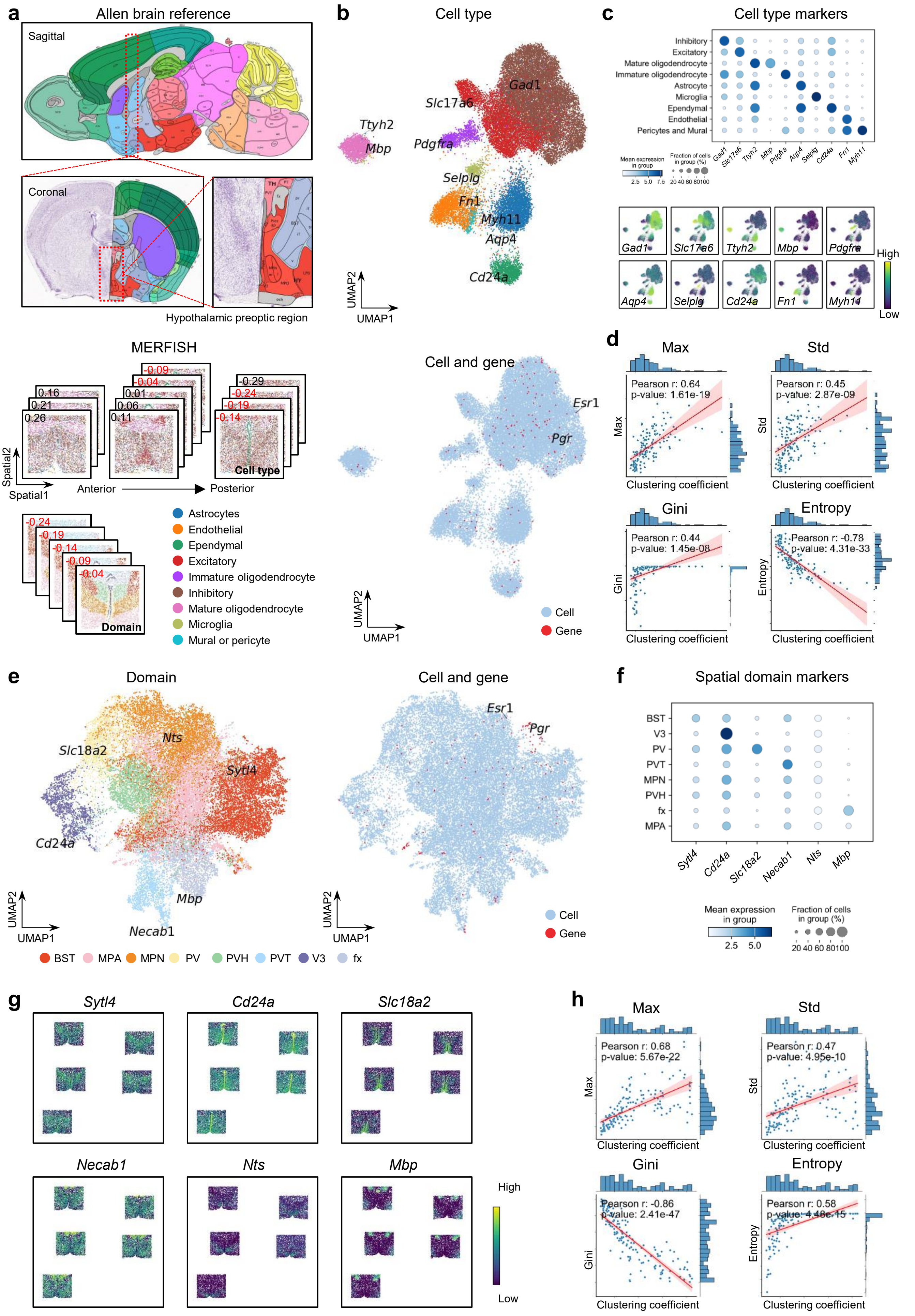
a
Allen brain reference
Sagittal
Coronal
Hypothalamic preoptic region
MERFISH
Cell type
Domain
Spatial2
Spatial1
Anterior
Posterior
Astrocytes
Endothelial
Ependymal
Excitatory
Immature oligodendrocyte
Inhibitory
Mature oligodendrocyte
Microglia
Mural or pericyte
b
Cell type
Gad1
Slc17a6
Ttyh2
Mbp
Pdgfra
Selplg
Fn1
Myh11
Aqp4
Cd24a
UMAP2
UMAP1
Cell and gene
Esr1
Pgr
Cell
Gene
c
Cell type markers
Inhibitory
Excitatory
Mature oligodendrocyte
Immature oligodendrocyte
Astrocyte
Microglia
Ependymal
Endothelial
Pericytes and Mural
Mean expression in group
Fraction of cells in group (%)
High
Low
d
Max
Std
Gini
Entropy
Pearson r: 0.64
p-value: 1.61e-19
Pearson r: 0.45
p-value: 2.87e-09
Pearson r: 0.44
p-value: 1.45e-08
Pearson r: -0.78
p-value: 4.31e-33
Clustering coefficient
e
Domain
Nts
Slc18a2
Sytl4
Cd24a
Mbp
Necab1
BST
MPA
MPN
PV
PVH
PVT
V3
fx
f
Spatial domain markers
g
Sytl4
Cd24a
Slc18a2
Necab1
Nts
Mbp
High
Low
h
Max
Std
Gini
Entropy
Pearson r: 0.68
p-value: 5.67e-22
Pearson r: 0.47
p-value: 4.95e-10
Pearson r: -0.86
p-value: 2.41e-47
Pearson r: 0.58
p-value: 4.48e-15
Clustering coefficient

**Fig. S25.** Cell and gene co-embeddings by STAX on the MERFISH dataset. **a**, Details of the MERFISH dataset including cell-type and domain annotations. **b**, UMAP plot of the cell and gene co-embeddings under setting 1 generated by STAX at the cell level. **c**, Dot plot and heatmap displaying the marker genes identified in the original literature. **d**, The correlation between the clustering coefficient and four SIMBA metrics including MAX, Std, Gini, and Entropy under setting 1. **e**, UMAP plot of cell and gene co-embeddings under setting 2 generated by STAX at the domain level. **f**, Dot plot highlighting the marker genes identified in the original literature. **g**, Heatmap showing the expression patterns of marker genes identified in the original literature. **h**, The correlation between the clustering coefficient and four SIMBA metrics including MAX, Std, Gini, and Entropy under setting 2.

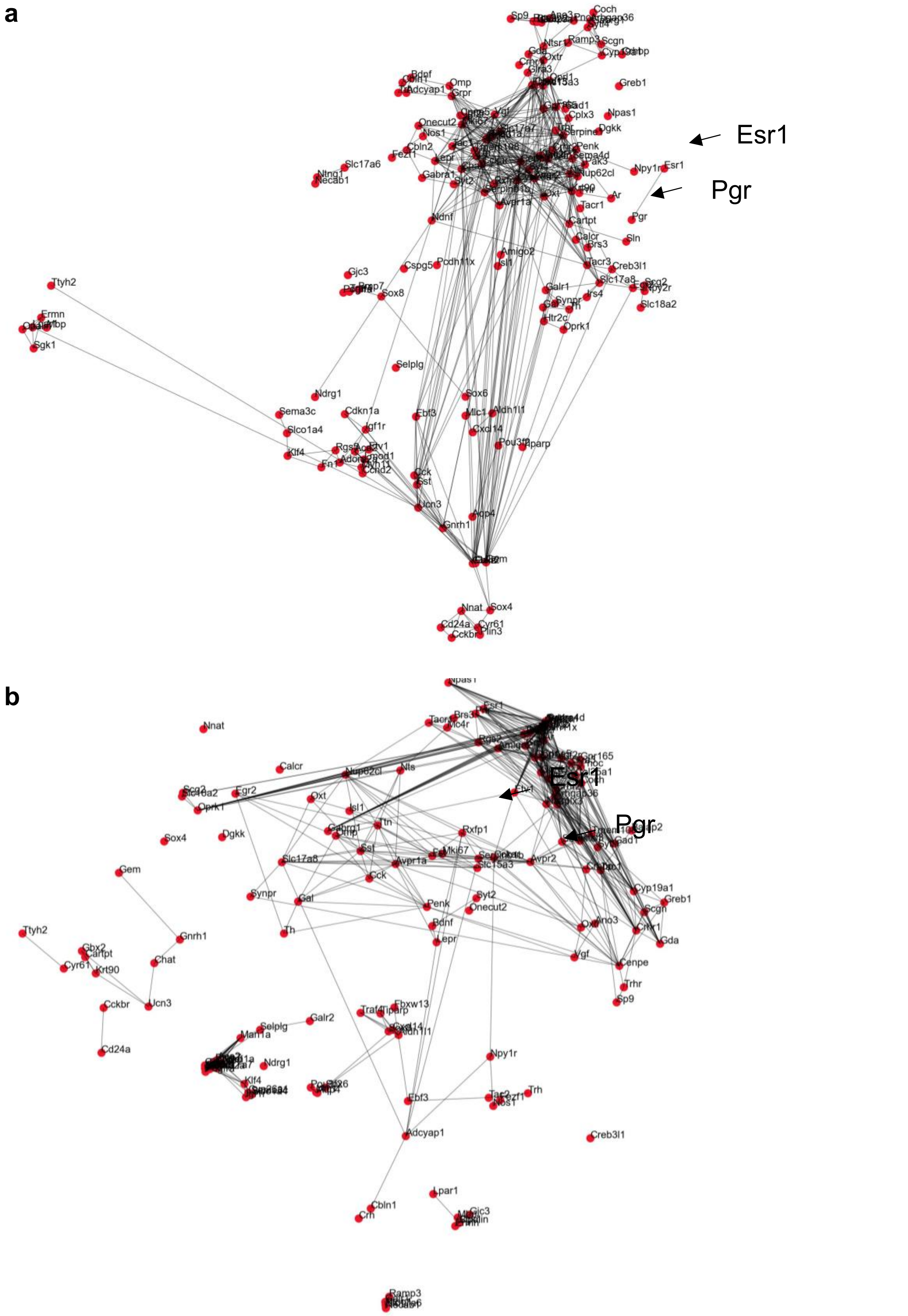


**Fig. S26.** Gene-gene network by STAX on the MERFISH dataset. **a**. A gene-gene network derived from the cell-gene co-embedding space of the MERFISH dataset under setting 1 (neighbor graph with self-loops only). This network highlights potential co-expression among genes, providing insights into gene-gene relationships within a cell-centric context. **b**. A gene-gene network derived from the cell-gene co-embedding space of the MERFISH dataset under setting 2 (neighbor graph with a radius parameter of 50). This network incorporates spatial context, revealing gene-gene interactions that are influenced by spatial proximity and domain-specific patterns.

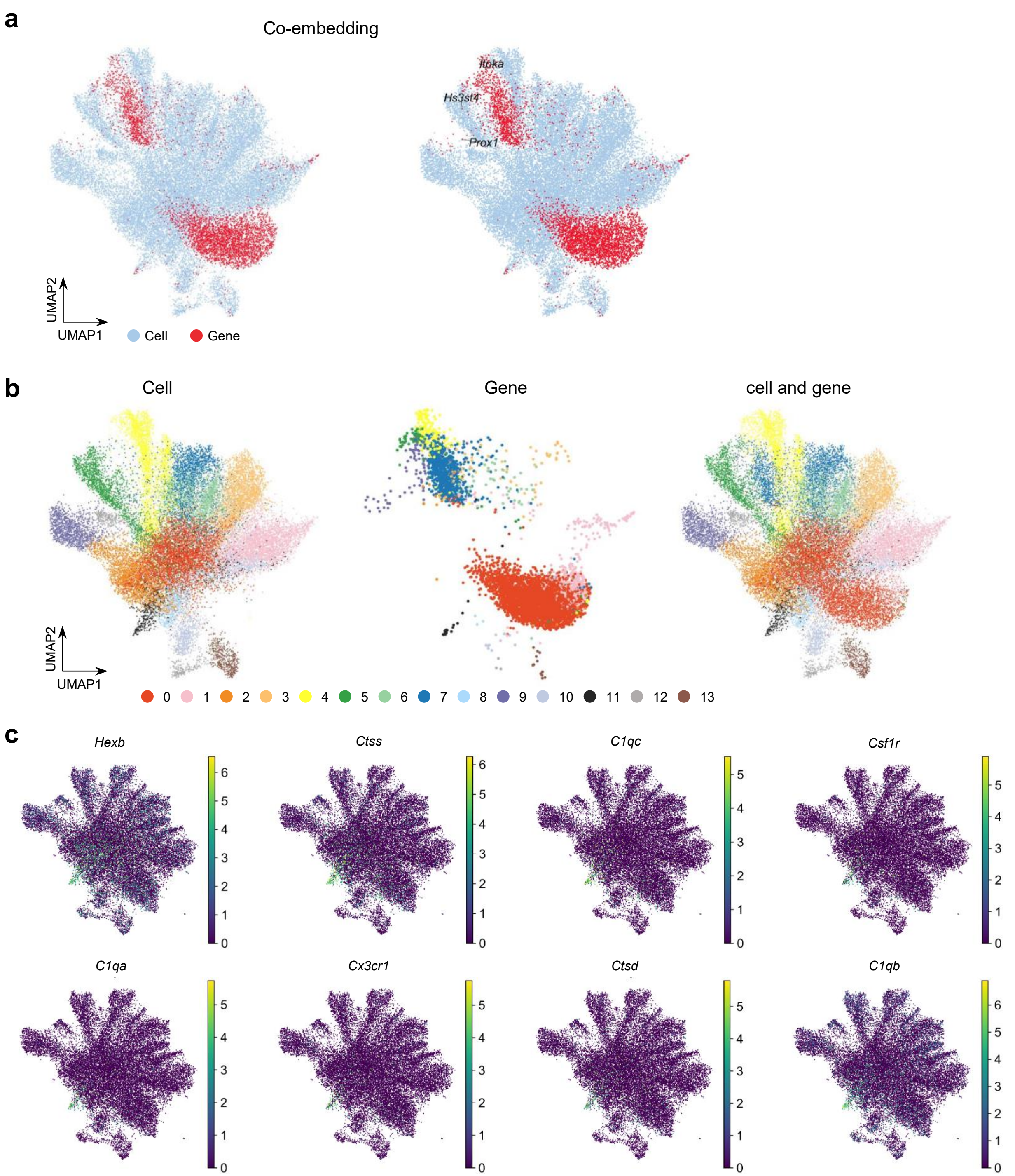


**Fig. S27.** Cell and gene co-embeddings by STAX on the hippocampus dataset. **a**, Cell and gene co-embeddings of Alzheimer's disease and healthy tissue slices. **b**, Transfer of domain annotation (Louvain) from cell to gene in the co-embedding space. **c**, Spatial map of marker genes in Alzheimer's disease and healthy tissue slices.

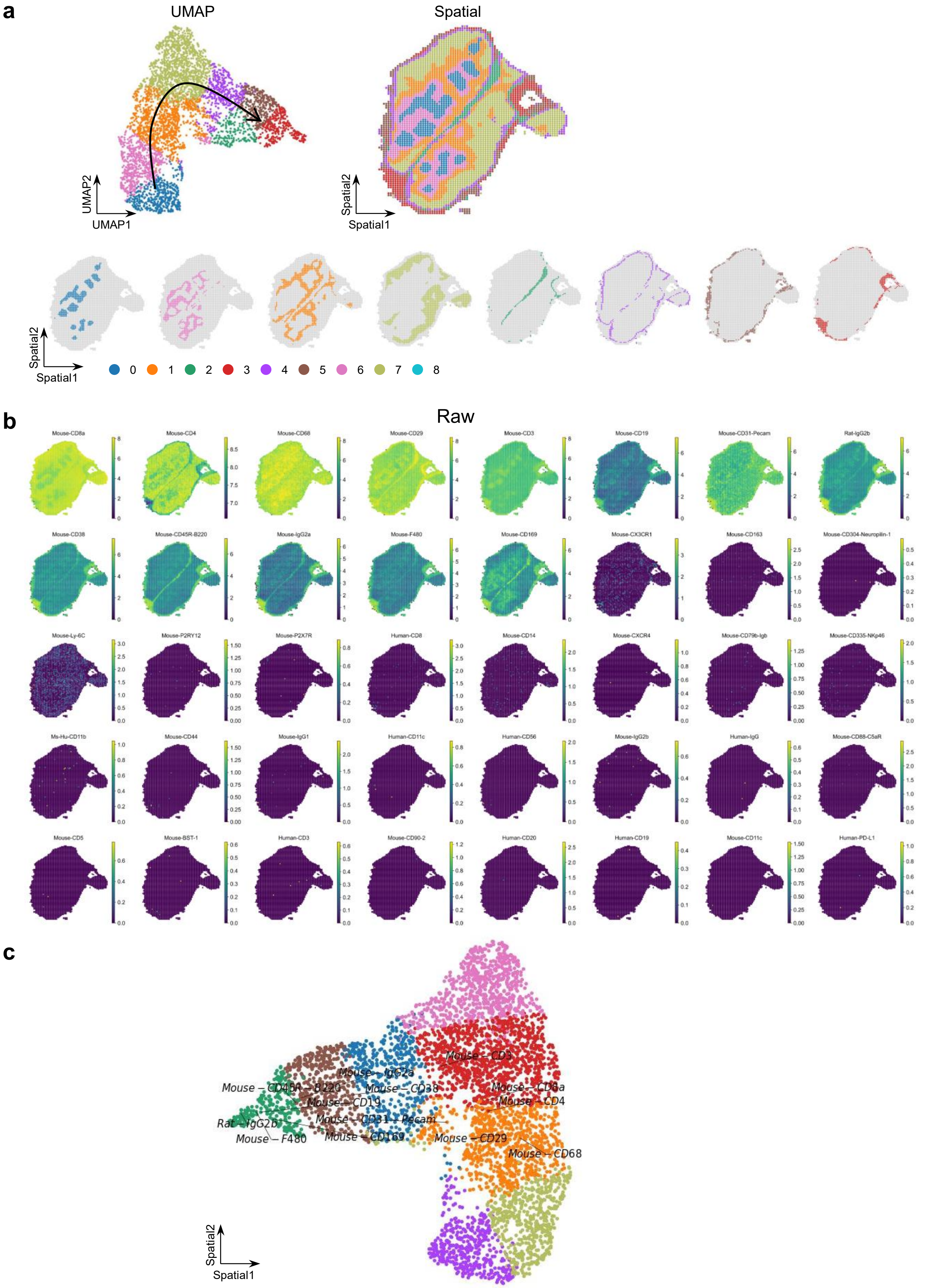


**Fig. S28.** Spatial analysis by STAX on the mouse thymus spatial proteomics dataset. **a**, Spatial domains identified by STAX showing the distinct regions. **b**, Spatial distribution of all protein expression in the dataset. **c**, Cell and gene co-embeddings revealing the relationships between cells and genes in a shared low-dimensional space.

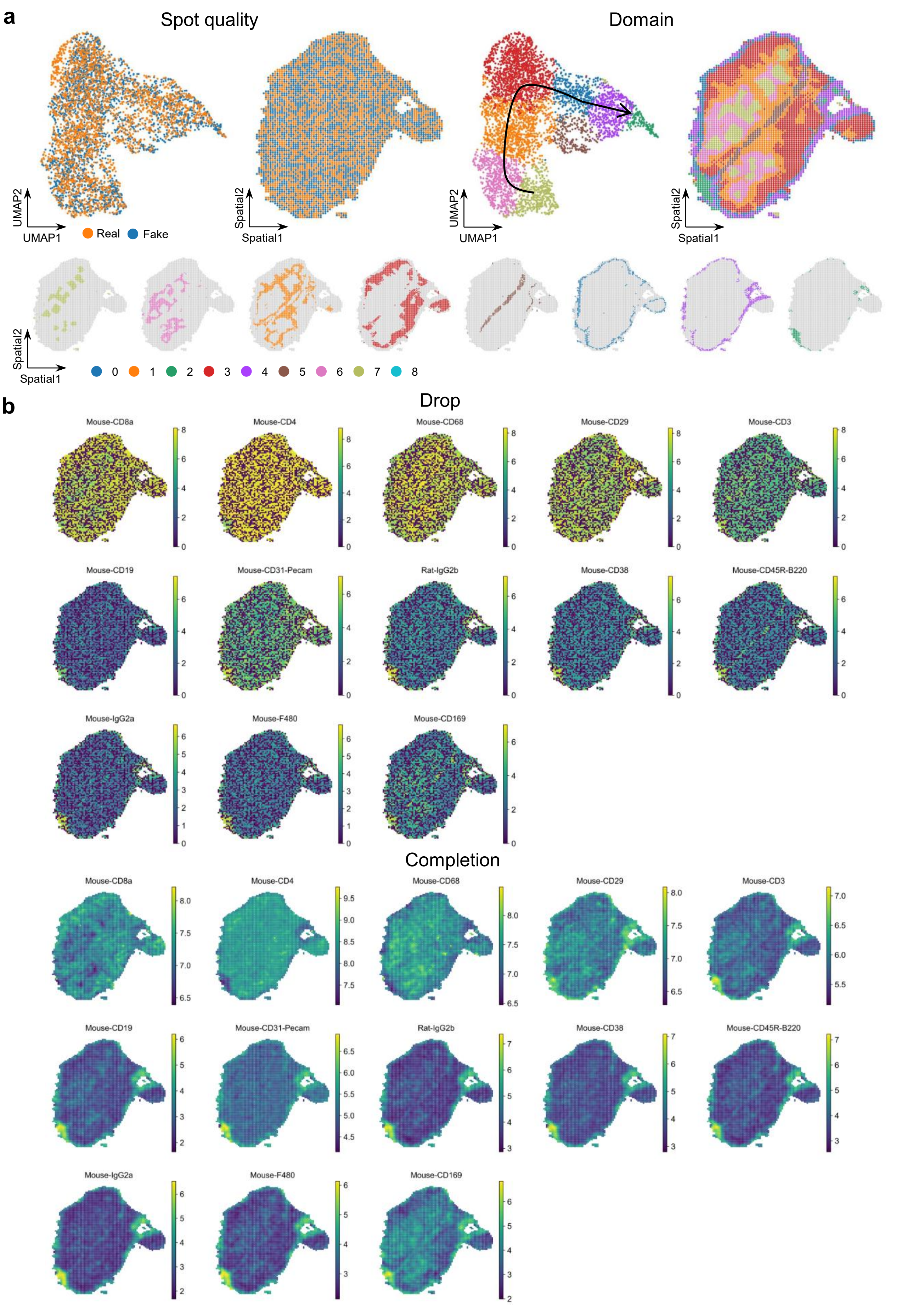


**Fig. S29.** Spatial spot completion by STAX on the proteomics dataset with being randomly discarded 50% of the spots. **a**, Spot completion and domain identification on the completion data. **b**, Spatial expression patterns of key proteins before and after completion.

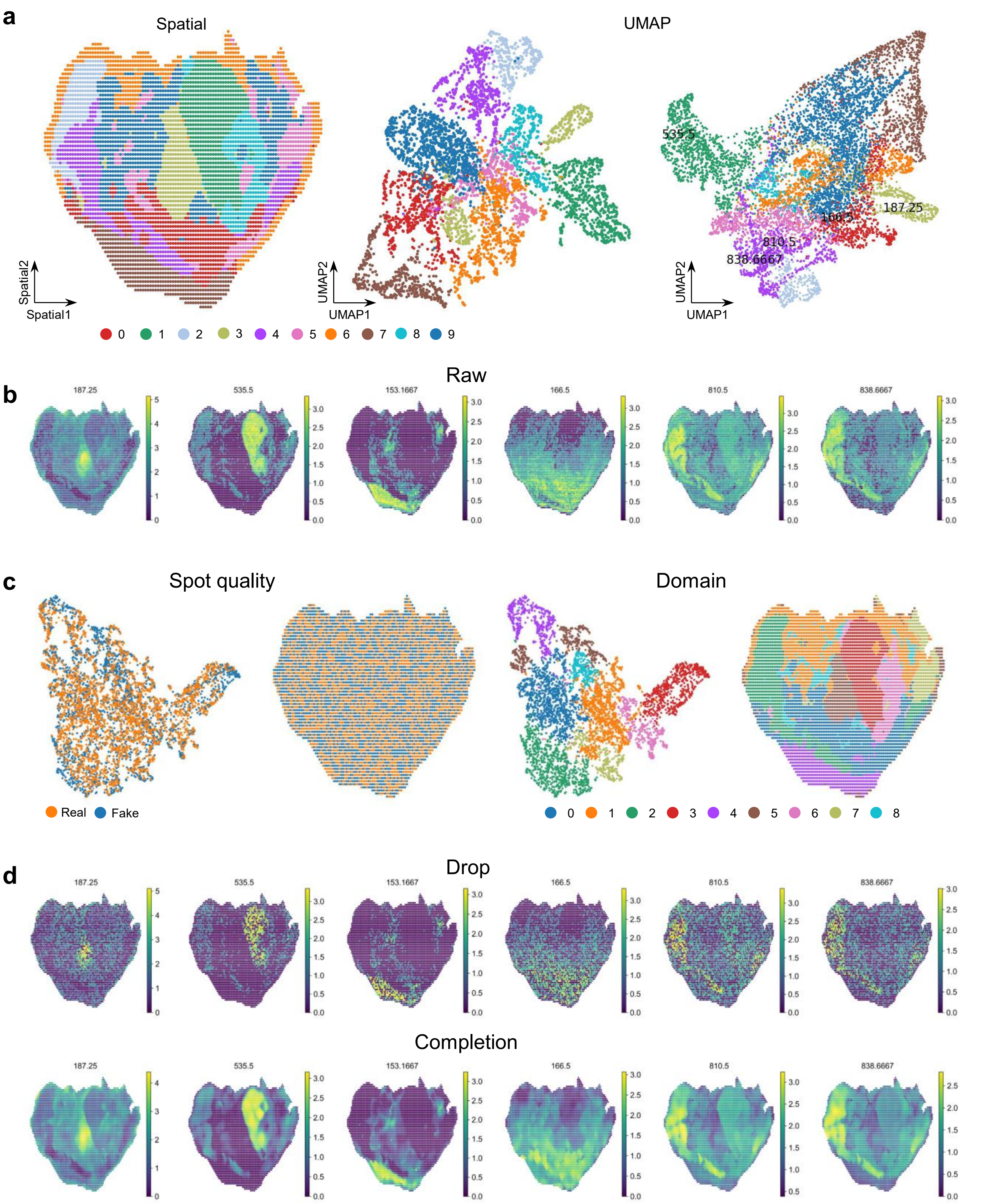


**Fig. S30.** Application to the pig embryo spatial metabolomics dataset by STAX. **a**, Spatial domain identification (left) and cell-metabolite co-embedding (right) of several marker peaks. **b**, Spatial expression patterns of several marker peaks in the dataset. **c**, Spot completion and domain identification based the completion data. **d**, Spatial expression patterns of marker peaks before and after completion.

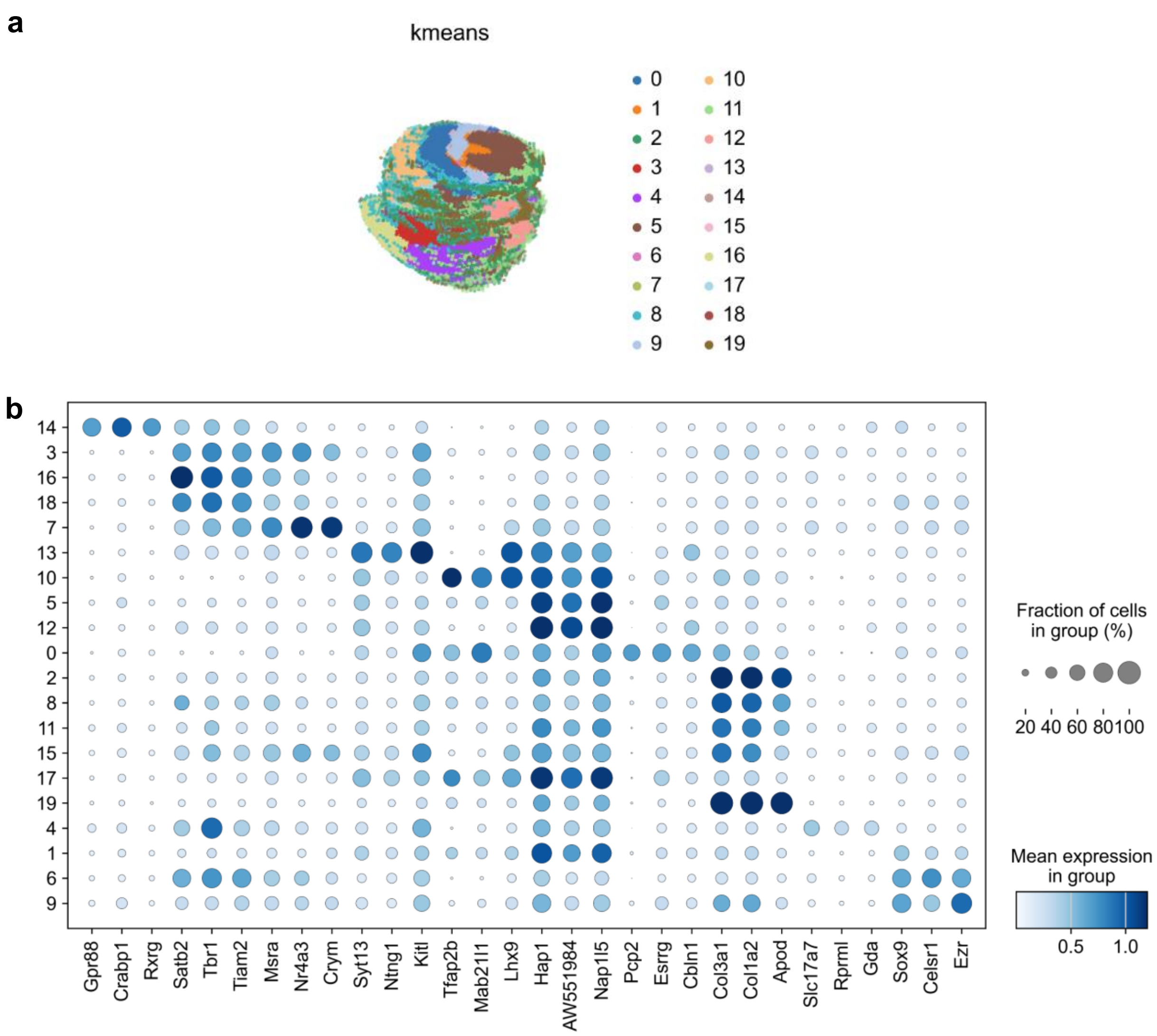


**Fig. S31.** Spatial domains identified by STAX on the Magic-seq dataset. **a**, 3D spatial plot of domains identified by STAX with kmeans=20. **b**, Dot plot of marker gene expression for 20 kmeans clusters.

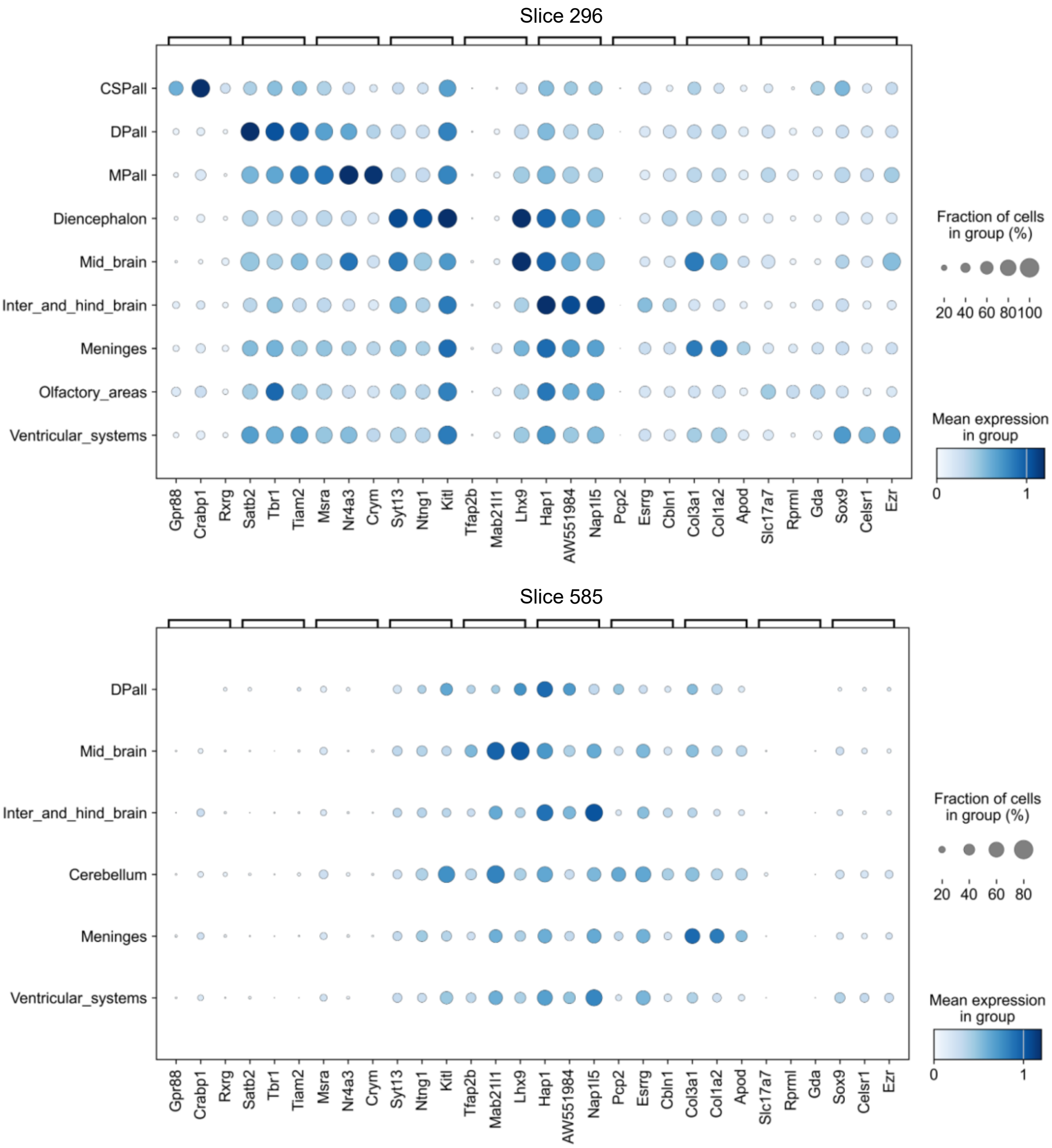


**Fig. S32.** Dot plot of marker gene expression in slices 295 and 585 for spatial domains identified by STAX on the Magic-seq dataset.

**a**

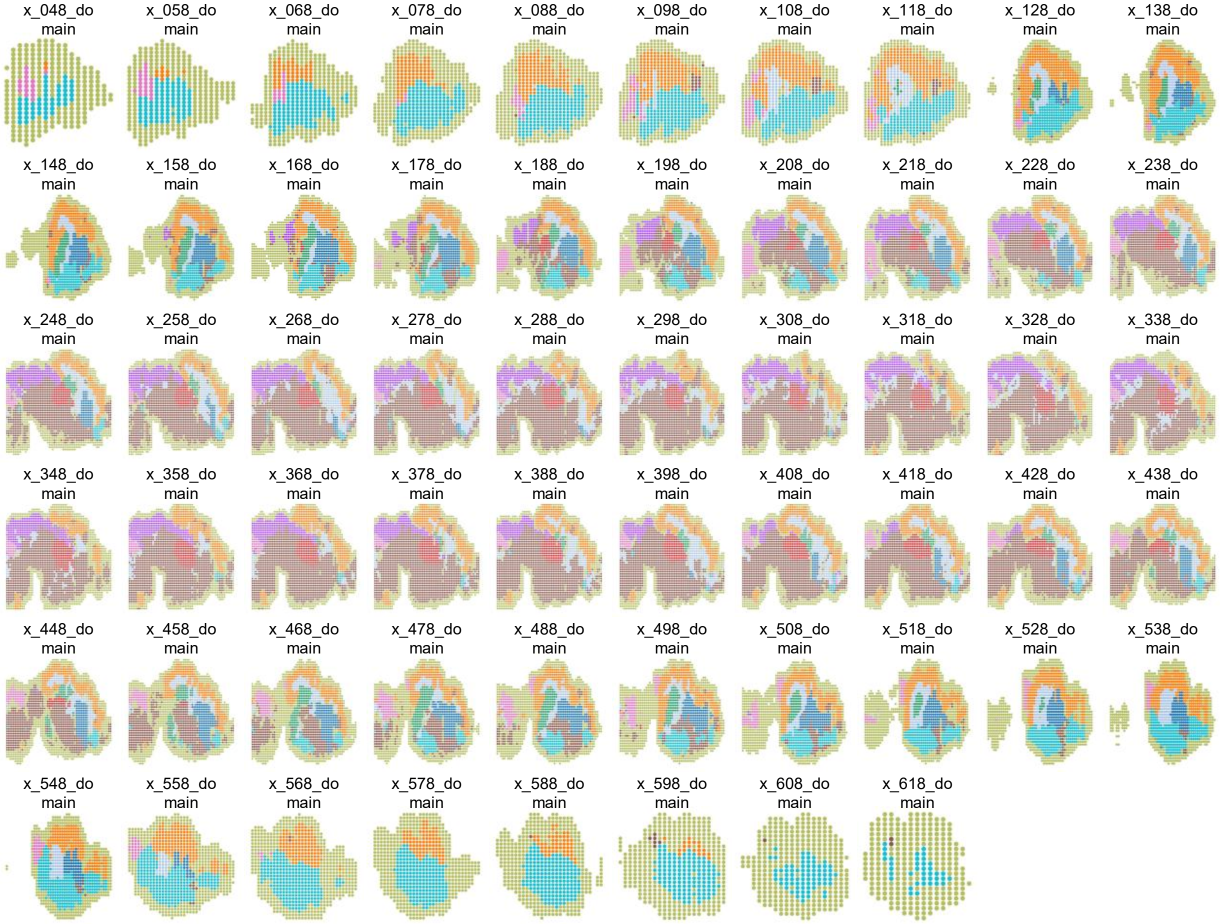


**Fig. S33.** Spatial domains identified by STAX on the simulation dataset with 90k spots (100μm) from the Sagittal view.

**a**

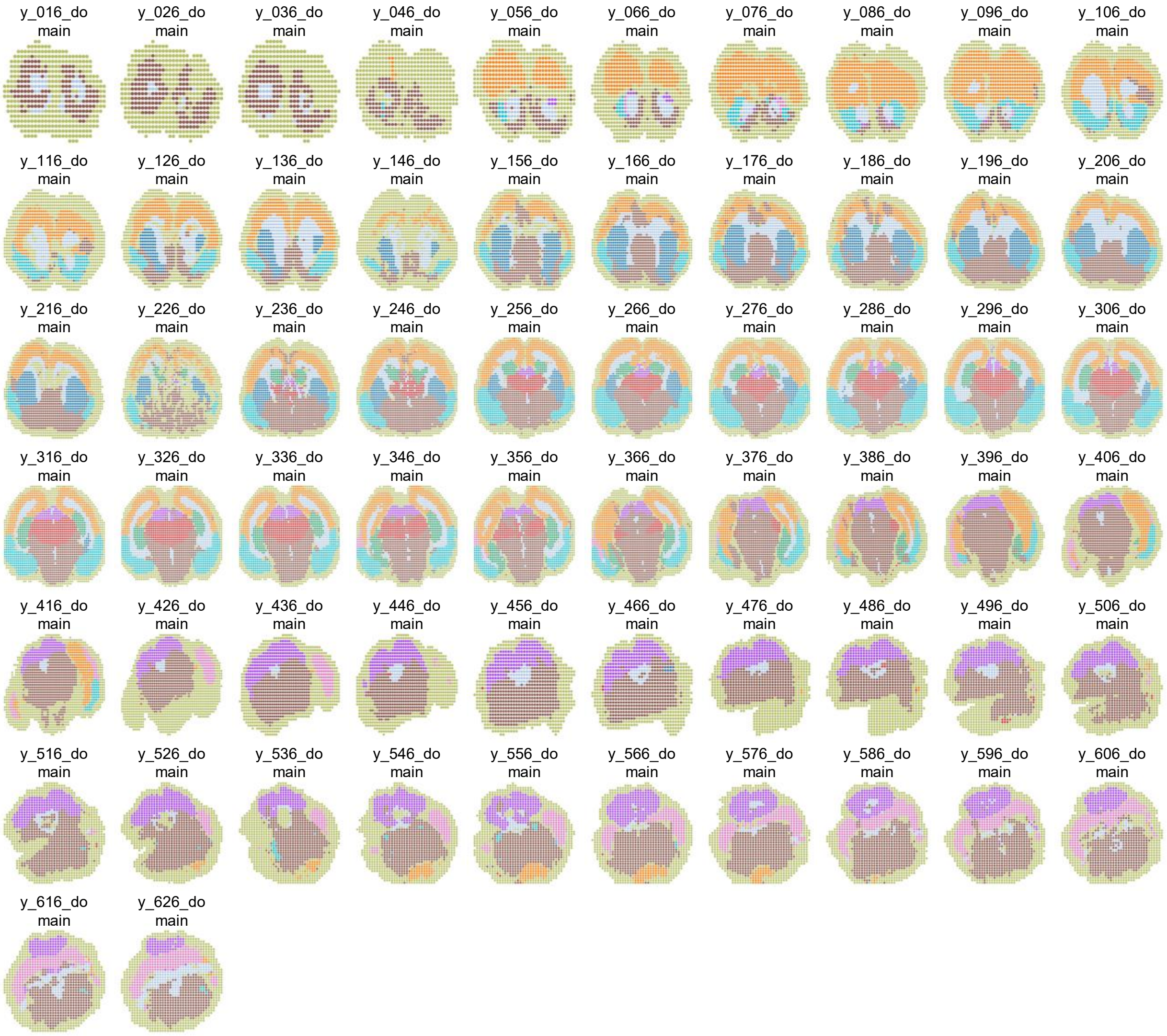


**Fig. S34.** Spatial domains identified by STAX on the simulation dataset with 90k spots (100μm) from the Coronal view.

a

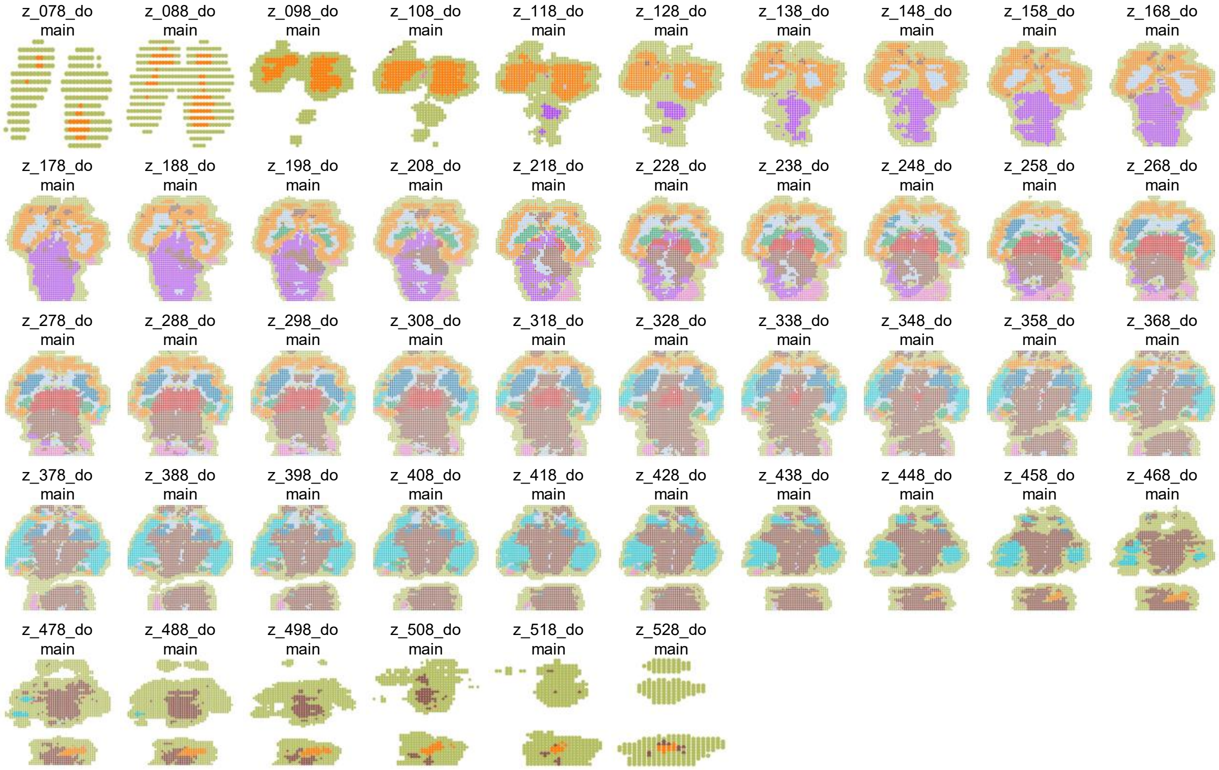


**Fig. S35.** Spatial domains identified by STAX on the simulation dataset with 90k spots (100μm) from the Transverse view.

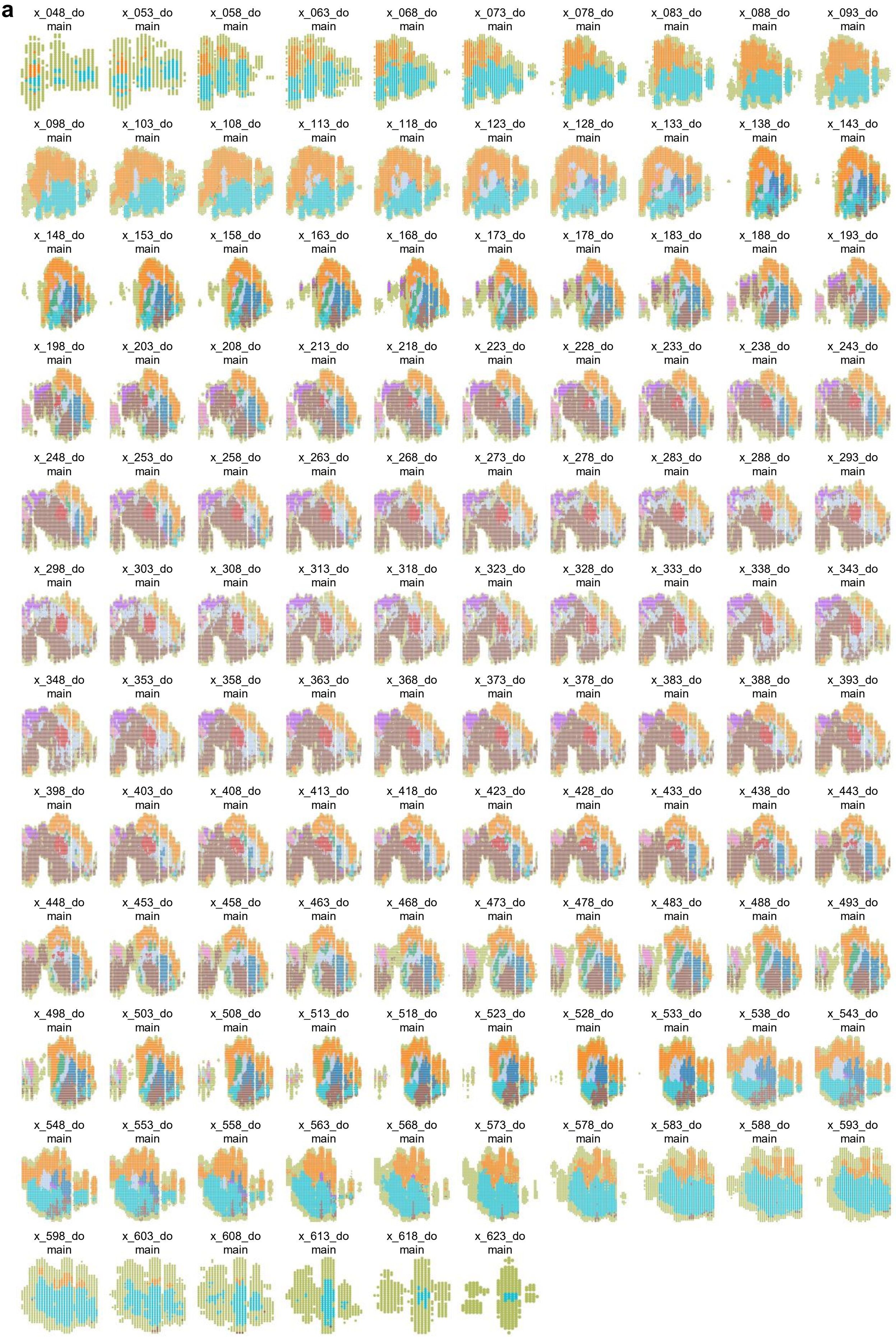


**Fig. S36.** Spatial domains identified by STAX on the simulation dataset with 570k spots (50μm) from the Sagittal view.

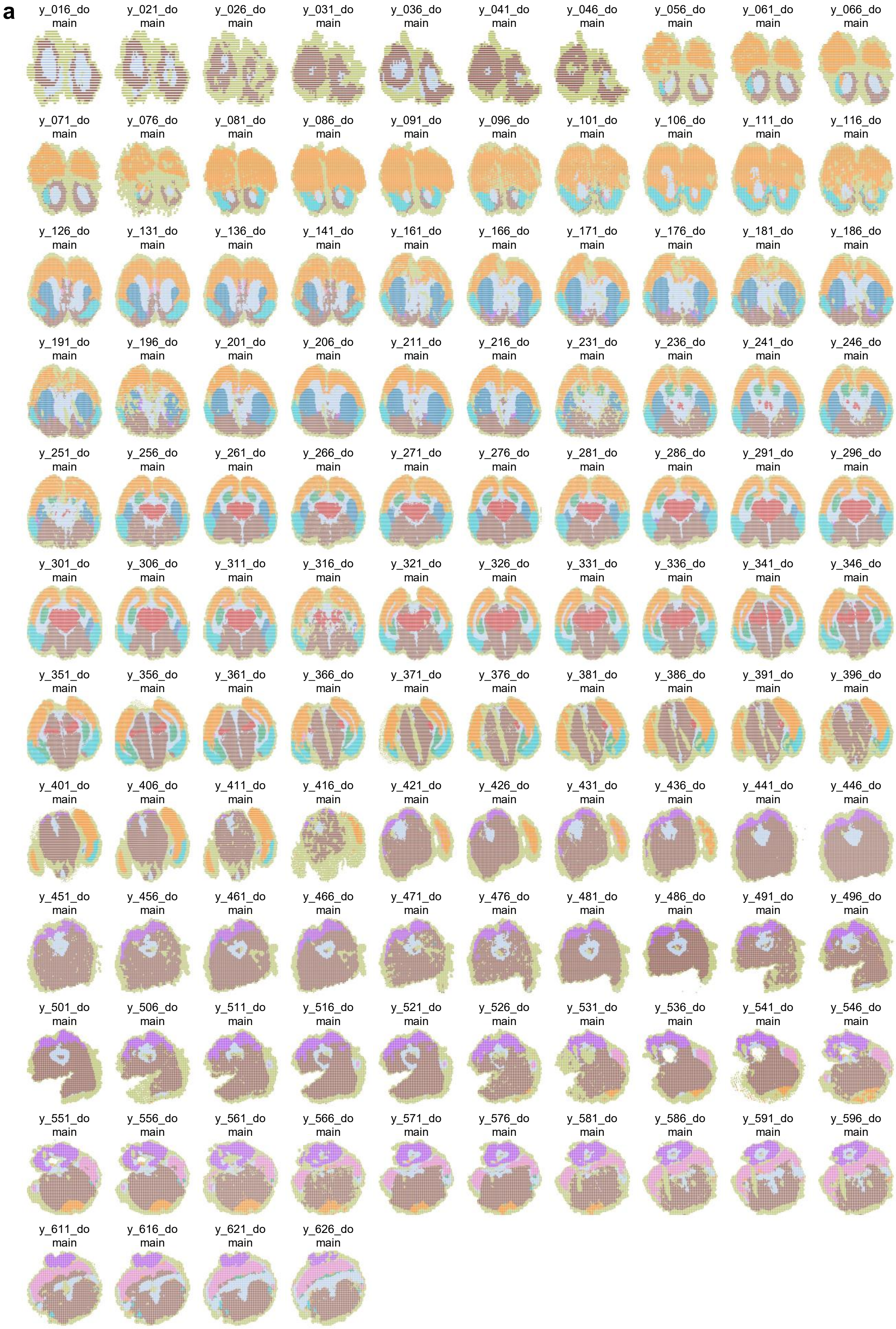


**Fig. S37.** Spatial domains identified by STAX on the simulation dataset with 570k spots (50µm) from the Coronal view.

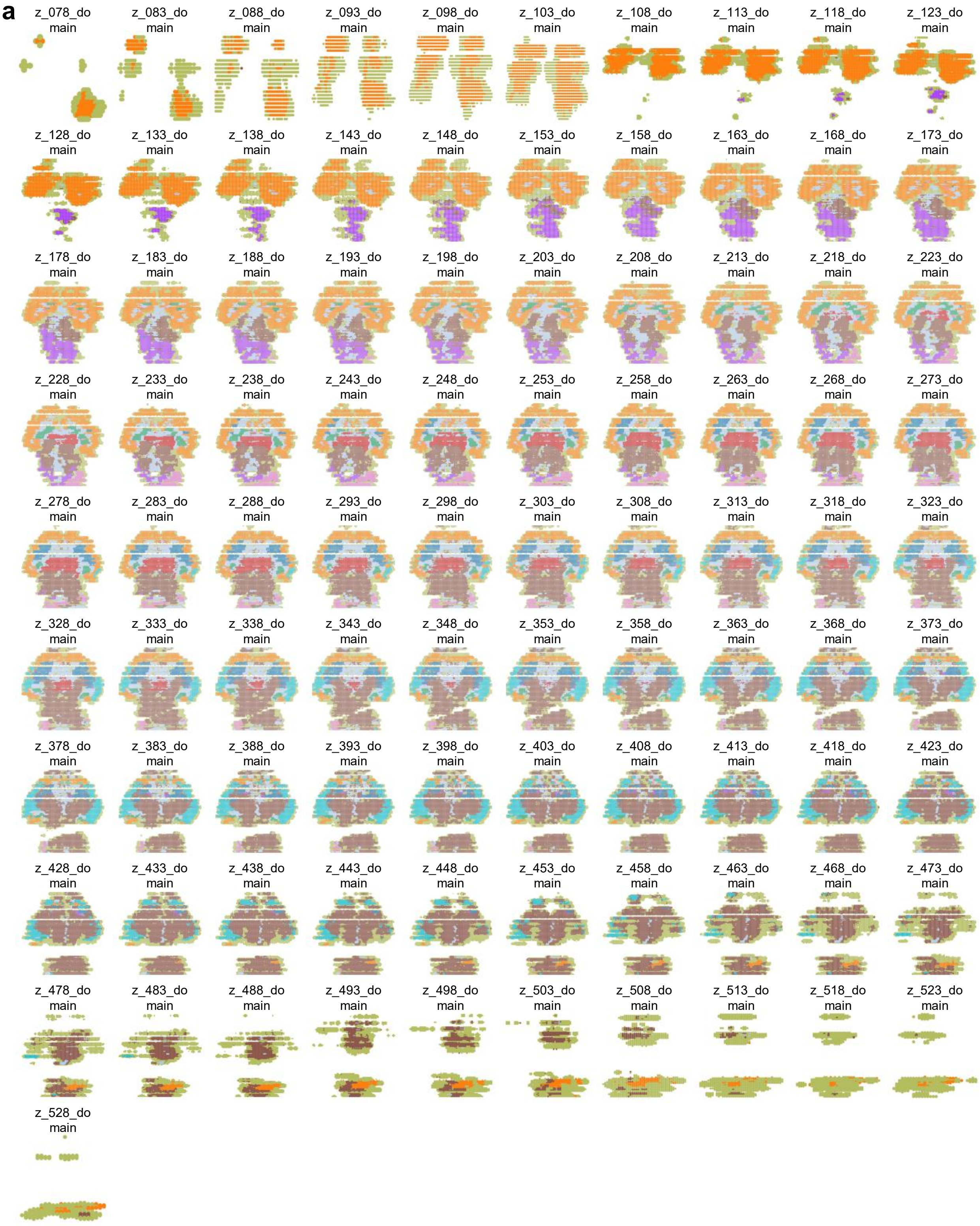


**Fig. S38.** Spatial domains identified by STAX on the simulation dataset with 570k spots (50μm) from the Transverse view.

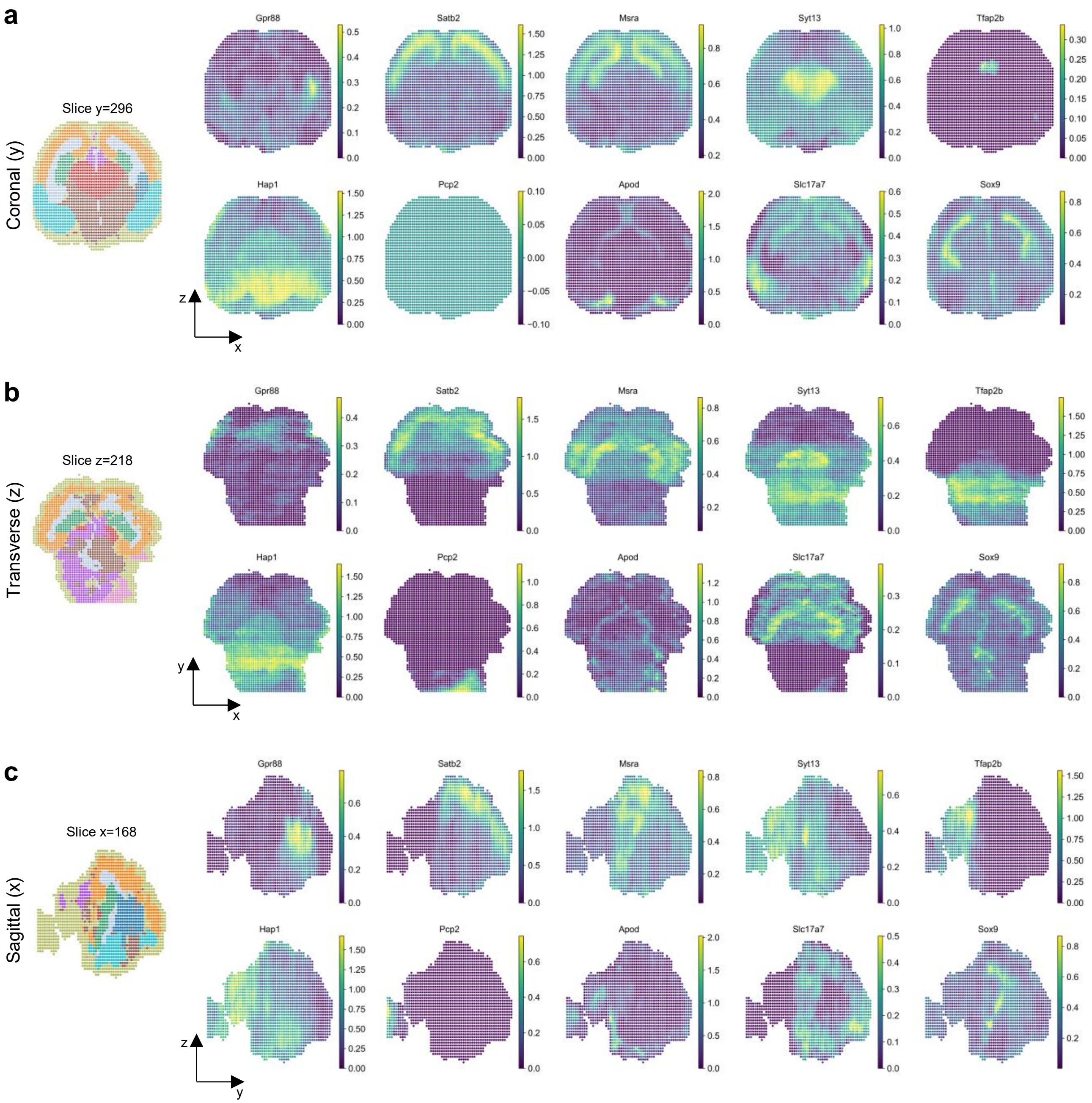


**Fig. S39.** Spatial domains identified by STAX on the simulation dataset with 90k spots (100μm). **a**, The domain and heat map of 10 domain marker genes expressed in slice y=296. **b**, The domain and heat map of 10 domain marker genes expressed in slice z=218. **c**, The domain and heat map of 10 domain marker genes expressed in slice x=168.

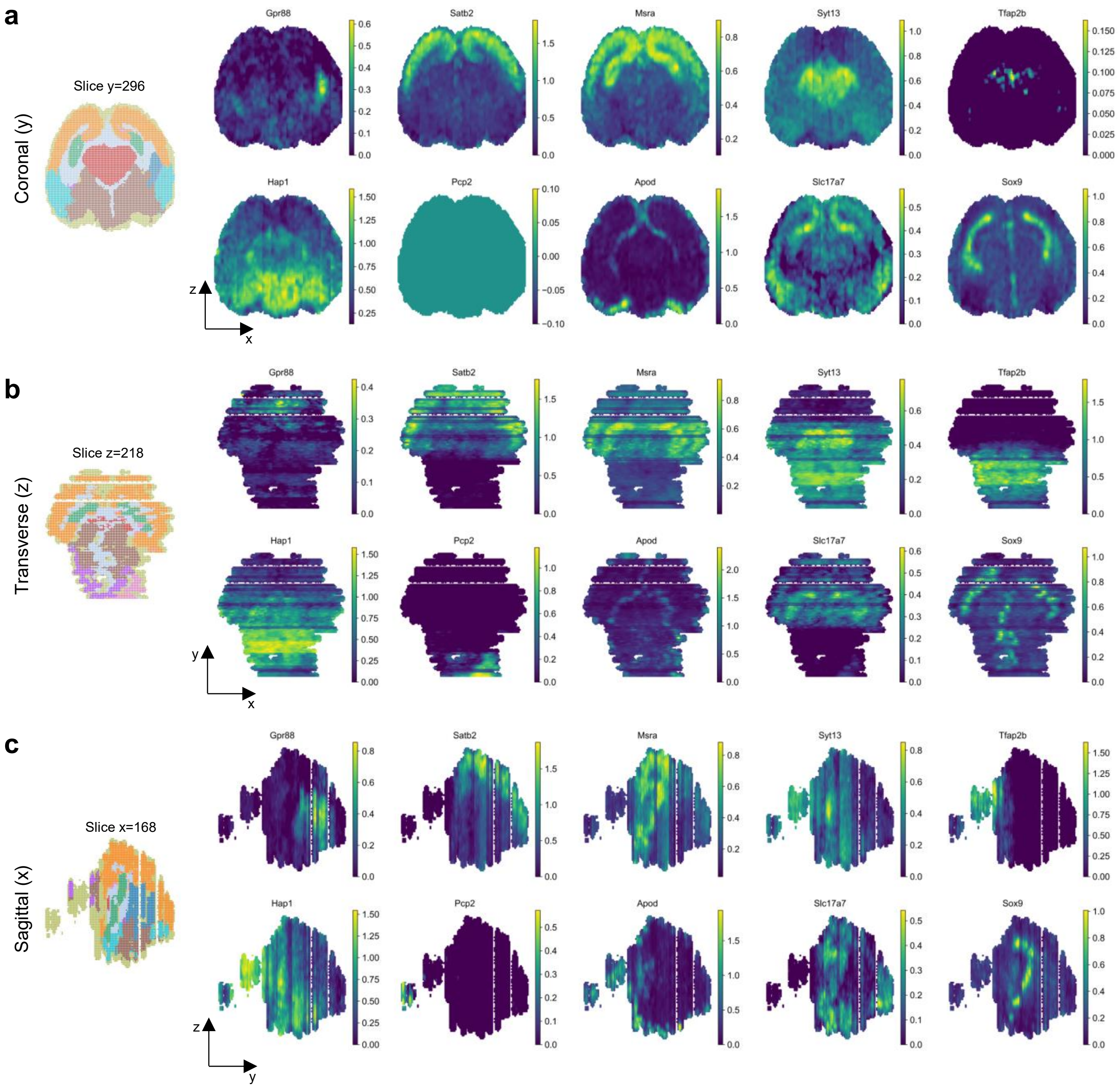


**Fig. S40.** Spatial domains identified by STAX on the simulation dataset with 570k spots (50μm). **a**, The domain and heat map of 10 domain marker genes expressed in slice y=296. **b**, The domain and heat map of 10 domain marker genes expressed in slice z=218. **c**, The domain and heat map of 10 domain marker genes expressed in slice x=168.
